# Resolving the Dynamic Interplay of the Conformational States in the TPR Domain of Human AMPylase FicD

**DOI:** 10.1101/2024.12.18.629078

**Authors:** Svenja Runge, Ecenaz Bilgen, Aymelt Itzen, Don C. Lamb

## Abstract

The human Fic-enzyme FicD plays an important role in regulating the Hsp70 homolog BiP in the endoplasmic reticulum (ER): FicD reversibly modulates BiP’s activity through attaching an adenosine monophosphate (AMP) to the substrate binding domain. This reduces BIP’s chaperone activity by shifting it into a conformation with reduced substrate affinity. Crystal structures of FicD in the apo, ATP-bound, and BiP-bound states revealed significant conformational variability in the tetratricopeptide repeat (TPR) motifs. In this study, we investigate the conformational dynamics of FicD’s TPR motifs using two-color single-molecule Förster resonance energy transfer (smFRET). We demonstrate that the TPR motifs exhibit conformational dynamics between the TPR-out and TPR-in conformations on timescales ranging from milliseconds to microseconds. In addition, we extend our investigation on multiple labeling positions within FicD, revealing how conformational dynamics vary depending on the location within the TPR motif. We quantify the motions with dynamic photon distribution analysis (PDA) for the FRET constructs and propose a conformational landscape model for FicD where the TPR-in/out states exist in an equilibrium that is altered due to the presence of ATP and BiP.

## Introduction

The heat shock protein 70 (Hsp70) family of proteins is essential for protein homeostasis from bacteria to humans. Despite their various functions which include assisting in protein folding, targeting and transportation of newly synthesized proteins, they share common structural and mechanistic features^1^. BiP is the Hsp70 homolog found in the endoplasmic reticulum (ER) of mammalian cells where it participates in translocation of proteins across the ER, targets misfolded proteins for degradation and aids in the folding of nascent proteins^2^. Like all Hsp70 proteins, BiP shows an adenosine-triphosphate (ATP) dependent conformational cycle that regulates its chaperoning activity: In the ATP-bound state of the nucleotide binding domain (NBD), the substrate binding domain (SBD) loosely associates with unfolded client proteins^3^. However, after ATP-hydrolysis to ADP, the SBD gains high affinity to the client, thereby stabilizing it until subsequent replacement of ADP with ATP releases the client. This nucleotide-dependent cycle is accompanied with structural changes in the Hsp70: In the ATP-state, the SBD docks to the NBD whereas, in the ADP-bound form, the SBD dislodges from the NBD^4–6^.

Being involved in ER homeostasis, BiP is highly susceptible to changes in the protein folding load and contributes to regulation of the unfolded protein response (UPR)^7^. BiP’s ability to bind client proteins is regulated by the ER-resident “filamentation induced by cyclic-AMP’’ (Fic) domain containing protein FicD^8,9^. FicD is the only representative of the Fic-enzyme family within the human genome and is highly conserved among eukaryotic species^10^. Using ATP as a co-substrate, the bi-functional enzyme can reversibly transfer an adenosine-monophosphate (AMP) to the hydroxyl group of residue T518 in the SBD of BiP, a process termed AMPylation^11–14^. AMPylation traps BiP in a conformation where the NBD and SBD are docked, establishing a pool of ATP hydrolysis deficient BiP with low client-binding affinity^15^. While AMPylation-mediated BiP inactivation occurs during low client protein load, the deAMPylation of BiP by FicD re-activates BiP during protein-folding stress^16^. This posttranslational regulation of BiP has been proven to be essential in stress resilience of cells and tissues^17^ and mutations in FicD were linked to motoneuron and pancreatic diseases^18,19^.

The N-terminus of FicD (amino acid residues 1-105) contains a single transmembrane domain and is structurally uncharacterized. In contrast, the ER luminal domains are well studied: two tetratricopeptide repeat (TPR) motifs, each composed of two α-helices, are connected to the Fic domain by a helical linker^20,21^ (Fig. 1A). The Fic domain contains the conserved sequence motifs of this protein family, i.e. the catalytic H_cat_PFx(D/E)GN(G/K)R motif and the inhibitory (S/T)XXXE_inh_(G/N) motif^22,23^. Mutation of the catalytic H363 (H_cat_) to alanine abrogates the enzymatic activity of Fic enzymes. The inhibitory E234 (E_inh_) instead inhibits AMPylation and is required for deAMPylation; mutation of E234 to glycine results in enhanced AMPylation activity of FicD^22^. In the ER, functional regulation is conferred by the dimerization state of FicD: The dimer interface between the Fic domains is connected to the catalytic center via a hydrogen bond network, favoring deAMPylation by imposing rigidity to E_inh_. In monomerized FicD (e.g. by the L258D mutation), a higher flexibility within this network allows deflection of the repressive residue and leads to the AMPylation competent binding of Mg:ATP. In addition, the binding of ATP was proposed to allosterically regulate FicD’s dimerization equilibrium and affinity to BiP^21^. The TPR motifs of FicD are essential for target recognition and binding during AMPylation and deAMPylation reactions^24^. Moreover, an intrinsic flexibility of this domain is linked to both catalytic activities of FicD, as reaction rates are significantly reduced upon fixation to the interdomain linker helix via an engineered disulfide bond^25^.

**Fig. 1:**
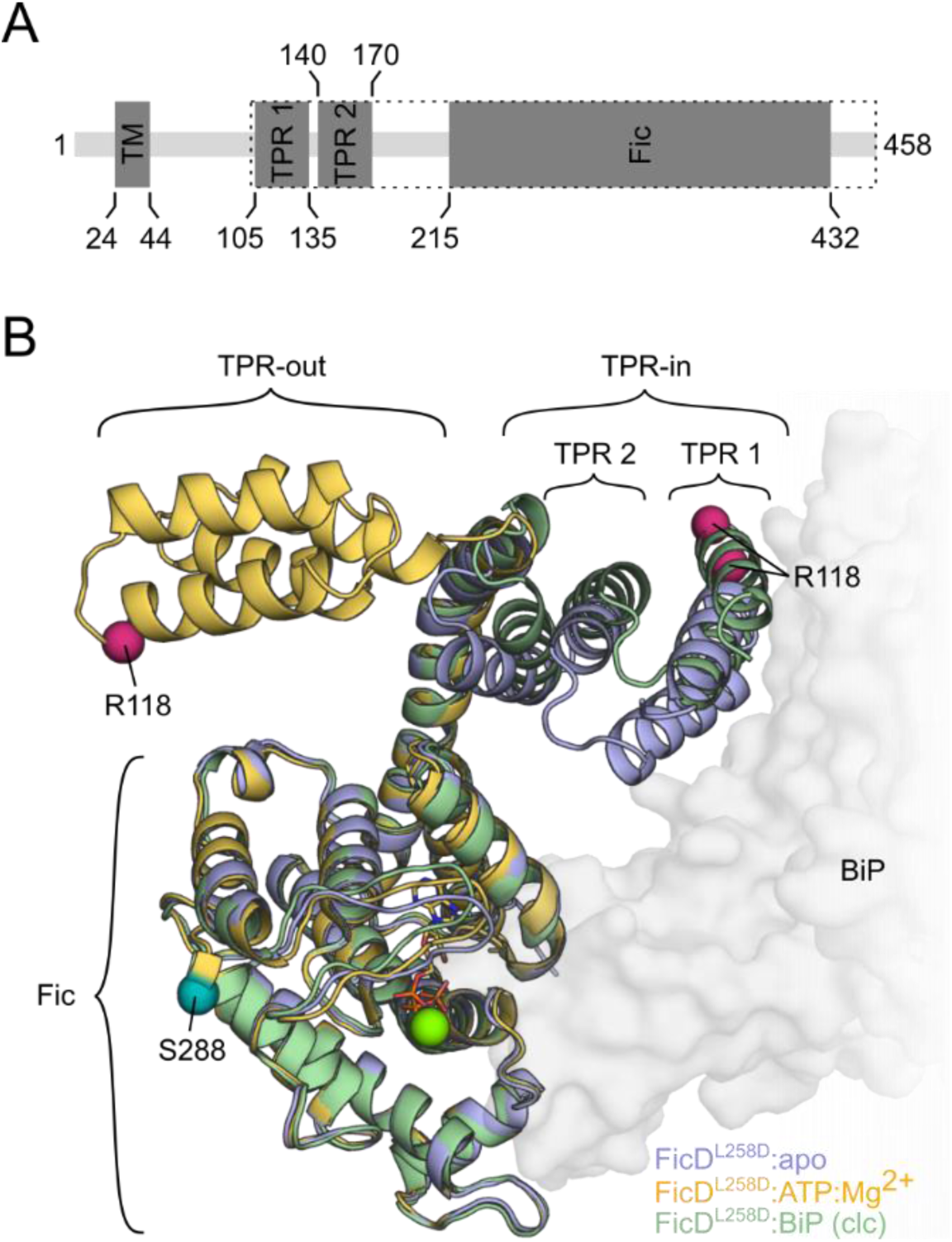
The structure and conformations of FicD. **A:** Domain organization of FicD. The construct used in this study is highlighted by the dotted frame (amino acids 102-458). TM: transmembrane-domain, TPR: tetratricopeptide repeat motif, Fic: filamentation induced by cyclic-AMP domain. **B:** Superimposed representations of the monomeric FicD^L258D^ crystal structures in the apo-state (light blue, PDB 6I7J^21^), bound to ATP (yellow, PDB 6I7K^21^, stick representation: ATP, green sphere: Mg-ion) and in a covalently linked complex (clc) with BiP (light green, PDB 6ZMD^24^, surface representation: BiP). The smFRET labeling positions R118 (magenta) and S288 (teal) are shown as spheres on the C𝛼-atoms of the amino acid residues.

A comparison of the crystal structures of monomeric nucleotide-free (apo-state) versus ATP-bound FicD^L258D^ hints at a potential rearrangement between the Fic and TPR domain in the nucleotide-bound state^21^. In the apo-state, the TPR domain crystallized in a commonly observed TPR-in conformation (Fig. 1B, light blue) that is structurally similar to its conformation in the dimeric enzyme when bound to BiP (Fig. 1B, green)^20,21,24,25^. In contrast, the TPR domain of the ATP-bound FicD^L258D^ crystallized in a TPR-out conformation, being flipped outwards by ca. 180°^21^ (Fig. 1B, yellow). Although the crystal structures are informative, details of FicD dynamics and of its structural transitions in solution are missing. Therefore, we applied single-molecule Förster Resonance Energy Transfer (smFRET) spectroscopy to probe the interdomain rearrangements of FicD in dependence of its interactions with BiP and the co-substrate ATP. Our experiments reveal that both the TPR-in and TRP-out conformations coexist in equilibrium in-solution with transitions occurring on the millisecond to microsecond timescales. We demonstrate that ATP and BiP modulate the transitions of the TPR domain. Furthermore, the conformational dynamics of FicD are conserved in both the monomeric and dimeric states, but are altered upon its interaction with BiP and is dependent on the AMPylation state of BiP.

## Results

### The conformation of the TPR-domain in FicD is dynamic

To investigate the conformational flexibility of the TPR-motifs relative to the Fic domain in solution, we fluorescently labeled monomeric FicD^L258D^ (Fig. 1B) for smFRET experiments. The monomeric mutant was chosen since domain rearrangements were observed for monomeric FicD crystals. Cysteine substitutions were implemented for labeling of the protein with maleimide-functionalized fluorophores ATTO532 as the donor dye (D) and ATTO643 as the acceptor dye (A). We chose S288C as a labeling position on the surface of the Fic domain, which has been previously implemented for fluorescent labeling^21^, and combined it with R118C as a second position located on the 1^st^ TPR motif. The integrity of these constructs was validated via thermal stability analysis using nano differential scanning fluorimetry (nDSF) (Fig. S1A). After fluorescent labeling of FicD^L258D^ ^S288C^ ^R118C^ (hereafter referred to as FicD^R118C^), the presence of hetero, double-labeled species was confirmed via intact mass spectrometry (MS) (Fig. S1B, C). Note that the endogenous cysteine (C421) of FicD was not removed by mutation in our smFRET samples as FicD^C421S^ exhibits a substantially reduced thermal stability (Fig. S1A), and only a negligible amount (≤ 5%) of triple labeled species were detected by MS (Fig. S1C). Since the TPR domain is essential for the interaction with BiP^24,25^, BIP-AMPylation assays were used to ensure that the mutations and fluorescent dyes do not affect FicDs activity (Fig. S1D,E).

For the dual-color smFRET experiments, we used multiparameter fluorescence detection (MFD) with pulsed interleaved excitation (PIE)^26^. This technique measures the FRET efficiency of single, freely diffusing FicD^R118C^ in solution as they transverse the observation volume. By gathering statistics of single molecule events, we can resolve the various conformational states of FicD with respect to the relative orientation of the TPR motifs to the Fic domain. Surprisingly, the FRET efficiency histogram for FicD^R118C^ in the apo-state reveals two distance populations with mean FRET efficiency values of E=0.31 (low-FRET) and E=0.71 (high-FRET) (Fig. 2A, light blue, Table S1). From the FRET efficiency (E) and the dye-pair’s specific Förster distance (R_0_=59 Å for the ATTO532-ATTO643 dye pair), the inter-dye distances in solution can be calculated using the formula 𝐸 = 1⁄(1 + (𝑅⁄𝑅_0_)^6^). For the apo-state of FicD^R118C^, the FRET efficiencies translate to distances of 67 Å and 49 Å for the low and high-FRET populations, respectively. Upon the addition of ATP, two highly similar states with FRET efficiencies of E=0.33 (R=66 Å) and E=0.73 (R=50 Å) are observed. However, the contribution of the high-FRET state is higher compared to the low-FRET state in the presence of ATP (Fig. 2A, yellow, Table S1).

**Fig. 2:**
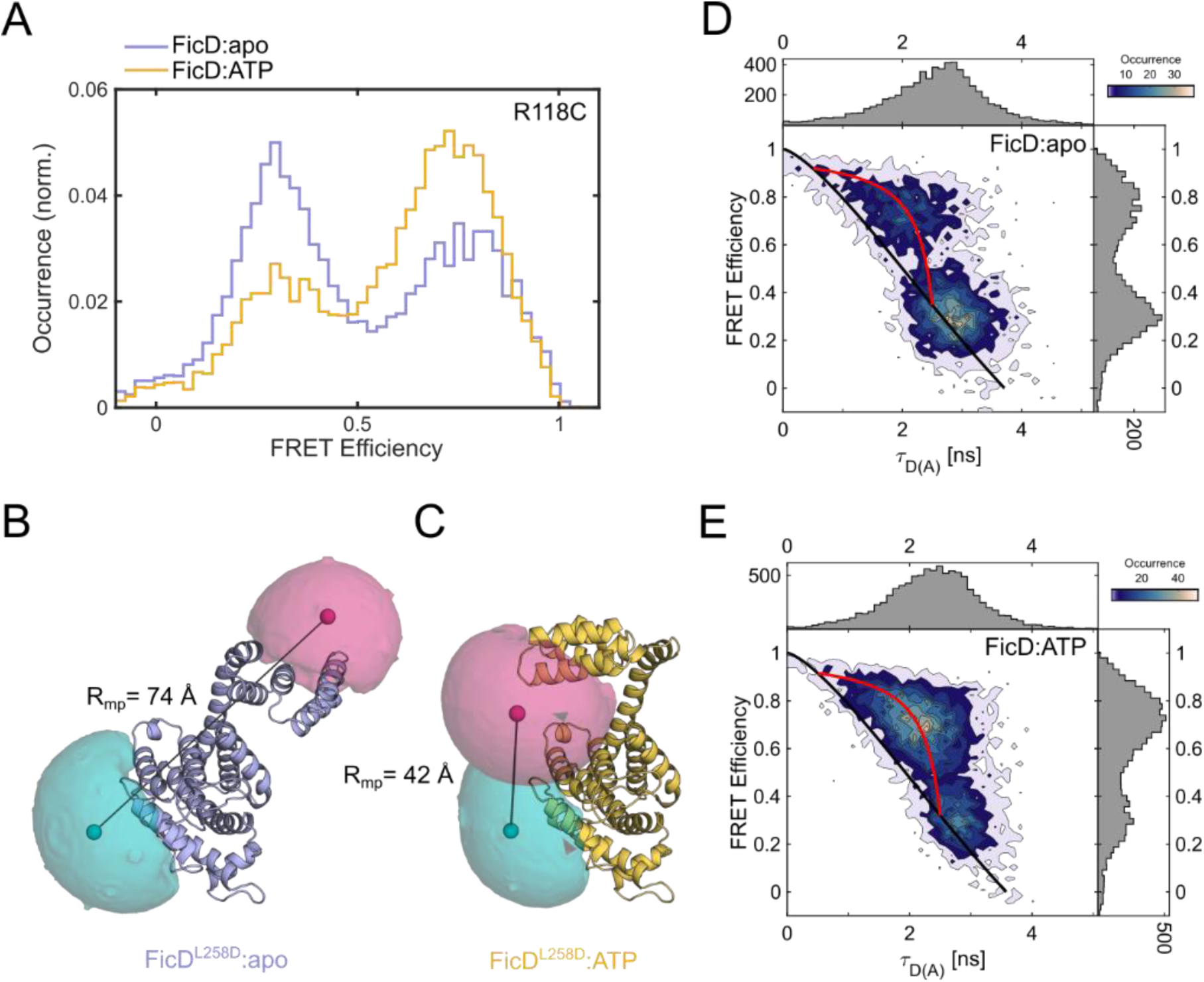
The conformational equilibrium of FicD in solution. **A**: SmFRET efficiency histograms of FicD^R118C^ alone (light blue) and in the presence of ATP (yellow). **B-C:** Accessible volume (AV) calculations of **B**: FicD in the apo-state (light blue, PDB 6I7J) and **C**: in the ATP-bound state (yellow, PDB 6I7K). The magenta (donor) and teal (acceptor) surfaces represent the possible spatial distribution of the fluorophores whereas the spheres indicate the mean positions of the donor and acceptor dyes. Black lines indicate the distances between both mean positions (R_mp_). **D-E**: 2D plots of donor fluorescence lifetime in the presence of an acceptor (𝜏_𝐷(𝐴)_) vs FRET efficiency (E) for **D:** FicD^R118C^:apo and **E:** FicD^R118C^ in the presence of ATP. Black and red lines represent the static and dynamic FRET lines, respectively.

To check the correlation between the experimentally detected states and the aforementioned crystal structures^21^, we performed accessible volume (AV) calculations^27^ based on the apo (PDB 6I7J^21^) and ATP-bound (PDB 6I7K^21^) FicD crystal structures. The theoretically accessible positions of the fluorescent dyes (ATTO532 and ATTO643) attached at positions R118C and S288C (surface clouds in Fig. 2B,C, Table S2) allow us to predict the mean distance between the dye pair (R_mp_). The R_mp_ of 74 Å for the apo-state of FicD roughly corresponds to the measured low-FRET conformation of FicD^R118C^ (R=66-67 Å). For the ATP-bound conformation of FicD, the AV calculations predicted a R_mp_ of 42 Å, which is similar to the distance measured for the high-FRET conformation of FicD^R118C^ (R=49-50 Å). As this conformation was also enhanced in the presence of ATP (Fig. 2A, yellow), we attribute the high-FRET population with a FRET efficiency value of ca. 0.72 to a TPR-out conformation as represented by the FicD:ATP crystal structure (Fig. 1C, yellow). Similarly, the low-FRET population with an efficiency of ca. 0.32 (Fig. 2A) likely corresponds to a TPR-in conformation as represented by the apo-state FicD crystal structure (Fig. 1B, light blue). From the smFRET experiments we can clearly conclude that the TPR-out and TPR-in conformations of FicD exist in an equilibrium in solution, and that the addition of ATP shifts this conformational equilibrium towards the TPR-out conformation.

### FicD interconverts between TPR-in and TPR-out conformations

To evaluate whether the TPR-in and TPR-out conformations of FicD interconvert dynamically on the millisecond timescale, we made use of the fluorescence lifetime information available from the MFD-PIE measurements^26^. In a two-dimensional plot of the fluorescence lifetime of the donor dye in the presence of acceptor (𝜏_𝐷(𝐴)_) vs the intensity-based FRET efficiency (E) (E-𝜏_𝐷(𝐴)_ plot), the static FRET line (black line in Fig. 2D, E; Eq. 2) represents the theoretical E-𝜏_𝐷(𝐴)_correlation for static molecules. Populations that undergo conformational transitions during their diffusion through the observation volume appear shifted to the right from the static FRET line and can be analyzed using a dynamic FRET line (red line in Fig. 2D, E; Eq. 3)^28,29^. For systems that show fluctuations on the µs to ms timescale between two conformational states, fitting the fluorescence lifetime distribution of 𝜏_𝐷(𝐴)_ to a biexponential function determines the two endpoints for the dynamic FRET line.

In the E-𝜏_𝐷(𝐴)_ plot for FicD^R118C^ in the absence and presence of ATP (Fig. 2D, E), the low-FRET population at E=∼0.32 lies on the static FRET line (black) of the E-𝜏_𝐷(𝐴)_ plot, indicating that this population represents a TPR-in conformation that is static on the ms timescale. The corresponding lifetime of 𝜏_𝐷(𝐴)_ = 2.8 𝑛𝑠 translates into in a fluorophore distance of 72 Å (Table S3). In contrast, the high-FRET populations at E=0.72 reveal a substantial shift to the right of the static FRET line (Fig. 2D, E). The lifetime analysis of the dynamic high-FRET population of FicD^R118C^ in the absence of ATP revealed endpoints of the dynamic FRET line (red) at fluorescence lifetimes of 𝜏_𝐷(𝐴)_ = 2.4 𝑛𝑠 and 𝜏_𝐷(𝐴)_ = 0.5 𝑛𝑠. This corresponds to FRET efficiencies of 0.35 and 0.87 and fluorophore distances of 65 Å and 43 Å, respectively (Fig. 2D, Table S3). This indicates a dynamic flipping of the TPR domain on the µs to ms timescale between a TPR-in (65 Å) and a TPR-out (43 Å) state. Notably, these dynamic endpoints correlate even more accurately with the AV calculations (Table S2) as the high-FRET state in the 1D smFRET histogram (Fig. 2A) is a dynamically averaged state due to flipping of the TPR domain. The 2D E-𝜏_𝐷(𝐴)_plot in the presence of ATP also indicates dynamics with an increased fraction of the dynamically interconverting FicD^R118C^ population between the TPR-in and -out states (Fig. 2E). In conclusion, the lifetime analysis of FicD provides evidence that the observed high-FRET population is the result of dynamic interconversion between a rather stable TPR-in conformation and a transient TPR-out conformation. Thus, three distinguishable conformational states are observed: the static, low-FRET TPR-in conformation (state 1) versus the dynamically interconverting low-FRET TPR-in (state 2) and high-FRET TPR-out (state 3) conformations.

### BiP affects the conformational distribution of FicD

#### Non-AMPylated BiP

Next, we investigated the influence of BiP on the conformational equilibrium of FicD. These smFRET experiments were performed using near full-length (amino acids 19-654), ATP-hydrolysis deficient BiP^T229A^ ^5,30^. In addition to the reduction of BiP-dependent ATP hydrolysis, this mutant is a more efficient AMPylation substrate for FicD^12^. The affinity for the interaction between BiP and FicD is approximately 2-20 µM^16,25,31^. Hence, we used 100 µM of unlabeled BiP (5-50-fold K_D_/K_M_) to saturate the enzyme-substrate complex. In comparison to the apo- and ATP-bound states, the FRET efficiency histograms of FicD^R118C^ in the presence of BiP show shifts in the absence and presence of ATP for both the static and the dynamically averaged FRET populations to lower FRET values of 0.24 and 0.61, respectively. (Fig. 3A, light blue, Table S2). Thus, the distance for the static TPR-in conformation of FicD:BiP increases to 72 Å, compared to ca. 67 Å in the absence of BiP (Fig. 2D, E). This is consistent with our AV calculations based on the crystal structure of a covalently linked complex (clc) between FicD and BiP (PDB 6ZMD)^24^, which predict a shift to a larger dye distance in the presence of BiP (R=78 Å) compared to the apo-state (R=74 Å) (Fig. S2, Table S3). The addition of ATP to FicD^R118C^ and BiP further shifted the static, low-FRET population to lower values (E=0.19, R=75 Å) (Fig. 3D, orange).

**Fig. 3:**
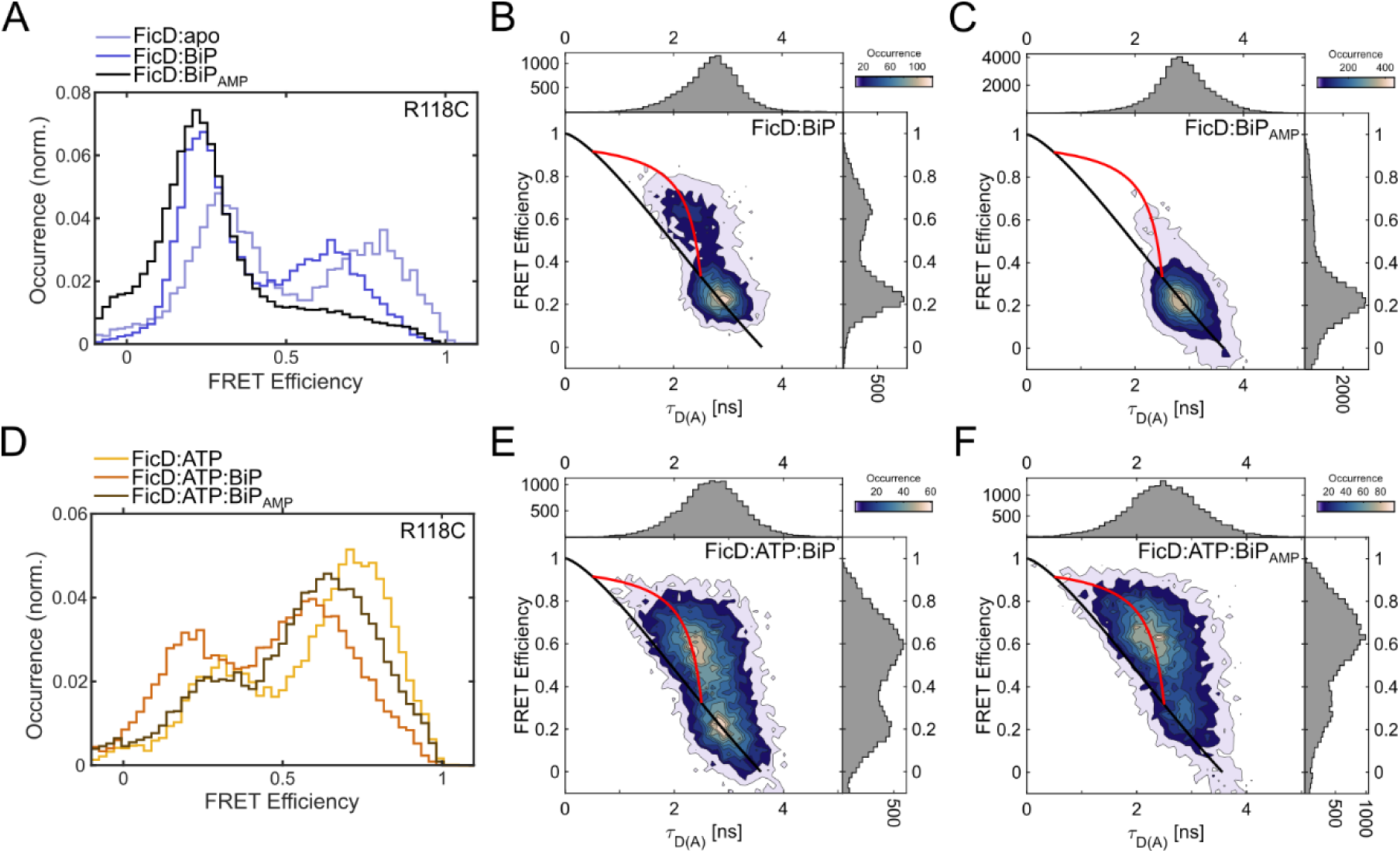
Binding to BiP alters the conformational equilibrium of FicD^R118C^. **A:** SmFRET efficiency histograms of FicD:apo (light blue) in comparison to FicD in the presence of BiP (blue) and BiP_AMP_ (black). **B-C:** 2D plots of donor fluorescence lifetime in the presence of an acceptor (𝜏_𝐷(𝐴)_) vs FRET efficiency (E) for **B:** FicD:BiP and **C:** FicD:BiP_AMP_. Black and red lines represent the static and dynamic FRET lines, respectively. **D**: SmFRET efficiency histograms of FicD:ATP (yellow) in comparison to FicD:ATP in the presence of BiP (orange) and BiP_AMP_ (brown). **E-F:** 2D plots of donor fluorescence lifetime in the presence of an acceptor (𝜏_𝐷(𝐴)_) vs FRET efficiency (E) for **E:** FicD:ATP:BiP and **C:** FicD:ATP:BiP_AMP_. Black and red lines represent the static and dynamic FRET lines, respectively.

The 2D E-𝜏_𝐷(𝐴)_ plot of FicD:BiP (Fig. 3B) confirms the dynamic averaging of the high-FRET population due to fluctuations between the dynamic TPR-in (E=0.29, R=68 Å) and TPR-out (E=0.86, R=43 Å) conformations. Moreover, the FRET efficiency histograms and E-𝜏_𝐷(𝐴)_plots indicate that the contribution of the dynamically interconverting high-FRET population increases upon addition of ATP to FicD and BiP (Fig. 3E, Table S1). To confirm the binding of FicD^R118C^ to BiP under the applied conditions, we performed a fluorescence correlation spectroscopy (FCS) analysis of the smFRET measurements and calculated the average diffusion coefficients (D) for the labeled protein (Fig. S3, Table S4). Due to the inverse correlation with the hydrodynamic radius, binding events between FicD^R118C^ and BiP can be expected to reduce the diffusion coefficient. Compared to apo-state FicD, the addition of ATP caused a minor (ca 5 %) decrease in the diffusion coefficient, whereas the addition of BiP and BiP with ATP resulted in a reduction by almost 20 % and 35 %, respectively. Thus, there is binding between FicD and BiP, which is enhanced in the presence of its co-substrate ATP (Fig. S3, Table S4). These results overall suggest that BiP-binding induces moderate structural changes in the TPR conformations of FicD and the dynamic characteristics of the TPR domain, including the effect of ATP, are not restricted by the interaction with BiP (Tables S1, S2 and S4).

#### AMPylated BiP

Since FicD can deAMPylate BiP_AMP_, we additionally investigated the interaction of the monomeric FicD^R118C^ with AMPylated BiP. In the presence of BiP_AMP_, FicD^R118C^ shows a major peak for the static, low-FRET TPR-in conformation and the corresponding decrease in the dynamic, high-FRET population (Fig. 3A, black, Fig. 3C Table S2). The conformation of the major static, TPR-in population remains similar with a dye separation of 75 Å (Fig. 3C, Table S3). Previous research established a high-affinity interaction between catalytically inactive FicD and AMPylated BiP^21^. As monomerized FicD^L258D^, which is used in these experiments, is defective in deAMPylation^25^, it is likely that the interaction between BiP_AMP_ and monomeric FicD results in a stabilized enzyme-target complex. In addition, there is a decrease in conformational flexibility of FicD on the observable timescales. This hypothesis is further supported by a significant decrease (ca. 40 %) in the diffusion coefficient of FicD^R118C^ compared to the apo-state protein (Fig. S3, Table S4). Therefore, we conclude that FicD is strongly bound to BiP in the static, TPR-in conformation. The diffusion coefficients determined for the low-FRET (static, TPR-in conformation) and the high-FRET (dynamic) subpopulations are similar, indicating that BiP_AMP_ is also strongly bound to the dynamic populations (Table S16). Interestingly, the simultaneous addition of ATP with BiP_AMP_ restored the dynamic behavior of FicD^R118C^ (Fig. 3F) with population distributions similar to that observed for ATP:FicD in the absence of BiP (Table S1, S3). To verify FicD binding to BiP, we performed a species FCS analysis on the single-molecule bursts that correspond to the static TPR-in state (E<0.4) and the dynamic population fluctuating between the TPR-in and TPR-out conformations (E>0.4). We performed this using the acceptor channel after acceptor excitation to avoid complications from the FRET signal. The species FCS analysis indicated strong binding of FicD to BiP in the dynamic conformations but a higher diffusion coefficient for the static, TPR-in conformation (Table S16). The diffusion coefficient of the static, TPR-in conformation is still significantly lower than for FicD alone indicating transient binding of FicD to BiP in this conformation. In summary, the interaction of monomeric FicD^R118C^ with AMPylated BiP leads to a loss of conformational dynamics, which can be restored by the addition of ATP. This effect may be a result of the competition between the AMP moiety on BiP and free ATP for the same active site of FicD.

### Conformational dynamics involve the 1^st^ and 2^nd^ TPR motifs

To further investigate the conformation of the TPR motifs, we extended the analysis of FicD’s conformational dynamics to alternative labeling positions within the TPR domain, each in combination with S288C as a consistent second labeling position on the Fic domain (Fig. 4A). Residue L104C, which is N-terminal to the first α-helix of the TPR domain, was introduced to investigate the position and dynamics of the 1^st^ TPR motif (FicD^L104C^, Fig. 4A). Despite the similar AV-predicted distances of 60 and 59 Å for apo- and ATP-bound states, respectively (Table S2), two distinct FRET populations were detected for FicD^L104C^:apo at E=0.36 (R=65 Å) and E=0.71 (R= 51 Å) (Fig. 4B, light blue, Table S6). Similar to FicD^R118C^, the addition of ATP altered the population distribution in favor of the high-FRET state (Fig. 4C, yellow, Table S6), whereas the addition of BiP shifted both populations to lower FRET efficiencies (Fig. 4B, blue, Table S6). The pronounced dominance of the low-FRET population in the presence of BiP_AMP_ is conserved for FicD^L104C^ as well (Fig. 4B, black). For all measured conditions, the E-𝜏_𝐷(𝐴)_plots of monomeric FicD^L104C^ exhibit both low and high-FRET populations. The low-FRET population (R=68-72 Å, Table S5) lies on the static FRET line and can thus be assigned to the static TPR-in conformation of FicD (Fig. S4). A fluorescence lifetime analysis revealed significant sub-ms dynamics for the high-FRET populations, except for in the presence of BiP_AMP_. Analogous to FicD^R118C^, these were attributed to conformational switching between the dynamic TPR-in (65 Å) and TPR-out (41-47Å) conformations (Table S3, Fig. S4). Overall, BiP and ATP induced similar effects on the distance population distribution of both 1^st^ TPR motif mutants, which validates the proposed conformational equilibrium of FicD in solution.

**Fig. 4:**
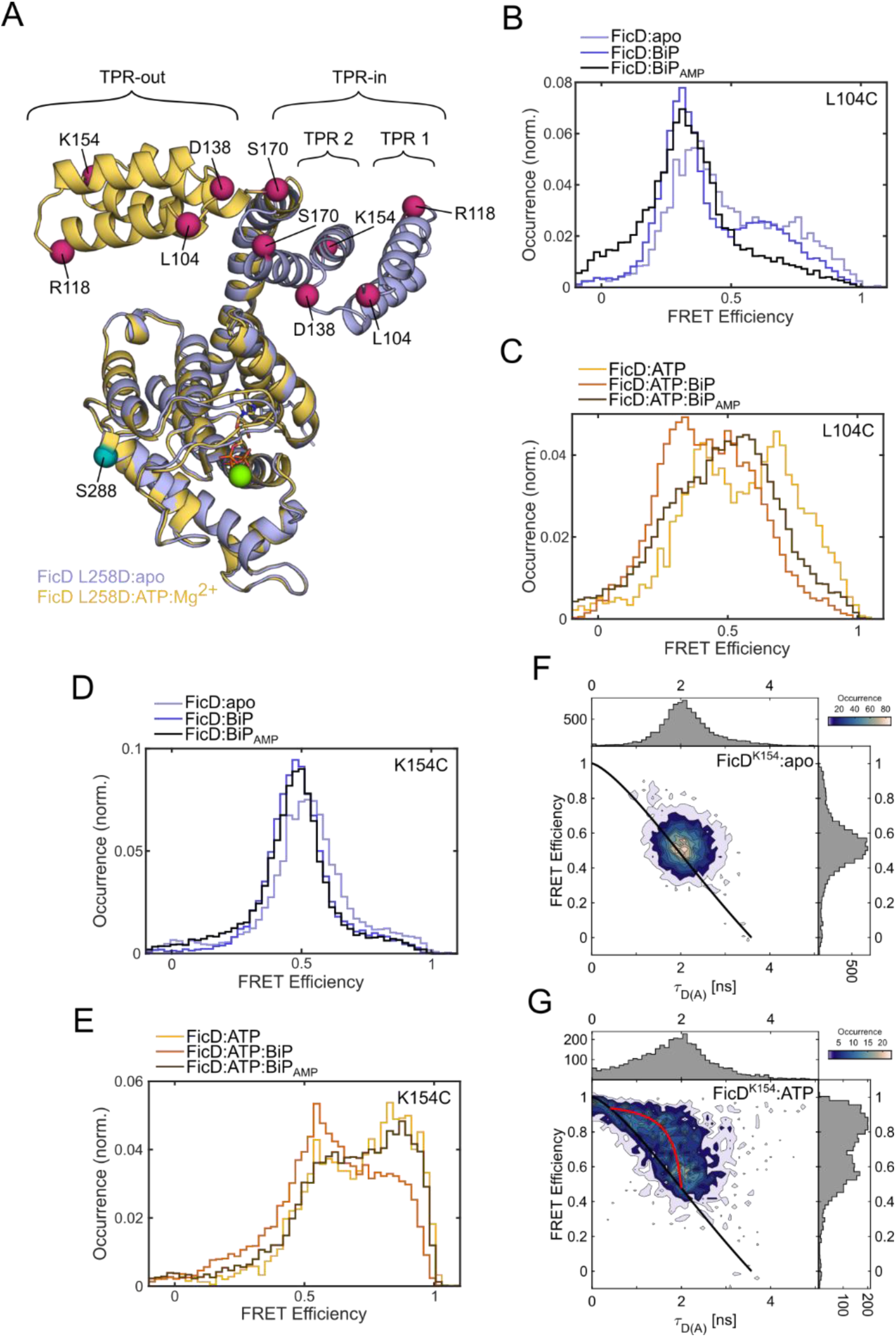
Both TPR motifs of FicD are involved in conformational switching. **A:** Superimposed representations of monomeric FicD^L258D^ crystal structures in the apo-state (light blue, PDB 6I7J (Perera et al., 2019)) and bound to ATP (yellow, PDB 6I7K (Perera et al., 2019), stick representation ATP, green sphere Mg-ion). The various smFRET labeling positions within the TPR domain (magenta) and the Fic domain S288 (teal) are indicated by spheres on the C𝛼-atoms of the amino acid residues. **B:** SmFRET efficiency histograms of FicD^L104C^:apo (light blue) in comparison to FicD^L104C^ in the presence of BiP (blue) and BiP_AMP_ (black). **C**: SmFRET efficiency histograms of FicD^L104C^:ATP (yellow) in comparison to FicD^L104C^:ATP in the presence of BiP (orange) and BiP-AMP (brown). **D:** SmFRET efficiency histograms of FicD^K154C^:apo (light blue) in comparison to FicD^K154C^ in the presence of BiP (blue) and BiP_AMP_ (black). **E**: SmFRET efficiency histograms of FicD^K154C^:ATP (yellow) in comparison to FicD^K154C^:ATP in the presence of BiP (orange) and BiP_AMP_ (brown). **F-G**: 2D plots of donor fluorescence lifetime in the presence of an acceptor (𝜏_𝐷(𝐴)_) vs FRET efficiency (E) for **F:** FicD^K154C^:apo and **G:** FicD^K154C^:ATP. Black and red lines represent the static and dynamic FRET lines, respectively.

Next, we investigated whether the conformational flexibility is conserved throughout the full TPR domain. We fluorescently labeled the 2^nd^ TPR motif at the loop residue K154C in addition to S288C on the Fic domain (FicD^K154C^, Fig. 4A). Here, only a single population at E=0.53 (R=58 Å) was observed for FicD^K154C^ in the apo-state (Fig. 4D, light blue, Table S8), which corresponds well with the expected distance of 62 Å from the AV simulations (Table S3). In the E-𝜏_𝐷(𝐴)_plot, this population appears on the static FRET line (Fig. 4F), indicating a single static conformation. In the presence of ATP, however, a high-FRET population at E=0.86 (R=44 Å) emerges in addition to the slightly shifted static, low-FRET population at E=0.58 (R=56 Å) (Fig. 4E, yellow, Table S7). The observation of a high-FRET state is consistent with the AV simulations for the ATP bound enzyme, where the predicted distance between the dyes decreases to 54 Å (Table S3). The high-FRET efficiency state is also observed for FicD^K154C^ in the presence of BiP or BiP_AMP_ together with ATP, (Fig. 4E, Table S8). The E-𝜏_𝐷(𝐴)_plots for FicD^K154C^ in the presence of ATP indicate that the emerging high-FRET states are dynamic (Fig. 4G, Fig. S5). The endpoints of the dynamic fluctuations were determined to be 41 Å and 61 Å (Table S7) and the lifetime analysis revealed contributions from an additional static state at 71 Å (Fig. S5, Table S7). The fraction of this static, low-FRET state with a fluorescence lifetime 2.8 ns is higher in the presence of BiP and ATP, and BiP_AMP_ and ATP (Table S7). We attribute this conformation at 71 Å to the TPR-in conformation of FicD that is static on the ms timescale, similarly to the results of the FicD^R118C^ or FicD^L104C^ FRET constructs. Furthermore, the static TPR-in state that was detected in the absence of ATP (R=58 Å) is similar to the dynamic, TPR-in state, which is a donor-acceptor separation of 61 Å according to the fluorescence lifetime analysis (Table S7). Therefore, FicD^K154C^ showed 3 distinct conformational states as detected for FicD^R118C^ or FicD^L104C^. However, the apparent nucleotide dependence of the dynamics of FicD^K154C^ stands in stark contrast to the smFRET probes labeled on the 1^st^ TPR motif.

To further explore this difference, we labeled FicD at two additional TPR domain residues, each in combination with S288C: residue D138C located in the loop region between the 1^st^ and 2^nd^ TPR motif, and S170C at the loop between the TPR domain and the helical domain linker of FicD (Fig. 4). Comparable to FicD^K154C^, smFRET measurements of these mutants revealed a single FRET population in the apo-state (Fig. S6A,C, light blue histograms). In contrast to the previous FRET mutants, the AV simulations in both cases predicted the distance between the labeling dyes to increase from the apo-to the ATP-bound state (Table S3). A decrease in FRET efficiency was indeed observed for the D138C construct. Here, a single, high-FRET population is detected in the apo-state and the corresponding E-𝜏_𝐷(𝐴)_ plot indicates moderate conformational dynamics (Fig. S7). In the presence of ATP and BiP or BiP_AMP,_ however, the high-FRET population peak broadens to lower FRET-efficiency values (Fig. S7D,F). The respective E-𝜏_𝐷(𝐴)_ plots indicate dynamic fluctuations between 49 Å and 71-79 Å with the high-FRET population of the apo-state representing one of the endpoints (Fig. S7, Table S9). Interestingly, the addition of ATP did not change the observed behavior unless BiP was present, in contrast to other labeling positions.

For FicD^S170C^, the single distance population of the apo-state is consequently broadened in the presence of ATP, irrespective of the addition of BiP or BiP_AMP_. (Fig. S6). The E-𝜏_𝐷(𝐴)_ plots for these conditions indicate an increased dynamic averaging of this population due to fluctuations between conformations with donor-acceptor dye separations of 49 Å and 68-72 Å (Fig. S8, Table S10). These values differ to the expected distances given the AV simulations suggesting that the crystal structures do not fully describe the structure of the dynamic protein. In summary, both TPR motifs of FicD undergo conformational dynamics. The low-FRET and high-FRET populations observed for constructs labeled on the 1^st^ TPR motif (L104C, R118C) can be attributed to the TPR-in and TPR-out conformations of FicD observed in crystallography (Fig. 1B). In contrast, the conformations detected for the 2^nd^ TPR motif differ from what is expected from these crystal structures.

### ATP influences the dynamics of the TPR-in - TPR-out transitions

After the observation of conformational dynamics from the E-𝜏_𝐷(𝐴)_plots for several FicD constructs, we performed a dynamic photon distribution analysis (PDA)^32^ to quantify the timescales of the TPR motions of FicD. By analyzing the smFRET histograms with different time binnings, PDA can extract the contribution of each state to the FRET distribution and the rate constants of the dynamic switching between different conformations. Based on the conformational states obtained from the fluorescence lifetime analysis, the dynamic TPR-in and TPR-out conformations for FicD^R118C^ were defined with fluorophore distances of 43 Å and 65 Å, respectively (Fig. 5A, blue and orange, respectively). We define the transition rates between these two conformations as k_out_ for the switch from TPR-in to TPR-out and k_in_ for the reverse transition (Fig. 5A, yellow). We define a third conformational state as the static, TPR-in conformation, with a distance between the fluorophores of 71 Å in the PDA model to describe the additional, static low-FRET population, which was observed under all conditions for FicD^R118C^ (Fig. 5A, green). To account for acceptor blinking or impurities, we included another static, low-FRET population at 92 Å (Fig. 5A, purple) into the model. We globally applied this 2-state dynamic model to all datasets to extract the transition rates of the TPR motion under each condition (Fig. 5B,C, Fig. S9, Table S11). In the absence of ATP or BiP, the transition rates of k_in_= 1.08 ms^-1^ and k_out_= 0.88 ms^-1^ (Table S8) indicate a faster transition from the TPR-out to the TPR-in conformation, which suggests a preference of the apo-state of FicD^R118C^ to adopt the dynamic TPR-in conformation. In addition, a higher fraction of molecules are in the static TRP-in population under these conditions (Table S1). Substantially faster rates were obtained for FicD^R118C^:ATP with k_in_= 1.78 ms^-1^ and k_out_= 2.57 ms^-1^. Here, the rise of k_out_ over k_in_ suggests a faster transition of FicD^R118C^ to the dynamic TPR-out than to the TPR-in conformation, which correlates with the increase of the dynamically interconverting, high-FRET population observed in the E-𝜏_𝐷(𝐴)_ plots of FicD^R118C^:ATP (Fig. 2D). Also here, there is an increase in the fraction of dynamic molecules and a decrease in the static TRP-in conformation (Table S1). In the presence of BiP, the ratio between k_out_ and k_in_ decreases (Fig. 5D), highlighting a preference of the TPR-in conformation and a higher fraction of TRP-in static molecules are observed. The lowest ratio of k_out_ to k_in_ was obtained for BiP_AMP_ (Fig. 5D), where the rate of the TPR domain moving in becomes dominating and the dynamically interconverting population is depleted significantly (Fig. 3A,C). We therefore interpret the ratio of k_out_/k_in_ as a measure for FicD’s tendency to adopt the dynamic TPR-out conformation.

**Fig. 5:**
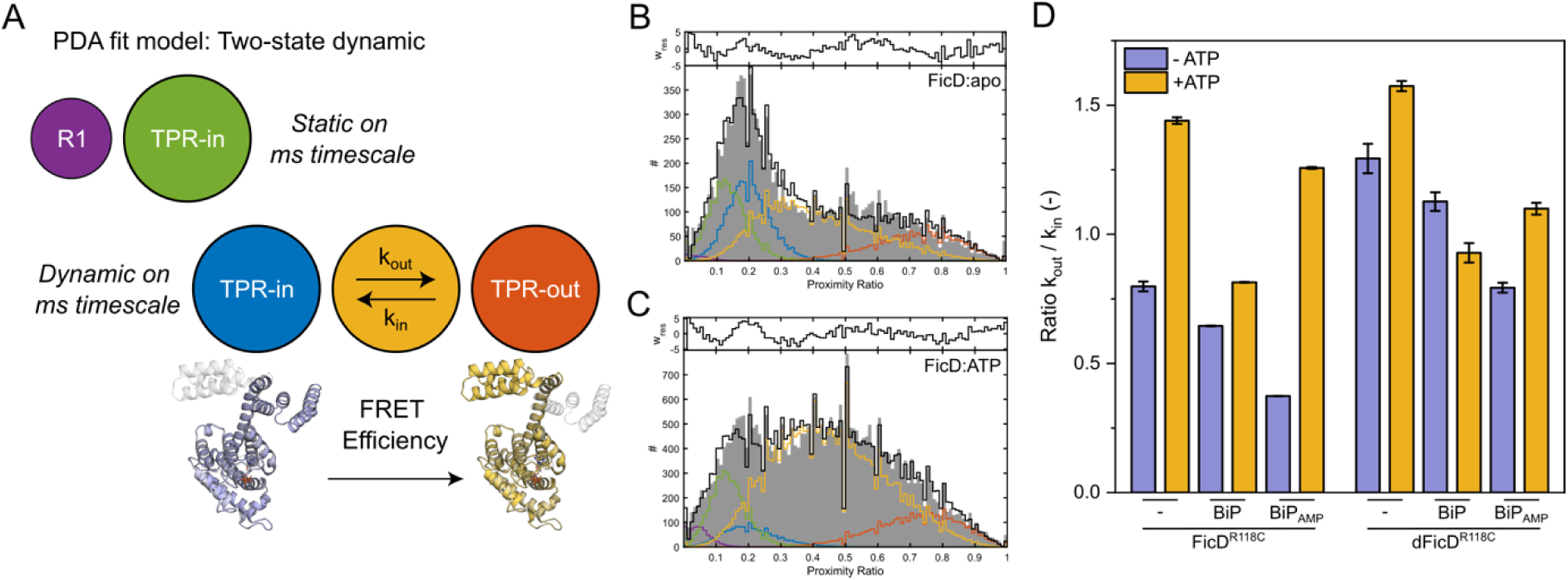
Analysis of the conformational states and dynamics of FicD^R118C^ using the dynamic photon distribution analysis (PDA). **A:** Schematic of the PDA model with two static (purple, green) and two dynamic (blue, orange) conformational states that were incorporated into a global PDA fit model for FicD^R118C^. The dynamically interconverting population is shown in yellow. **B-C:** For dynamic PDA, uncorrected FRET efficiency (proximity ratio) histograms with time binnings of 0.5 ms, 0.75 ms and 1.0 ms were used to calculate the distance distribution for different FRET populations and dynamics between them. The data for the time window size of 0.75 ms is shown. Proximity ratio histograms of FicD^R118C^ for **B:** the apo-state and **C:** in the presence of ATP were fitted using the 2-state dynamic PDA model. Results from the PDA fits are given in Table S8. **D:** Comparison of the ratio k_out_/k_in_ obtained from PDA analysis of monomeric (L258D) or dimeric FicD labeled at R118C and S288C.

To quantify the kinetics of FicD^L104C^, we globally applied the same 2-state dynamic PDA model to all measurement conditions (Fig. S10, Table S12). The distances used for the dynamic populations during the PDA analysis were fixed based on the fluorescence lifetime analysis: TPR-in (60 Å) and TPR-out (42 Å) conformations. An additional state with a distance separation of 65 Å was also included to account for the static TPR-in conformation. Although the addition of ATP does not generally increase the ratio of k_out_ to k_in_ for FicD^L104C^, similar trends as for FicD^R118C^ are observed in the transition rates for the interaction with BiP (Fig. S12): the presence of BiP and BiP_AMP_ reduce the k_out_ to k_in_ ratio. Again, this effect is most evident for BiP_AMP_ and can be reverted by the addition of ATP.

### The 2^nd^ TPR motif becomes dynamic in the presence of ATP

For FicD^K154C^, dynamics were observed only for the measurements in the presence of ATP. One dominant population is observed with a FRET efficiency of ca. 0.5. However, to describe the conformation of FicD for all conditions with a similar model, we performed a global fit to the conditions without ATP using a 3-state static model. The distance distributions were assumed to be Gaussian and the position for the peaks of the distance distributions were fixed to 41 Å, 57 Å and 74 Å. These values were determined via initial PDA fits to the individual smFRET histograms and fit to the fluorescence lifetime distributions. The results from the global PDA analysis describe both the smFRET histograms and the fluorescence lifetime distributions well (Fig. S11, Table S7, Table S13). For the measurements in the presence of ATP, we included dynamic transitions between the 41 Å and 57 Å populations along with the static population at 74 Å (Fig. S11, Table S13). Overall, the dynamic rates for switching between the TPR-in (57 Å) and TPR-out (41 Å) conformations are reduced compared to FicD^R118C^ or FicD^L104C^ in the presence of ATP, indicating slower fluctuations (Table S13). However, also for FicD^K154C^, the addition of BiP reduced the ratio of k_out_ to k_in_, highlighting the bias towards the TPR-in conformation (Fig. S12) and the fraction of the static, TPR-in state increases (Table S13). These results indicate that the conformational shifts between TRP-in and TRP-out states of FicD extend into the 2^nd^ TPR motif. However, in contrast to the 1^st^ TPR motif, the fluctuations first become prominent in the presence of ATP.

In summary, we quantified the transition rates of the conformational switching of the TPR domain for several FicD mutants (R118C, L104C, K154C) in the absence and presence of ATP (Fig. 5E, Fig. S12). Notably, the addition of ATP generally increases the absolute transition rates of FicD compared to the apo-state, suggesting overall faster dynamics of the enzyme in the presence of its co-substrate. Moreover, the presence of BiP or BiP_AMP_ reduces the k_out_/k_in_ ratio (Fig. 5E, Fig. S12) as well as the fraction of the static TRP-in state, thus favoring TPR-in conformation. This preference for the TPR-in conformation presumably is a result of enzyme-substrate complex formation where FicD binds BiP in the TPR-in conformation^24,25^.

### FicD’s conformational dynamics are conserved for the dimeric protein

In contrast to the monomerized FicD^L258D^ mutant, no rearrangements of the TPR domain were observed for dimeric wildtype FicD in crystal structures of the apo- or ATP-bound states^20,21^. This observation raises the question of whether the dynamics for the monomeric FicD^L258D^ are conserved for the dimeric protein. Since monomeric and dimeric FicD fulfil antagonizing catalytic functions (AMPylation versus deAMPylation, respectively)^21^, a change in the enzyme’s dynamics is conceivable. Therefore, we extended our investigation to the more physiological dimeric enzyme. To ensure dimer formation in the smFRET experiments, we added a high excess of unlabeled, wild type FicD (50 nM unlabeled vs ∼ 100 pM labeled protein) to the measurements, which is well above the K_d_ for dimerization (1 nM for FicD^21^). Dimer exchange between labeled and unlabeled FicD then ensures a maximum of one labeled subunit per detectable dimer. Dimer formation of dFicD^R118C^ was validated FCS, where slower diffusion coefficients are detected under all measurements conditions when compared to the monomeric enzyme (Fig. S3, Table S4).

In the absence and presence of ATP, the FRET efficiency histograms and corresponding E-𝜏_𝐷(𝐴)_ plots obtained for dimeric FicD labeled at residues R288C and R118C (dFicD^R118C^) display a similar conformational landscape as obtained for the monomeric FicD^R118C^ (Fig. 6, Fig. S13 and Table S14). Consistent with the previous results, the apo-state of dFicD^R118C^ reveals a static, low-FRET population at E=0.28 (69 Å), which is assigned to the static TPR-in conformation and a dynamic, high-FRET population at E=0.75 (49 Å), which is attributed to a flipping motion of the TPR domain between dynamic TPR-in and TPR-out conformations (Table S15). The conformational equilibrium, however, is shifted towards the dynamic, high-FRET population in the dimeric enzyme and the static, low-FRET population is less abundant compared to the monomer (compare Table S2 and Table S15). The addition of ATP further shifted the conformational equilibrium towards the dynamic, high-FRET population in all measurements (Table S14, S15). An additional, a small high-FRET population (E=0.91, 40 Å) is observed in all conditions with ATP, which we attribute to photophysics of the fluorophores rather than an additional conformational state of the dimeric FicD^R118C^ (Table S15). From the fluorescence lifetime analysis, we concluded that the dimeric FicD^R118C^ shows similar dynamic interconversions between the TPR-in and TPR-out states (Fig. S13, Table S14) as observed for the monomeric enzyme.

**Fig. 6:**
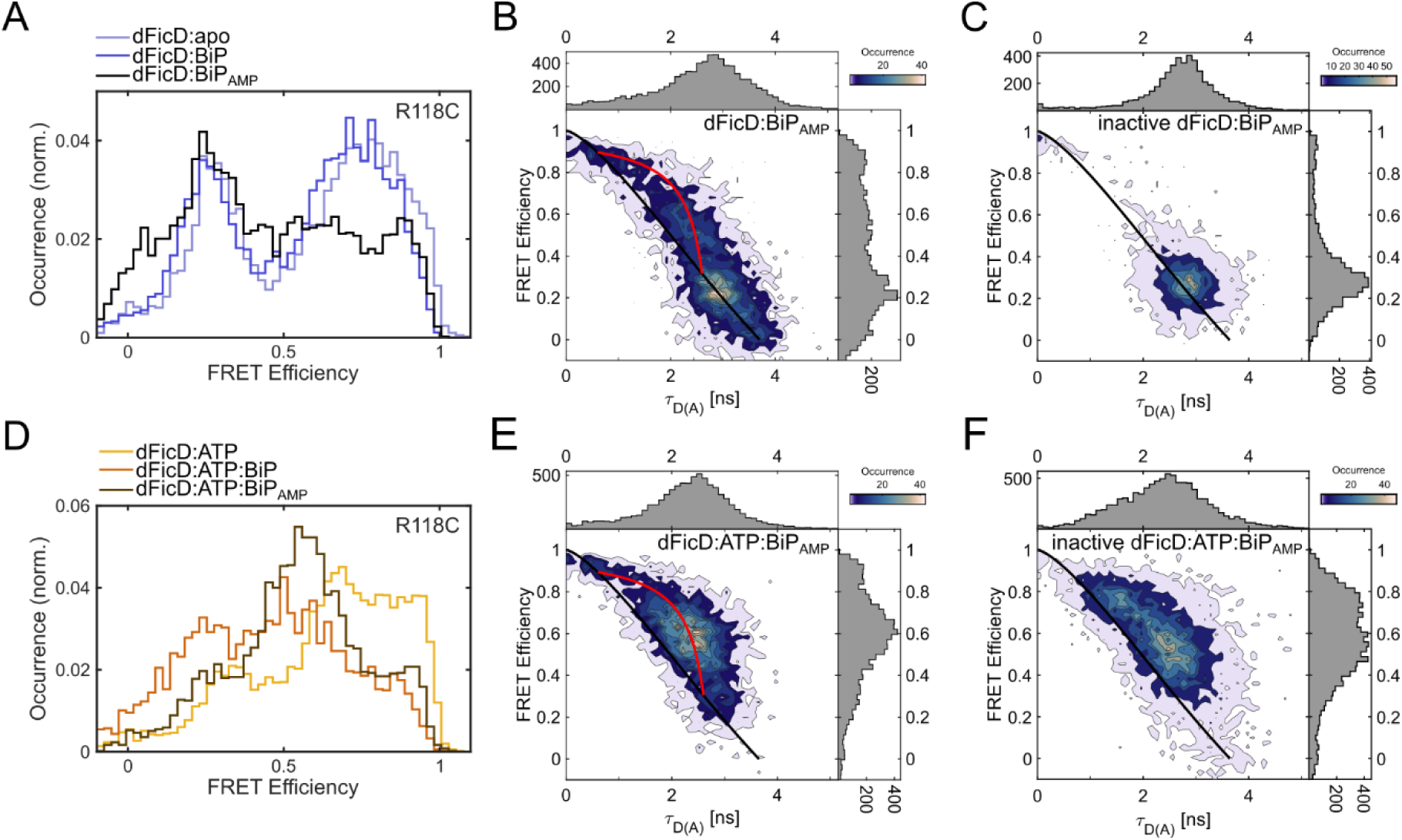
The conformational equilibrium of dFicD^R118C^. **A:** SmFRET efficiency histograms of dFicD^R118C^:apo (light blue) in comparison to dFicD^R118C^ in the presence of BiP (blue) and BiP_AMP_ (black). **B-C:** 2D plots of donor fluorescence lifetime in the presence of an acceptor (𝜏_𝐷(𝐴)_) vs FRET efficiency (E) for **B:** dFicD^R118C^:BiP_AMP_ and **C:** inactive dFicD^R118C^ ^H363A^:BiP_AMP_ FicD^R118C^. Black and red lines represent the static and dynamic FRET lines, respectively. **D**: SmFRET efficiency histograms of dFicD^R118C^:ATP (yellow) in comparison to dFicD^R118C^:ATP in the presence of BiP (orange) and BiP_AMP_ (brown). **E-F:** 2D plots of donor fluorescence lifetime in the presence of an acceptor (𝜏_𝐷(𝐴)_) vs FRET efficiency (E) for **E:** dFicD^R118C^:ATP:BiP_AMP_and **F:** inactive dFicD^R118C^ ^H363A^:ATP:BiP_AMP_ FicD^R118C^. Black and red lines represent the static and dynamic FRET lines, respectively.

Most importantly, the dynamic, high-FRET population of dFicD^R118C^ is also observed in the presence of AMPylated BiP (Fig. 6A,B). Although the static TPR-in conformation is favored in this condition (Table S15), dynamic fluctuations were observed between the dynamic TPR-in (69 Å) and TPR-out (46 Å) conformations (Fig. 6B, Table S14). This single condition stands in stark contrast to monomeric FicD^R118C^, where the dynamic, high-FRET population is strongly depleted in the presence of BiP_AMP_ (Fig. 3A,C). To gain further insights into the interaction of FicD with BiP, we analyzed the smFRET data using FCS. When analyzing the ACF from the entire measurement, the slowest diffusion coefficients were observed for both FicD and dFicD in the presence of BiP_AMP_ (Table S4). As a next step, we performed species FCS on the static and dynamic subpopulations. Similar diffusion coefficients were observed for the static TPR-in and the dynamic conformation with the exception of monomeric FicD in the presence of BiP_AMP_ and ATP, and dimeric FicD in the presence of BiP_AMP_ (Table S16). We then analyzed FicD and dFicD interactions with BiP_AMP_ in the absence and presence of ATP using a 2-component fit to the ACFs where the diffusion coefficients were fixed to that of enzyme only and enzyme bound to BiP_AMP_. The fraction of bound enzyme for each subpopulation is given in Table S17. For the dynamic subpopulation, the fraction of bound enzymes does not change upon addition of ATP. In contrast, the fraction of bound molecules in the static, TPR-in state decreases for both FicD and dFicD. However, there is still a significant fraction of bound molecules interacting with BiP_AMP_ indicating the presence of transient interactions (Table S17). Thus, one can rule out the interpretation that the static, TRP-in conformation that of FicD bound to BiP and the dynamic conformation is only present for FicD alone. Furthermore, due to the high excess of BiP_AMP_ (100 µM) to enzyme (50 nM unlabeled dFicD) and the slow catalyzation rate of k_cat_ (10^-2^ s^-1^) for deAMPylation of BiP by dFicD^25^, only minimal deAMPylation of the sample is expected during the 2 h measurement. This assumption is confirmed by the 2D mean macrotime versus FRET efficiency plots of dFicD^R118C^:BiP_AMP_, as the dynamic high-FRET population does not increase during the measurement due to increasing amounts of unmodified BiP (Fig. S14). In fact, there could be a potential decrease in the dynamic high-FRET population time.

As deAMPylation active dFicD is dynamic when bound to BiP_AMP_, the catalytic process may be coupled to the dynamics. To investigate this possibility, we measured a fluorescently labeled, catalytically inactive dFicD^R118C^ ^H363A^ mutant in the presence of BiP_AMP_ (Fig. S15C). Here, the dynamic behavior was almost completely depleted, similar to the results for monomeric FicD^R118C^ (Fig. 3A,C and Fig. 6C), supporting the hypothesis that dynamic switching of the TPR domain of dFicD^R118C^ may correlate with its deAMPylation activity. Upon addition of ATP to the BiP_AMP_:FicD complex, the monomeric FicD^R118C^, inactive dFicD^R118C^ and the active dFicD^R118C^ all exhibit dynamics (Fig. 6D and Table S14).

Other differences between monomeric and dimeric FicD were identified in the PDA analysis (Fig. S16 and Table S16). The absolute rates of dFicD^R118C^ are overall lower compared to monomeric FicD^R118C^, indicating slower conformational switching. However, the ratio of k_out_ to k_in_ is increased for the dimeric enzyme, in line with the higher preference of dFicD^R118C^ to adopt the dynamic TPR-out conformation for all measured conditions (Fig. 5D). The effect of ATP, in general, is less pronounced for the dimeric enzyme. More importantly, ATP does not increase but decrease k_out_/k_in_ for dFicD^R118C^:BiP in contrast to all other conditions (Fig. 5D). Here, the enhancing effect of ATP might be reduced by the formation of stabilized dFicD^R118C^:ATP:BiP ternary complexes, as dimeric FicD is AMPylation defective^21^. We conclude that the dFicD deAMPylase undergoes dynamic conformational fluctuations in solution similar to those observed for the monomeric AMPylase, but that the distribution between these conformations differs in favor of the dynamic TPR-out conformation.

## Discussion

In this study, we investigated the conformational interdomain dynamics of the human FicD enzyme using smFRET. The resulting data reveal dynamic conformational rearrangements of FicD’s TPR domain in solution, leading to the definition of an equilibrium between three distinguishable conformational states: the static TPR-in conformation, which corresponds to the crystal structure of FicD bound to BiP (PDB: 6ZMD^24^) and a dynamic, interconverting population fluctuating between a TPR-in and a TPR-out conformation (Fig. 7). The annotation of conformational states in this study are based on AV calculations using crystal structures of FicD in the apo-state (TPR-in, PDB 6I7J^21^), of FicD bound to ATP (TPR-out, PDB: 6I7K^21^), or of FicD complexed with BiP (TPR-in, PBD: 6ZMD^24^) (Fig. 2, Fig. S2, Table S3) and the measured FRET efficiencies. Especially the experimental dye separations obtained for monomeric and dimeric FicD^R118C^ are comparable to the distances predicted by AV simulations (Table S1, S3 and S14). All three states are consistently detected for monomeric and dimeric FicD (dFicD) labeled at residue R118C and confirmed by the labeling of additional positions (L104C, K154C) on the 1^st^ TPR motif.

**Fig. 7:**
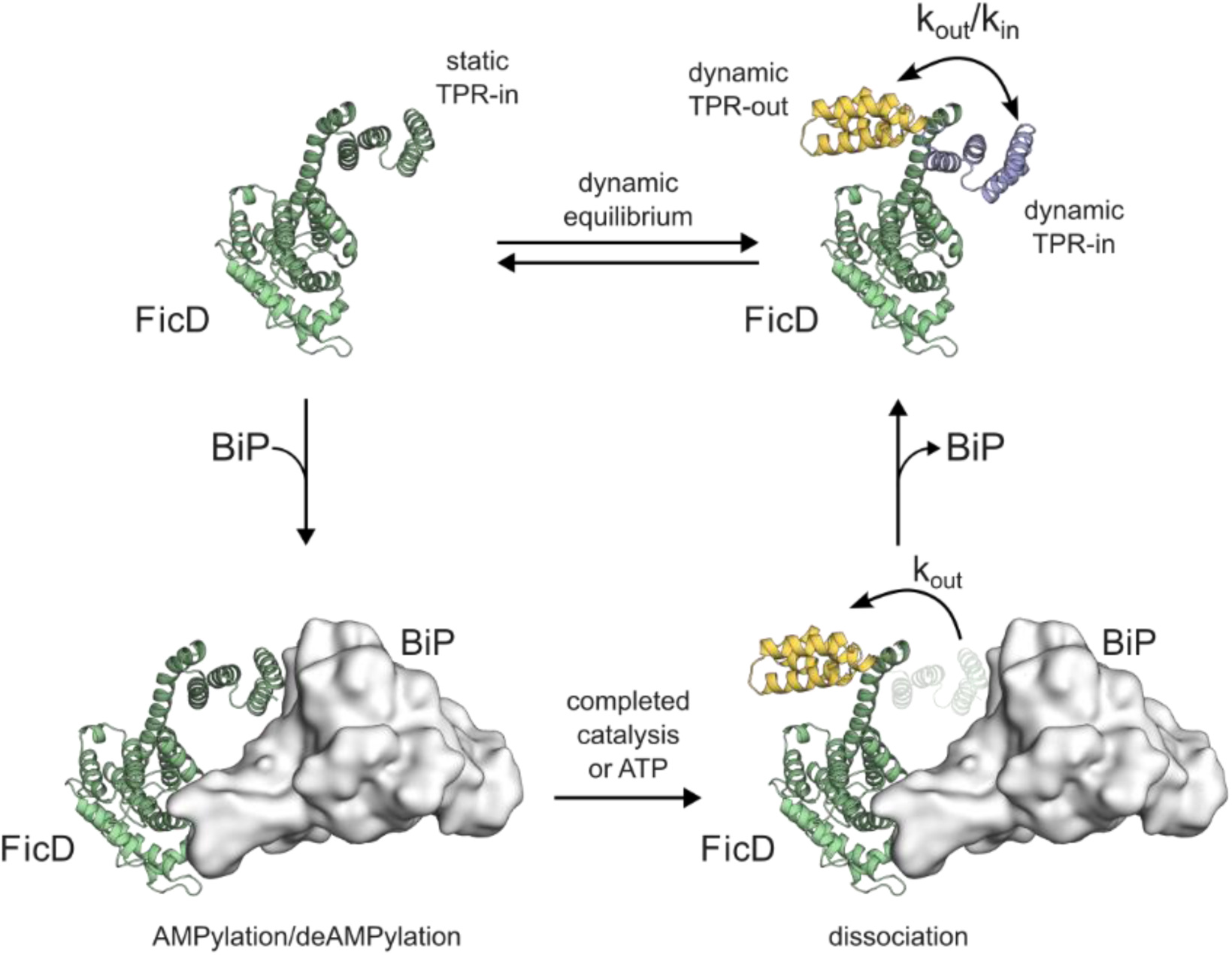
Dynamics of the TPR domain of FicD. A schematic representation of the conformational landscape of FicD in the absence and presence of BiP. According to our smFRET data, the dynamically interconverting TPR-in and TPR-out as well as the static TPR-in conformation of FicD exist in equilibrium. The static TPR-in conformation likely equals the conformation of FicD in complex with BiP. Hypothetically, the TPR domain adopts the TPR-out conformation upon completed catalysis or binding of FicD to ATP, enabling dissociation of the enzyme-target-complex.

Transitions between the TPR-in and TPR-out conformations are observed on the millisecond timescale indicating that the presence of the TPR-out conformation is transient in nature. ATP shifts the conformational equilibrium towards the dynamic state whereas BiP_AMP_ binding promotes the static state. For mutants on the 2^nd^ TPR motif (K154C, D138C, S170C), we detected emerging dynamics only in the presence of ATP. This suggests a decoupling of the dynamics observed in the different motifs of the TPR domain. Also, minor differences in solution structures are expected as the measured FRET efficiencies for the 2^nd^ TPR motif FRET constructs FicD^D138C^ and FicD^S170C^ showed deviations from the AV predictions. Considering the static character of crystal structures and potential crystal-packing artifacts, a deviation to data obtained in solution is not surprising. In this respect, it is interesting to note that the TPR domain of each FicD molecule in the FicD:ATP crystal structure^21^ is intimately intertwined with a neighboring one (Fig. S18). Our in-solution smFRET data can provide an explanation of this observation: The TPR motifs’ fast conformational switching may constantly sample different structural states of FicD, of which one particular state permits the formation of the interfaces observed in the crystal. On the other hand, these extensive crystal contacts likely lock the TPR domain in one of many possible conformations, which provides a possible explanation behind the deviation between some of the AV simulations and experimental smFRET data. Hence, we conclude, that although the TPR orientation in the crystal structure of FicD:ATP (PDB 6I7K^21^) is not identical to the TPR-out conformation in solution, it may serve as a helpful approximation.

Consistent for all the measured mutants, the fluorescence lifetime and photon distribution analyses (PDA) confirmed that the presence of ATP enhances the dynamic flipping motion of FicD (Fig. 5E, Fig. S12). More specifically, the increased ratio of k_out_ over k_in_ results in a shift of the equilibrium in favor of the dynamic interconverting TPR-out population of FicD. In contrast, binding to BiP causes the opposite effect and reduces the contribution of the dynamic population in favor of the static, TPR-in population. This effect was the strongest for AMPylated BiP. We further compared the conformational dynamics upon interaction with BiP between the monomeric FicD^L258D^ mutant and the dimeric protein. Here, we observed diverging conformational dynamics between monomeric and dimeric FicD in the presence of BiP and BiP_AMP_: while ATP enhances the dynamic flipping motion (k_out_/k_in_ ratio) in all other cases, it is reduced exclusively for dimeric and AMPylation deficient FicD in the presence of BiP and ATP (Fig. 5D). Even more pronounced is the depletion of the conformational dynamics of deAMPylation deficient enzymes (i.e. the monomeric L258d or inactivated H363A mutants) in the presence of BiP_AMP_. While the dimeric and deAMPylation active enzyme retains its dynamics, the deAMPylation inactive mutants persist in the static TPR-in conformation (Fig.3C, 6B,C). This observation suggests that the flipping motion of the TPR domain may be linked to the catalytic activity of FicD. Hypothetically, the dynamic switching of the TPR domain correlates with the AMPylation or deAMPylation activities. Since the TPR domain is required for binding to BiP^24,25^, the conformational rearrangement would contribute to dissociation of the post-catalytic complex and thereby enhance catalytic turnover (Fig. 7). Consistent with this hypothesis, the artificial fixing of the TPR domain to the connecting helix of the Fic domain reduces the deAMPylation activity of FicD^25^. This model can be extended by the observation that ATP recovered the dynamics of monomeric FicD in the presence of BiP_AMP_ (Fig. 3C,F). Binding to ATP has been proposed to allosterically weaken the affinity between FicD and BiP^21^. Similarly, our FCS analysis showed a decrease in the FicD^R118C^:BiP_AMP_ complex formation upon addition of ATP for both monomeric and dimeric FicD (Table S17). This effect may be explained by the increased flipping motion of FicD’s TPR domain upon nucleotide binding (Fig. 5D), which might stimulate complex dissociation irrespective of catalysis. ATP might thus prevent the accumulation of high-affinity enzyme-substrate complexes and contribute to efficient activity cycling of FicD^21^. However, it must also be considered that the competition between the AMP moiety attached to BiP and ATP for the active site of FicD reduces the complex formation and allows for dynamic flipping of freely diffusing FicD molecules (Fig. 3F, Table S3).

In summary, using smFRET, we could show that FicD undergoes conformational fluctuations in solution and while bound to BiP. These conformational fluctuations persist in the active dimer, even when bound to AMPylated BiP, although it was not observed for non-catalytic monomeric FicD^L258D^ nor the inactive dimer FicD ^H363A^. This suggests that not only is FicD dynamic, but that the dynamics are essential for regulating FicD’s function. These measurements also lay the groundwork for future studies that investigate the allosteric signaling between the nucleotide binding site and the TPR domain.

## Materials and methods

### Plasmid Construction

All FicD constructs were generated based on a pSF421 expression vector containing a 6xHis-GFP-tag followed by a Tobacco Etch Virus (TEV) protease cleavage site and FicD 102-458 (NCBI Accession NP_009007.2)^24^. For monomeric FicD, L258D and additional point mutations were introduced by site directed mutagenesis using the Q5^®^ Site-Directed Mutagenesis Kit (NEB) according to the manufacturer’s instructions. For dimeric FicD, GFP was exchanged for a small ubiquitin-like modifier (SUMO) pro-peptide. Vector and insert were amplified using overlapping primers and Q5^®^ High-Fidelity DNA Polymerase (New England Biolabs, NEB), separated by agarose gel electrophoresis and purified with the Monarch^®^ DNA Gel Extraction Kit (NEB). The fragments were ligated by sequence-and ligation-independent cloning (SLIC)^33^ using T4 DNA Polymerase (NEB). Point mutations were introduced as described above. After transformation into Mach1 cells (Invitrogen), cultivation and purification with the PureYield™ Plasmid Miniprep System (Promega), positive clones were confirmed by Sanger Sequencing (Microsynth AG). BiP constructs within a pProEx vector comprised a N-terminal 6xHis-tag, a TEV protease cleavage site and human BiP 19-654 (NCBI Accession NP_005338.1) as described previously^24^.

### Recombinant Protein Expression

For recombinant expression of FicD and BiP mutants, 100 ng plasmid DNA were transformed into chemically competent *E. coli* Rosetta DE3 cells (Novagen). After incubation at 37 °C overnight on LB-Agar supplemented with appropriate selection antibiotics, multiple colonies were harvested into antibiotic containing LB and incubated for 4-5 h at 37 °C. Precultures were diluted 25-fold into fresh medium and grown at 37 °C and 180 rpm until an OD_600nm_ of 0.5-0.7. Expression was induced with 0.5 mM isopropyl-β-D-thiogalactopyranosid (IPTG) and the temperature reduced to 22 °C for overnight expression. Cells were harvested at 6000 xg for 15 min, washed with 1x phosphate buffered saline (PBS) followed by a second centrifugation and freezing of the pellets at −20 °C until purification.

### Bacterial Cell Lysis

Cell pellets were thawed on ice and resuspended in 50-100 mL cold Buffer A per 1 L expression culture (compositions for each respective protein as described below). After incubation with DNaseI (AppliChem) for 10 min on ice, cells were lysed with a Constant Cell Disruption Unit (Constant Systems) at 1.8 kbar. The lysates were directly supplemented with 1 mM phenylmethanesulfonyl fluoride (PMSF) and 30 mM (FicD) or 25 mM (BiP) imidazole and cleared by centrifugation at 50000 xg at 4 °C for at least 30 min.

### Protein Purification

#### Purification of FicD

Cleared lysates were passed over a 5 mL EconoFit Nuvia IMAC Column (Bio-Rad) equilibrated in Buffer A (50 mM Hepes-NaOH pH 8, 500 mM NaCl, 1 mM MgCl2, 1 mM β-mercaptoethanol (β-ME)) supplemented with 30 mM imidazole. The column was washed with 10 CV of the same buffer, followed by additional washing steps with 10 CV of Buffer W (50 mM Hepes-NaOH pH 8, 1000 mM NaCl, 10 mM MgCl2, 10 mM imidazole, 1 mM β-ME) and 5 CV of Buffer W supplemented with 2.5 mM ATP. The fusion proteins were eluted by a gradient from 30-350 mM imidazole in Buffer A and protein containing fractions were pooled. Monomeric FicD mutants, expressed as 6xHis-GFP fusion, were mixed with in-house prepared TEV protease in a mass ratio of 1:20 and dialyzed against Dialysis Buffer (20 mM Hepes-NaOH pH 8, 25 mM NaCl, 1 mM MgCl2, 1 mM β-ME, 5 % glycerol) at 4 °C overnight. The dialysed sample was supplemented with 25 mM Imidazole and passed over a 5 mL HiTrap Q HP Column (Cytiva) coupled to an EconoFit Nuvia IMAC Column for simultaneous anion exchange chromatography and removal of all residual 6xHis-tagged fragments via IMAC. The columns were equilibrated and washed with Dialysis Buffer supplemented with 25 mM Imidazole. Elution was performed by gradually increasing salt concentrations in the Dialysis Buffer, with monomeric FicD eluting at 100-160 mM NaCl. Fractions of interest were pooled for size exclusion chromatography (SEC). Dimeric FicD mutants were expressed as 6xHis-Sumo fusion due to insufficient separation from residual, uncleaved fusion protein when expressed as 6xHis-GFP fusion. Respective pools from IMAC were mixed with in-house prepared Ulp1 protease in a ratio of 1:40 (w/w) prior to dialysis against Dialysis Buffer at 4 °C overnight. The dialysed sample was passed over a 5 mL HiTrap Q HP Column (Cytiva) to remove the 6xHis-Sumo-tag. The column was equilibrated and washed with a Dialysis Buffer. Elution was performed by gradually increasing salt concentrations in the Dialysis Buffer, with dimeric FicD eluting at 200-250 mM NaCl. Fractions of interest were pooled for SEC. Prior to SEC, the samples were concentrated to a volume ≤ 2 mL using Amicon® Ultra Centrifugal Filter units (Merck Millipore) and centrifuged for 5 min at 21000 xg, 4 °C. The sample was injected onto a HiLoad 16/600 Superdex 75 pg column (Cytiva) equilibrated in Buffer C (20 mM Hepes-KOH pH 7.4, 150 mM KCl, 1 mM MgCl2, 1 mM Tris(2-carboxyethyl)phosphine hydrochloride (TCEP), 5 % glycerol). Elution peaks were analyzed via sodium dodecyl-sulphate polyacrylamide gel electrophoresis (SDS-PAGE) and the fractions of highest purity were pooled, concentrated using Amicon® Ultra Centrifugal Filter units and stored at −80 °C after freezing in liquid nitrogen. The molar concentration of the samples was determined via UV-Vis spectroscopy at 280 nm (ε = 29340 M^-1^cm^-1^).

#### Purification of BiP

Cleared lysates containing 6xHis-tagged BiP T229A were passed over a 5 mL EconoFit Nuvia IMAC Column, which was equilibrated and washed with BiP-Buffer A (50 mM Hepes-NaOH pH 7.4, 400 mM NaCl) supplemented with 25 mM imidazole. Elution was performed using a gradient from 25-350 mM imidazole in BiP-Buffer A. Protein containing fractions were pooled, mixed with in-house prepared TEV protease in a ratio of 1:40 (w/w) and dialyzed against BiP-Dialysis Buffer (20 mM Hepes-NaOH pH 7.4, 100 mM NaCl) at 4 °C overnight. The dialysed sample was passed over a 5 mL EconoFit Nuvia IMAC Column to remove the 6xHis-tag while the BiP containing flow-through was collected. SEC was performed as described above, using a HiLoad 16/600 Superdex 200 pg column (Cytiva) equilibrated in HKM-Buffer (20 mM Hepes-KOH pH 7.4, 150 mM KCl, 10 mM MgCl_2_). Elution peaks were analyzed via SDS-PAGE and the fractions of highest purity were pooled, concentrated to approximately 500 µM using Amicon® Ultra Centrifugal Filter units. The molar concentration of the samples was determined via UV-Vis spectroscopy at 280 nm (ε = 30370 M^-1^cm^-1^). The concentrated samples were stored at −80 to −75 °C after freezing in liquid nitrogen.

#### Preparative AMPylation of BiP

To AMPylate BiP in preparative amounts, 10 mL reactions containing 85 nmol purified BiP, 4.25 nmol FicDE234G-L258D and 1.5 mM ATP in Assay-Buffer (25 mM Hepes-KOH pH 7.4, 100 mM KCl, 4 mM MgCl2, 1 mM CaCl2) were incubated for 16 h at 23 °C. The reaction was subsequently concentrated and subjected to SEC in HKM-Buffer as described for unmodified BiP above. Intact LC-MS analysis was used to confirm ≥ 95 % modification.

#### Nano differential scanning fluorimetry

Purified FicD mutants were diluted to 0.15 mg/mL in Buffer C and centrifuged at 20,000 xg for 5 min at 4 °C before loading into Prometheus Standard Capillaries (NanoTemper Technologies). Protein fluorescence at 350 nm (F_350_) and 330 nm (F_330_) was measured by a Prometheus nDSF (NanoTemper Technologies) device over a temperature range from 15-80 °C. Data was analyzed using the MoltenProt Webserver^34,35^. Melting temperatures (T_m_) were obtained from the inflection points of the F_350_/F_330_-ratio vs. temperature plots and averaged over at least three technical replicates.

#### Fluorescent labeling of proteins

Lyophilized dyes (ATTO-TEC GmbH) were dissolved in anhydrous DMSO, flushed with nitrogen and stored in 5-20 µL aliquots at −80 to −75 °C. Molar dye concentrations were determined by UV-Vis spectroscopy using extinction coefficients and maximal absorption wavelengths as provided by the manufacturer. All steps during fluorescent labeling of FicD cysteine mutants were performed in degassed FL-Buffer (25 mM Hepes-KOH pH 7.4, 100 mM KCl, 1 mM MgCl_2_, 0.05 mM TCEP, 5 % glycerol). 10 nmol purified FicD were reduced with 10 mM dithiothreitol (DTT) in a reaction volume of 200 µL for 10-20 min at room temperature (RT). Subsequently, DTT was removed by washing four times with 15 mL FL-Buffer and re-concentrating by centrifugation in Amicon® Ultra Centrifugal Filter units for 15 min at 4 °C. Concentration of the washed sample was determined before addition of the acceptor dye (ATTO 643) to a molar ratio of 0.7:1 (dye:protein). After light-protected incubation for 1.5 h at RT, the donor dye (Atto-532) was added to a molar ratio of 2:1 (dye:protein), followed by light-protected incubation over night at 5-10 °C. Unbound dye was removed by washing the sample seven times as described above. The last concentration step was continued until sample concentrations of approximately 5-40 µM were reached. Successful fluorescent labeling and the removal of unbound dye were confirmed by SDS-PAGE and in-gel fluorescence detection. LC-MS was performed to estimate the labeling efficiency as a ratio of the peak intensities for donor-acceptor(D/A)-labeled species over unlabeled, D/D-, A/A-, single- and triple-labeled species. Labeled protein samples were stored in 5 µL aliquots at −80 to −75 °C after freezing in liquid nitrogen.

#### Intact liquid chromatography mass spectrometry (LC-MS)

For verification and quantification by LC-MS, all purified, fluorescently labeled or modified proteins were diluted to 0.1 mg/mL in ddH_2_O and centrifuged for 5 min at 21000 xg, 4 °C. 2 µL per sample were desalted using a ProSwift™ RP-4H 1 ×50mm column (Thermo Fisher Scientific) followed by injection into a maXis II ETD ESI LCMS (Bruker Daltonics). Data was analyzed using DataAnalysis (Version 5.1, Bruker Daltonics).

#### (de)AMPylation activity assay

To analyze AMPylation activity, 10 µM BiP were incubated with 0.1 µM of monomeric labeled FicD mutants for 2 h at RT in 25 mM Hepes-KOH (pH 7.4), 100 mM KCl, 4 mM MgCl_2_, 1 mM CaCl_2_ and 1.5 mM ATP. The deAMPylation activity was probed by incubating 10 µM BiP_AMP_ with 0.1 µM of dimeric labeled FicD mutants in the same buffer without ATP. All reactions were stopped by the addition of Laemmli buffer (50 mM Tris-HCL, pH 6.8, 2 % (w/v) SDS, 10 % (v/v) glycerol, 100 mM DTT, 0.001 % (w/v) bromophenol blue) and boiling at 95 °C for 5 min. 500 ng of BiP were separated via SDS-PAGE on 10 % polyacrylamide gels (homemade) and transferred on Immobilon®-P PVDF membranes (Merck-Millipore) for 1.5 h at 230 mA using a Biometra Fastblot B44 (Analytik Jena) device. After blocking the membranes with Roti®-Block (Carl Roth) in Tris buffered saline with 0.1 % Tween20 (TBS-T) for at least 1 goat anti-mouse IgG (H+L) HRP conjugate (#31430, Thermo Fisher Scientific) at a ratio of 1:20,000 in TBS-T. The membrane was washed again as above and incubated for 5 min at RT with a 1:1:1:1 mixture of the SuperSignal West™ reagents Pico PLUS and Dura (Thermo Fisher Scientific) to develop the peroxidase signal. Chemiluminescence was detected with an Intas ECL Chemocam (Intas Science Imaging Instruments). For the subsequent detection of BiP, the membranes were stripped in Roti®Free Stripping-Buffer (Carl Roth) for 5 min at 60 °C. After six washing steps in TBS-T, the detection procedure described above was repeated using the rabbit-anti-GRP78 primary antibody for BiP (Thermo Scientific, #PA5-34941) at a ratio of 1:5000 in Roti®-Block in TBS-T and the goat-anti-rabbit IgG H&L (HRP) preadsorbed (#ab7090, abcam) at a ratio of 1:20,000 in TBS-T as a secondary antibody.

#### Accessible volume calculations

The accessible volume calculations for the FicD FRET constructs were done with the FRET-restrained positioning and screening (FPS) software^27^. For FicD:apo (PDB 6I7J), FicD-ATP (PDB 6I7K) and FicD-BiP (PDB 6ZMD) the donor fluorophore Atto532 is placed on R118, L104, K154, D138 and S170 residues in combination with the acceptor fluorophore Atto643 on the S288 residue. For the labeling dyes Atto532 and Atto643, the three radii AV model was used with a Förster radius of 59 Å, dye linker-length of 21 Å and linker width of 4.5 Å. The dye parameters used for AV simulations were R_532_(1): 5.5 Å, R_532_(2): 4.5 Å, R_532_(3): 1.5 Å for Atto532 and R_643_(1): 7.15 Å, R_643_(2): 4.5 Å, R_643_(3): 1.5 Å for Atto643.

#### smFRET measurements and data analysis

Fluorescently labeled FicD monomers labeled with Atto532 and Atto643, were diluted to a final concentration of 100 pM in the smFRET assay buffer containing 25 mM Hepes-KOH (pH 7.4), 100 mM KCl, 4 mM MgCl_2_, 1 mM CaCl_2_. For the experiments containing BiP, the labeled FicD mutants were incubated and measured in the presence of 100 µM unlabeled BiP concentration. The ATP concentration was kept at 5 mM for all the measurements containing the nucleotide. The dimeric FicD experiments were conducted by pre-incubating 1nM of double labeled and 50 nM unlabeled FicD dimers at 37°C for the formation of the dimeric protein containing a double labeled monomer.

A custom-built confocal microscope equipped with multiparameter fluorescence detection (MFD) and pulsed interleaved excitation (PIE)^34^ was used to perform the smFRET experiments as previously described^26^. The labeled FRET samples were excited with 532 nm and 640 nm laser lines with 70 and 25 µW of powers measured on the objecive, respectively. MFD-PIE allows for the determination of FRET efficiency, stoichiometry, fluorescence lifetime and anisotropy for each single-molecule burst simultaneously. Accurate FRET efficiencies (E) were determined using the formula:

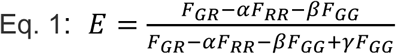

where F_GG_, F_GR_ and F_RR_ are the background-subtracted fluorescence signals detected in green/donor (GG), red/acceptor after donor excitation (GR) and acceptor channels (RR), respectively. Correction factors for detection efficiency (γ) spectral cross talk (α), direct excitation of the acceptor fluorophore (δ) were also used to compute the accurate FRET efficiencies.

Sub-millisecond dynamics of the FicD molecules were visualised by analysing the FRET efficiency versus donor fluorescence lifetime in the presence of acceptor plots. For a static system, there is a linear relationship between the FRET efficiency (E) and fluorescence lifetime of the donor in the presence of the acceptor (𝜏_𝐷(𝐴)_) according to the formula:

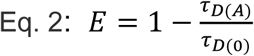

Where 𝜏_𝐷(0)_ is the fluorescence lifetime of donor-only species. The assessment of dynamics between states were based on the E versus plots where the theoretical static FRET line was defined according to the formula given in Eq.2 with a slight modification to account for linker dynamics at high-FRET efficiencies. If the protein switches between two states within the duration of the burst, the calculated FRET efficiency will be a species weighted value dependent on the respective FRET efficiencies^32^ and the time spent in the respective states, while the donor fluorescence lifetime will be determined using a photon-weighted average. Therefore, FRET efficiencies (E) for each state are calculated according to the formula:

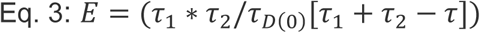

where is 𝜏 the photon weighted average of the donor lifetime, and 𝜏_1_and 𝜏_2_are the fluorescence lifetimes of the two different states. The higher number of photons collected from the lower-FRET efficiency state leads to a deviation towards the right of the static FRET-line and is indicative of sub-millisecond dynamics. The lifetime analysis of stoichiometry-filtered FRET populations for all the measured FicD constructs are given the supporting information file. All the analyzes of the collected data were done with the open-source PIE analysis with MATLAB (PAM) software^35^.

#### Fluorescence correlation spectroscopy

Fluorescence correlation spectroscopy (FCS) analysis was performed to obtain the diffusion coefficients of the labelled FicD molecules under different experimental conditions. The autocorrelation curves for the the acceptor channel after acceptor excitation were computed using the entire time-course of the smFRET measurements. The obtained autocorrelation functions (ACFs) were then fitted with the three-dimensional diffusion model function to obtain diffusion coefficients (𝐷) using the following equation:

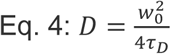

where 𝑤_0_ is the radial diameter of the confocal volume and 𝜏_𝐷_ is the diffusion time of the molecule.

#### Dynamic photon distribution analysis

A dynamic photon distribution analysis (PDA)^32^ was done to further assess the FRET states and extract the timescales of the sub-ms transitions between different conformations of the FicD constructs. PDA reveals the subpopulations and dynamics in a heterogeneous mixture of different states by investigating the photon statistics of the single molecule bursts that make up the width of the smFRET histograms. The PDA analysis was done by binning single-molecule FRET histograms to lengths of 0.5, 0.75 and 1.0 ms for the FicD constructs. The histograms of proximity ratio (PR) were computed from the raw photon counts of donor (S_D_) and FRET (S_F_) given in respective signal channels as described in Eq.5.

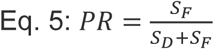

Due to the dynamically interconverting species, the PR histograms at different time-bins show changes. For the FicD^R118C^, dFicD^R118C^ and FicD^L104C^ constructs, a 2-state dynamic model was used for the dynamic PDA analysis. For the FicD^K154C^ construct which appears to be static on the sub-ms timescales in the absence of ATP, the PR histograms were fitted with a 3 state static model. Fluctuations between conformations were present for the measurements of FicD^K154C^ in the presence of ATP and they were fitted to a 2-state dynamic model. The 0.75-ms PR histograms of the dynamic PDA analysis and the summary of the values resulting from the global fits of all measured conditions are given in the Supporting information.

## Acknowledgement

Mass spectrometry was funded by the Deutsche Forschungsgemeinschaft (DFG, German Research Foundation – Projektnummer INST 152/859-1 FUGG). A.I. and D.C.L. acknowledges that support of the SFB 1035 project (German Research Foundation DFG, Sonderforschungsbereich 1035, Projektnummer 201302640) B05 (to AI) and A11 (to DL).

D.C.L. also acknowledges funding from the Federal Ministry of Education and Research (BMBF) and the Free State of Bavaria under the Excellence Strategy of the Federal Government and the Länder through the ONE MUNICH Project Munich Multiscale Biofabrication. We also acknowledge technical support from the SPC facility at EMBL Hamburg. A.I. acknowledges access to the core facilities and laboratories of the Centre for Structural Systems Biology (CSSB, Hamburg).

## Author contributions

S.R. generated all proteins, conducted biochemical and biophysical experiments, and analyzed the data. E.B. did single-molecule FRET experiments and performed fluorescence lifetime, FCS and photon distribution analyses. S.R., E.B., A.I. and D.C.L. interpreted the data and wrote the manuscript. All authors participated in manuscript editing and final approval.

## References

1. Rosenzweig, R., Nillegoda, N.B., Mayer, M.P. & Bukau, B. The Hsp70 chaperone network. Nat Rev Mol Cell Biol 20, 665–680 (2019).

2. Pobre, K.F.R., Poet, G.J. & Hendershot, L.M. The endoplasmic reticulum (ER) chaperone BiP is a master regulator of ER functions: Getting by with a little help from ERdj friends. J Biol Chem 294, 2098–2108 (2019).

3. Daugaard, M., Rohde, M. & Jaattela, M. The heat shock protein 70 family: Highly homologous proteins with overlapping and distinct functions. FEBS Lett 581, 3702–10 (2007).

4. Wieteska, L., Shahidi, S. & Zhuravleva, A. Allosteric fine-tuning of the conformational equilibrium poises the chaperone BiP for post-translational regulation. Elife 6(2017).

5. Yang, J., Nune, M., Zong, Y., Zhou, L. & Liu, Q. Close and Allosteric Opening of the Polypeptide-Binding Site in a Human Hsp70 Chaperone BiP. Structure 23, 2191–2203 (2015).

6. Voith von Voithenberg, L., et al. Comparative analysis of the coordinated motion of Hsp70s from different organelles observed by single-molecule three-color FRET. Proc Natl Acad Sci U S A 118(2021).

7. Gardner, B.M., Pincus, D., Gotthardt, K., Gallagher, C.M. & Walter, P. Endoplasmic reticulum stress sensing in the unfolded protein response. Cold Spring Harb Perspect Biol 5, a013169 (2013).

8. Faber, P.W. et al. Huntingtin Interacts with a Family of WW Domain Proteins. Human Molecular Genetics 7, 1463–1474 (1998).

9. Perera, L.A. & Ron, D. AMPylation and Endoplasmic Reticulum Protein Folding Homeostasis. Cold Spring Harb Perspect Biol 15(2023).

10. Worby, C.A. et al. The fic domain: regulation of cell signaling by adenylylation. Mol Cell 34, 93–103 (2009).

11. Ham, H. et al. Unfolded protein response-regulated Drosophila Fic (dFic) protein reversibly AMPylates BiP chaperone during endoplasmic reticulum homeostasis. J Biol Chem 289, 36059–69 (2014).

12. Preissler, S. et al. AMPylation matches BiP activity to client protein load in the endoplasmic reticulum. Elife 4, e12621 (2015).

13. Sanyal, A. et al. A novel link between Fic (filamentation induced by cAMP)-mediated adenylylation/AMPylation and the unfolded protein response. J Biol Chem 290, 8482–99 (2015).

14. Gulen, B. & Itzen, A. Revisiting AMPylation through the lens of Fic enzymes. Trends Microbiol 30, 350–363 (2022).

15. Preissler, S. et al. AMPylation targets the rate-limiting step of BiP’s ATPase cycle for its functional inactivation. Elife 6(2017).

16. Preissler, S., Rato, C., Perera, L., Saudek, V. & Ron, D. FICD acts bifunctionally to AMPylate and de-AMPylate the endoplasmic reticulum chaperone BiP. Nat Struct Mol Biol 24, 23–29 (2017).

17. Casey, A.K. et al. Fic-mediated AMPylation tempers the unfolded protein response during physiological stress. Proc Natl Acad Sci U S A 119, e2208317119 (2022).

18. Perera, L.A. et al. Infancy-onset diabetes caused by de-regulated AMPylation of the human endoplasmic reticulum chaperone BiP. EMBO Mol Med 15, e16491 (2023).

19. Rebelo, A.P. et al. BiP inactivation due to loss of the deAMPylation function of FICD causes a motor neuron disease. Genet Med 24, 2487–2500 (2022).

20. Bunney, T.D. et al. Crystal structure of the human, FIC-domain containing protein HYPE and implications for its functions. Structure 22, 1831–1843 (2014).

21. Perera, L.A. et al. An oligomeric state-dependent switch in the ER enzyme FICD regulates AMPylation and deAMPylation of BiP. EMBO J 38, e102177 (2019).

22. Engel, P. et al. Adenylylation control by intra- or intermolecular active-site obstruction in Fic proteins. Nature 482, 107–10 (2012).

23. Xiao, J., Worby, C.A., Mattoo, S., Sankaran, B. & Dixon, J.E. Structural basis of Fic-mediated adenylylation. Nat Struct Mol Biol 17, 1004–10 (2010).

24. Fauser, J. et al. Specificity of AMPylation of the human chaperone BiP is mediated by TPR motifs of FICD. Nat Commun 12, 2426 (2021).

25. Perera, L.A. et al. Structures of a deAMPylation complex rationalise the switch between antagonistic catalytic activities of FICD. Nat Commun 12, 5004 (2021).

26. Kudryavtsev, V. et al. Combining MFD and PIE for accurate single-pair Forster resonance energy transfer measurements. Chemphyschem 13, 1060–78 (2012).

27. Kalinin, S. et al. A toolkit and benchmark study for FRET-restrained high-precision structural modeling. Nature Methods 9, 1218–1225 (2012).

28. Opanasyuk, O. et al. Unraveling multi-state molecular dynamics in single-molecule FRET experiments. II. Quantitative analysis of multi-state kinetic networks. J Chem Phys 157, 031501 (2022).

29. Barth, A. et al. Unraveling multi-state molecular dynamics in single-molecule FRET experiments. I. Theory of FRET-lines. J Chem Phys 156, 141501 (2022).

30. Wei, J., Gaut, J.R. & Hendershot, L.M. In vitro dissociation of BiP-peptide complexes requires a conformational change in BiP after ATP binding but does not require ATP hydrolysis. J Biol Chem 270, 26677–82 (1995).

31. Sanyal, A. et al. Kinetic and structural parameters governing Fic-mediated adenylylation/AMPylation of the Hsp70 chaperone, BiP/GRP78. Cell Stress Chaperones 26, 639–656 (2021).

32. Kalinin, S., Valeri, A., Antonik, M., Felekyan, S. & Seidel, C.A.M. Detection of Structural Dynamics by FRET: A Photon Distribution and Fluorescence Lifetime Analysis of Systems with Multiple States. The Journal of Physical Chemistry B 114, 7983–7995 (2010).

33. Jeong, J.Y. et al. One-step sequence- and ligation-independent cloning as a rapid and versatile cloning method for functional genomics studies. Appl Environ Microbiol 78, 5440–3 (2012).

34. Muller, B.K., Zaychikov, E., Brauchle, C. & Lamb, D.C. Pulsed interleaved excitation. Biophys J 89, 3508–22 (2005).

35. Schrimpf, W., Barth, A., Hendrix, J. & Lamb, D.C. PAM: A Framework for Integrated Analysis of Imaging, Single-Molecule, and Ensemble Fluorescence Data. Biophys J 114, 1518–1528 (2018).

