## Supplementary Information for "Resolving the Dynamic Interplay of the Conformational States in the TPR Domain of Human AMPylase FicD"

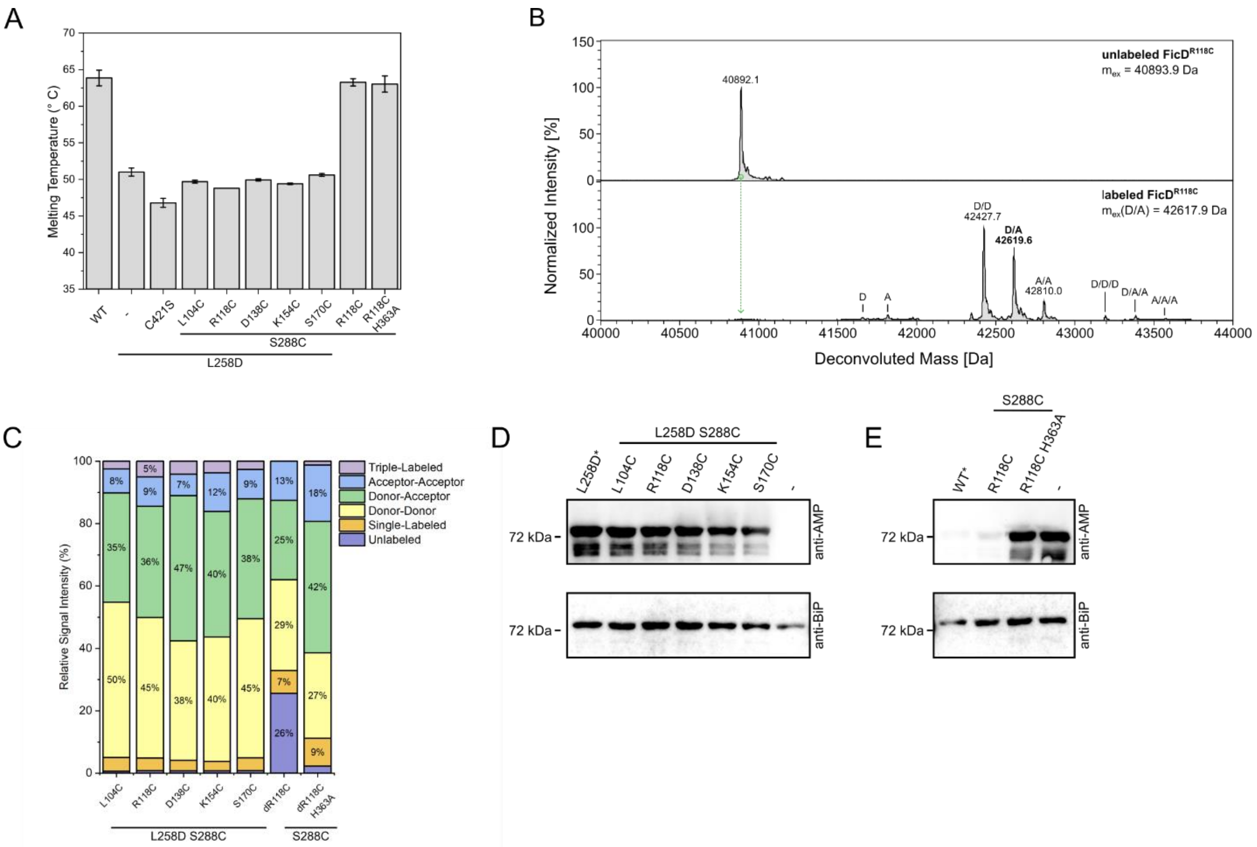

**Figure S1: Validation of the labeling efficiency and catalytic activity of the smFRET labeled samples. A:** Melting temperatures as determined by nanoDSF of unlabeled FicD mutants. Error bars represent 3-5 technical replicates per mutant.**B:** Exemplary deconvoluted mass spectra of monomeric FicD$^{R118C}$ before (top panel) and after (bottom panel) stochastic labeling with ATTO532 and ATTO643. Indicated are the expected masses ($m_{ex}$) for unlabeled and hetero, double-labeled (DA) FicD. Annotated are the species that are labeled with the donor-dye (D), acceptor-dye (A) or combinations thereof, with the measured masses given in Dalten for the main peaks. The mass shifts for a single, covalently linked ATTO532 or ATTO643 molecule are 766.8 Da and 957.2 Da, respectively (ATTO-TEC). **C:** Relative signal intensities for the differently labeled FicD-species as indicated in A for all smFRET samples included in this study. **D:** AMPylation Activity of monomeric and fluorescently labeled FicD mutants was analyzed by incubation of 0.1 µM FicD with 10 µM BiP$^{T229A}$ and 1.5 mM ATP. Monomeric, unlabeled FicD$^{L258D}$ (asterisk) served as a positive control. **E:** deAMPylation activity of dimeric and fluorescently labeled FicD mutants was analyzed by incubation of 0.1 µM FicD with 10 µM BiP-AMP$^{T229A}$ and 1.5 mM ATP. Dimeric, unlabeled FicD WT (asterisk) served as a positive control.

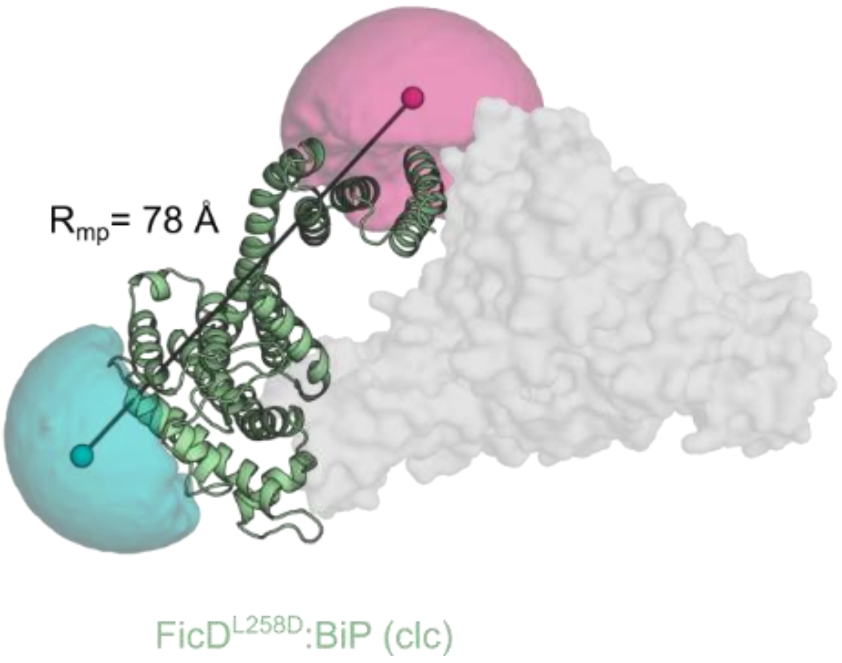

**Figure S2: Accessible volume (AV) calculation of FicD in complex with BiP.** The magenta (donor) and teal (acceptor) surfaces represent the possible spatial distribution of the fluorophores, the spheres indicate the mean positions of the donor and acceptor dyes. The black line indicates the distance between both mean positions ($R_{mp}$). The calculation and representation are based on PDB 6ZMD.

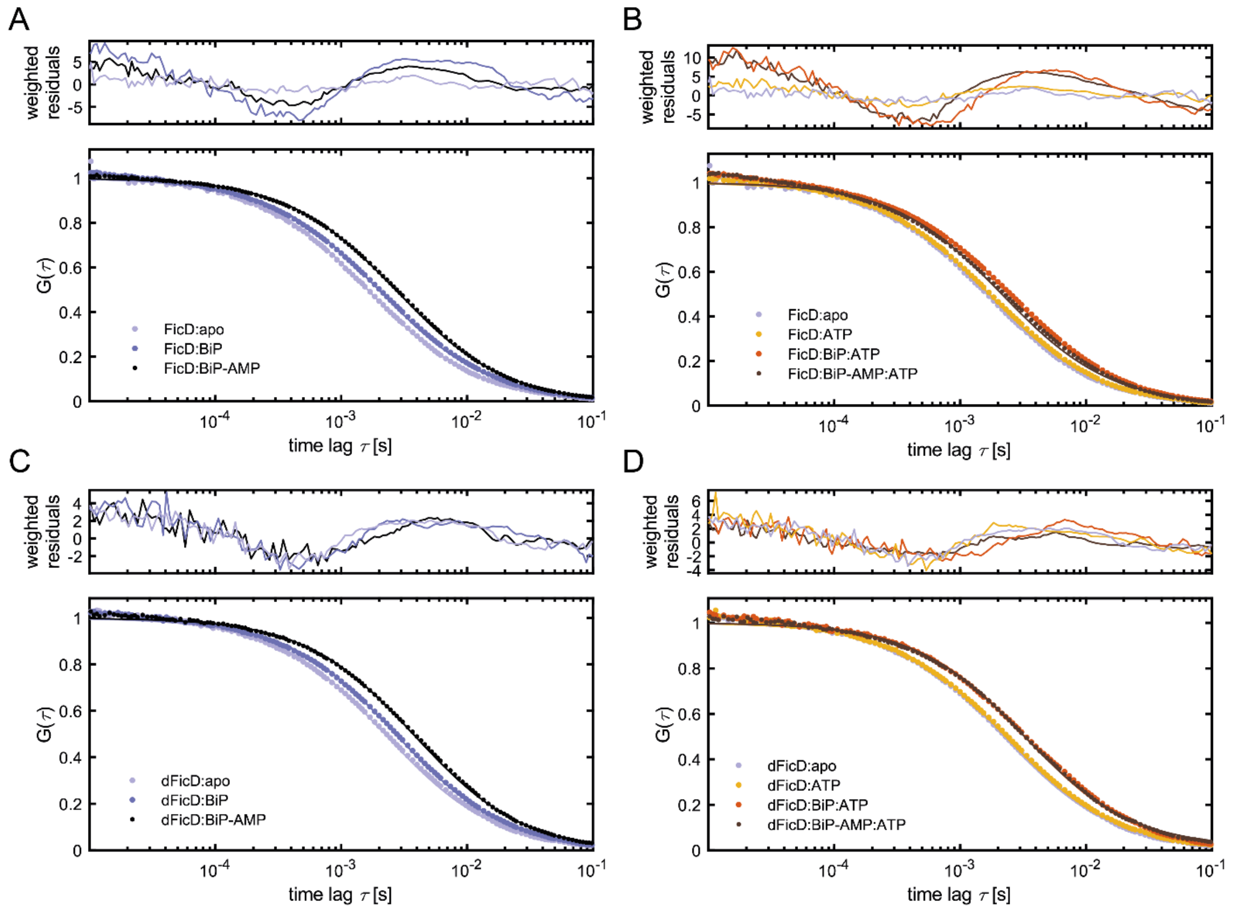

**Figure S3: Fluorescence cross-correlation spectroscopy (FCS) analysis of the monomeric and dimeric FicD$^{R118C}$.** FCS analysis was performed on monomeric or dimeric FicD$^{R118}$ FRET constructs labelled with Atto532 and Atto643 and the auto-correlation function (ACF) of acceptor channel, under the various experimental conditions are given in plots **A-D**. **A:** ACF curves of the monomeric FicD$^{R118C}$:apo (light blue) in comparison to FicD in the presence of BiP (blue) and BiP-AMP (black). **B:** ACF curves of the monomeric FicD$^{R118C}$:apo (light blue) in comparison to the FicD$^{R118C}$:ATP (yellow) in the presence of BiP (orange) and BiP-AMP (brown). **C:** ACF curves of the dimeric FicD$^{R118C}$:apo (light blue) in comparison to FicD in the presence of BiP (blue) and BiP-AMP (black). **D:** ACF curves of the dimeric FicD$^{R118C}$:apo (light blue) in comparison to the FicD$^{R118C}$:ATP (yellow) in the presence of BiP (orange) and BiP-AMP (brown). The resulting diffusion coefficient values from the fits are listed in **Table S4**.

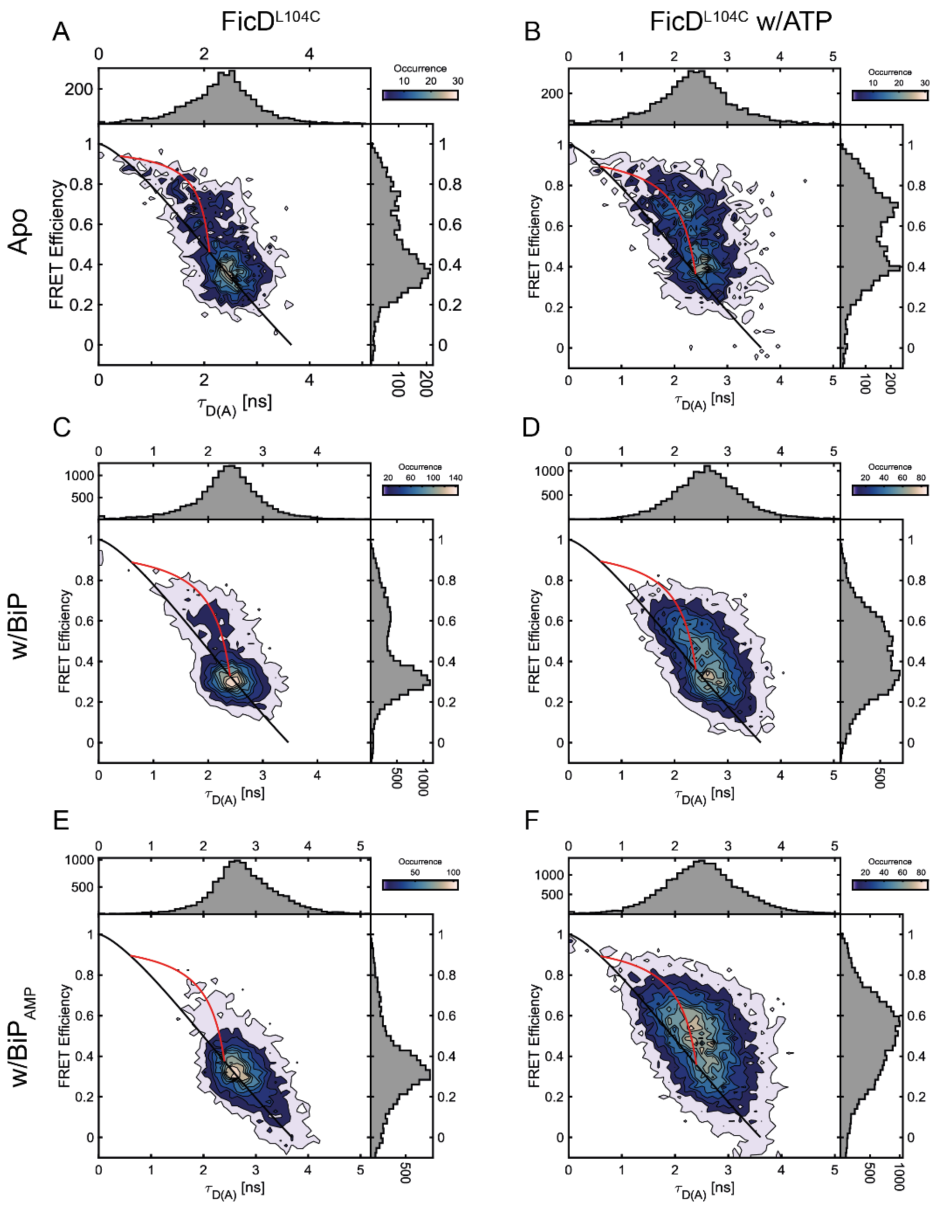

**Figure S4: 2D plots of FRET efficiency (E) vs. donor fluorescence lifetime in the presence of an acceptor ($\tau_{D(A)}$) for the monomeric FicD$^{L104C}$.** The 2D E-$\tau_{D(A)}$ plots are given for **A:** FicD$^{L104C}$ apo, FicD$^{L104C}$ in the presence of **B:** ATP, **C:** BiP, **D:** BiP and ATP, **E:** BiP$_{AMP}$ and **F:** BiP$_{AMP}$ and ATP. Populations of static molecules are described by the polynomial static FRET line (Eq. 2; black line). The molecules undergoing microsecond to millisecond conformational dynamics can be described theoretically using the dynamic FRET line (Eq. 3; red line). The start and end points were extracted from fluorescence lifetime analysis of all molecules using a biexponential model function (**Table S5**).

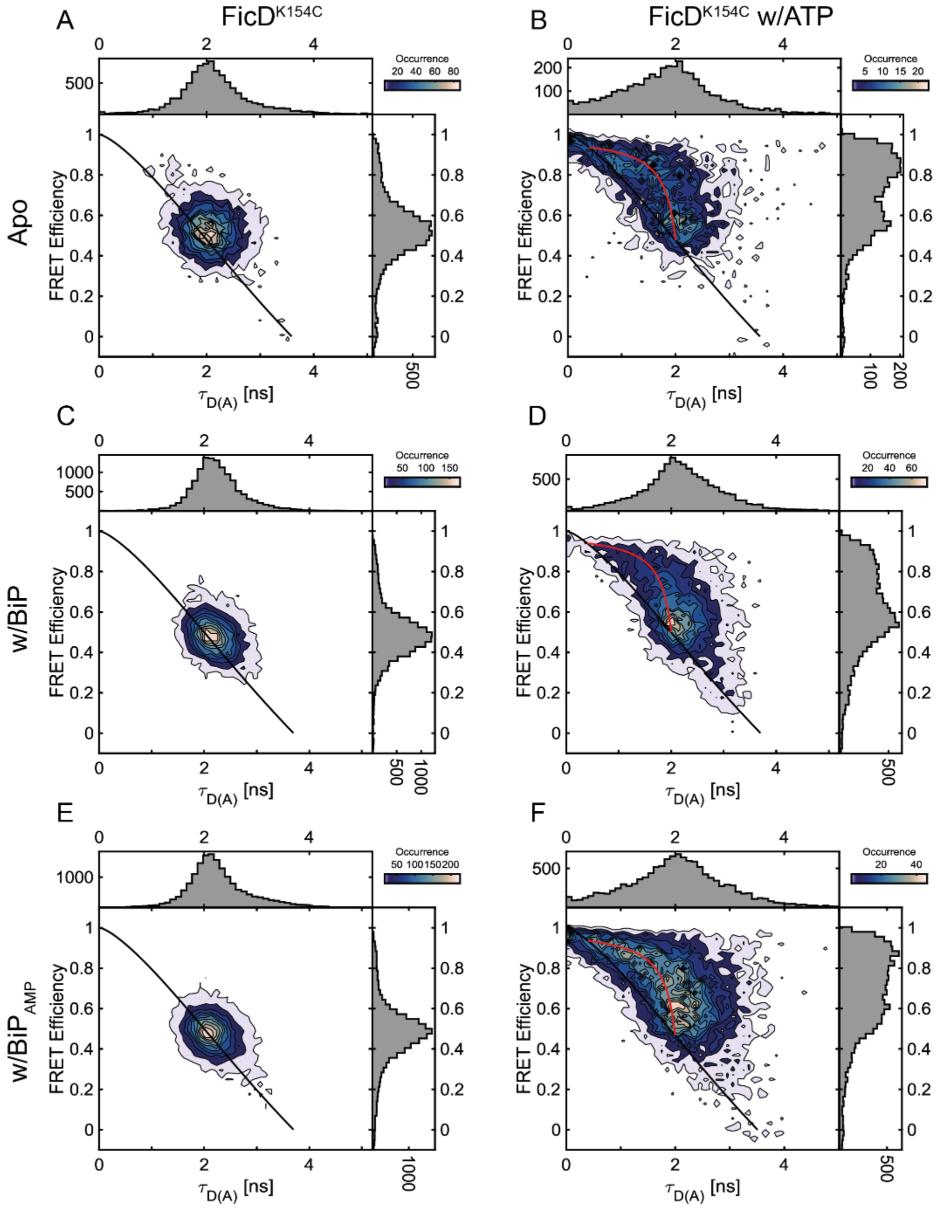

**Figure S5: 2D plots of FRET efficiency (E) vs. donor fluorescence lifetime in the presence of an acceptor ($\tau_{D(A)}$) for the monomeric FicD$^{K154C}$.** The 2D E-$\tau_{D(A)}$ plots are given for **A:** FicD$^{K154C}$ apo, FicD$^{K154C}$ in the presence of **B:** ATP, **C:** BiP, **D:** BiP and ATP, **E:** BiP$_{AMP}$ and **F:** BiP$_{AMP}$ and ATP. Populations of static molecules are described by the polynomial static FRET line (Eq. 2; black line). The molecules undergoing microsecond to millisecond conformational dynamics can be described theoretically using the dynamic FRET line (Eq. 3; red line). The start and end points were extracted from fluorescence lifetime analysis of all molecules using a biexponential model function (**Table S7**).

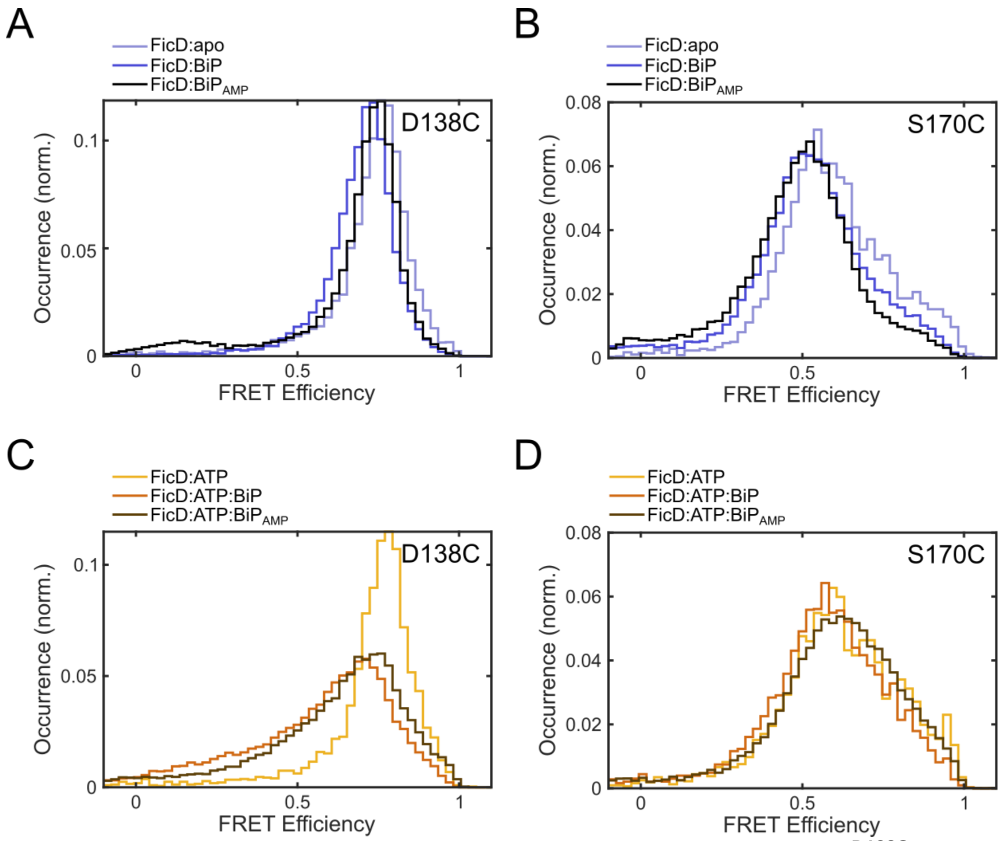

**Figure S6: FRET efficiency histograms of the monomeric FicD$^{D138C}$ and FicD$^{S170C}$ constructs. A:** SmFRET efficiency histograms of FicD$^{D138C}$:apo (light blue) in comparison to FicD$^{D138C}$ in the presence of BiP (blue) and BiP$_{AMP}$ (black). **B:** SmFRET efficiency histograms of FicD$^{S170C}$:apo (light blue) in comparison to FicD$^{S170C}$ in the presence of BiP (blue) and BiP$_{AMP}$ (black). **C:** SmFRET efficiency histograms of FicD$^{D138C}$:ATP (yellow) in comparison to FicD$^{D138CC}$:ATP in the presence of BiP (orange) and BiP$_{AMP}$ (brown). **D:** SmFRET efficiency histograms of FicD$^{S170C}$:ATP (yellow) in comparison to FicD$^{S170C}$:ATP in the presence of BiP (orange) and BiP$_{AMP}$ (brown).

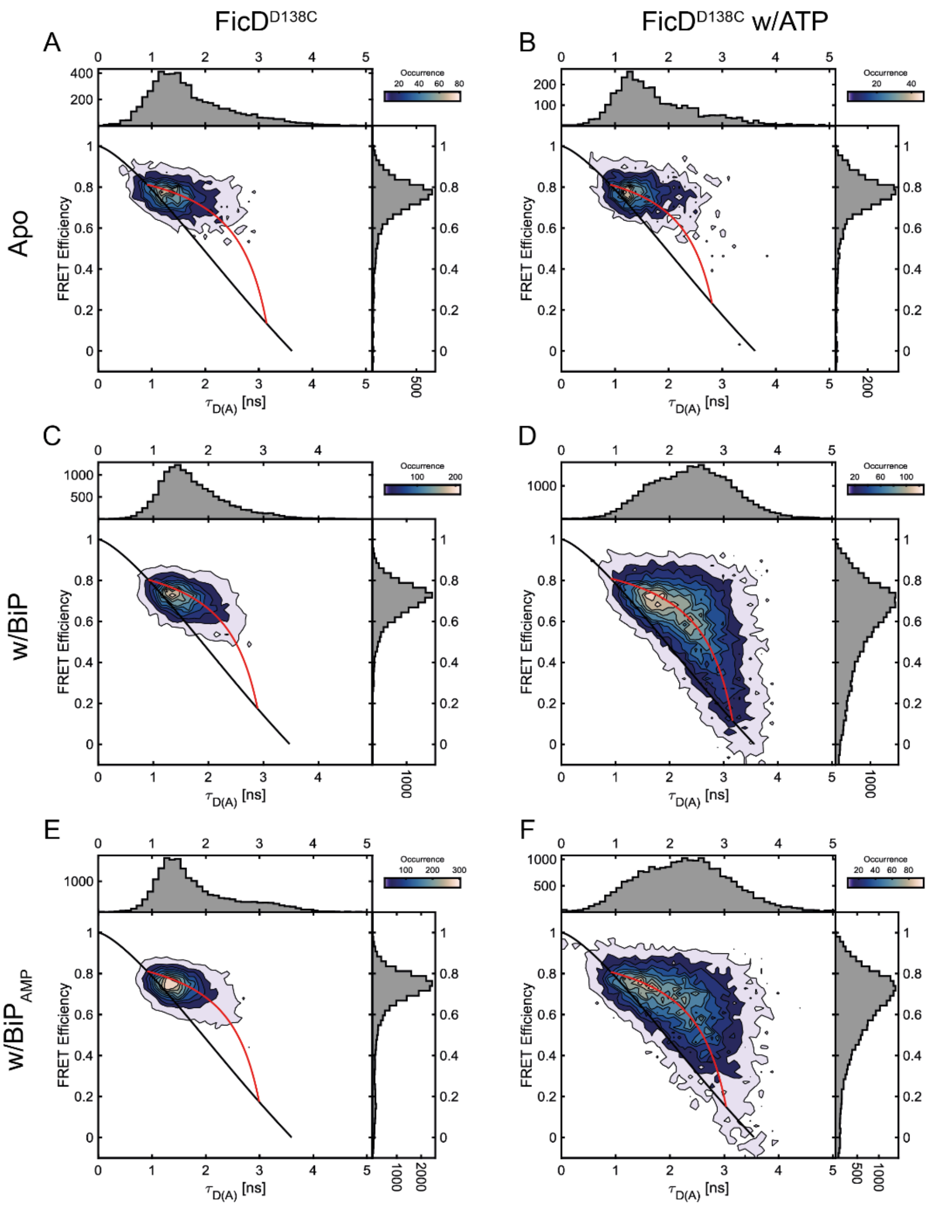

**Figure S7: 2D plots of FRET efficiency (E) vs. donor fluorescence lifetime in the presence of an acceptor ($\tau_{D(A)}$) for the monomeric FicD$^{D138C}$.** The 2D E-$\tau_{D(A)}$ plots are given for **A:** FicD$^{D138C}$:apo and FicD$^{D138C}$ in the presence of **B:** ATP, **C:** BiP, **D:** BiP and ATP, **E:** BiP$_{AMP}$ and **F:** BiP$_{AMP}$ and ATP. Populations of static molecules are described by the polynomial static FRET line (Eq. 2; black line). The molecules undergoing microsecond to millisecond conformational dynamics can be described theoretically using the dynamic FRET line (Eq. 3; red line). The start and end points were extracted from fluorescence lifetime analysis of all molecules using a biexponential model function (**Table S9**).

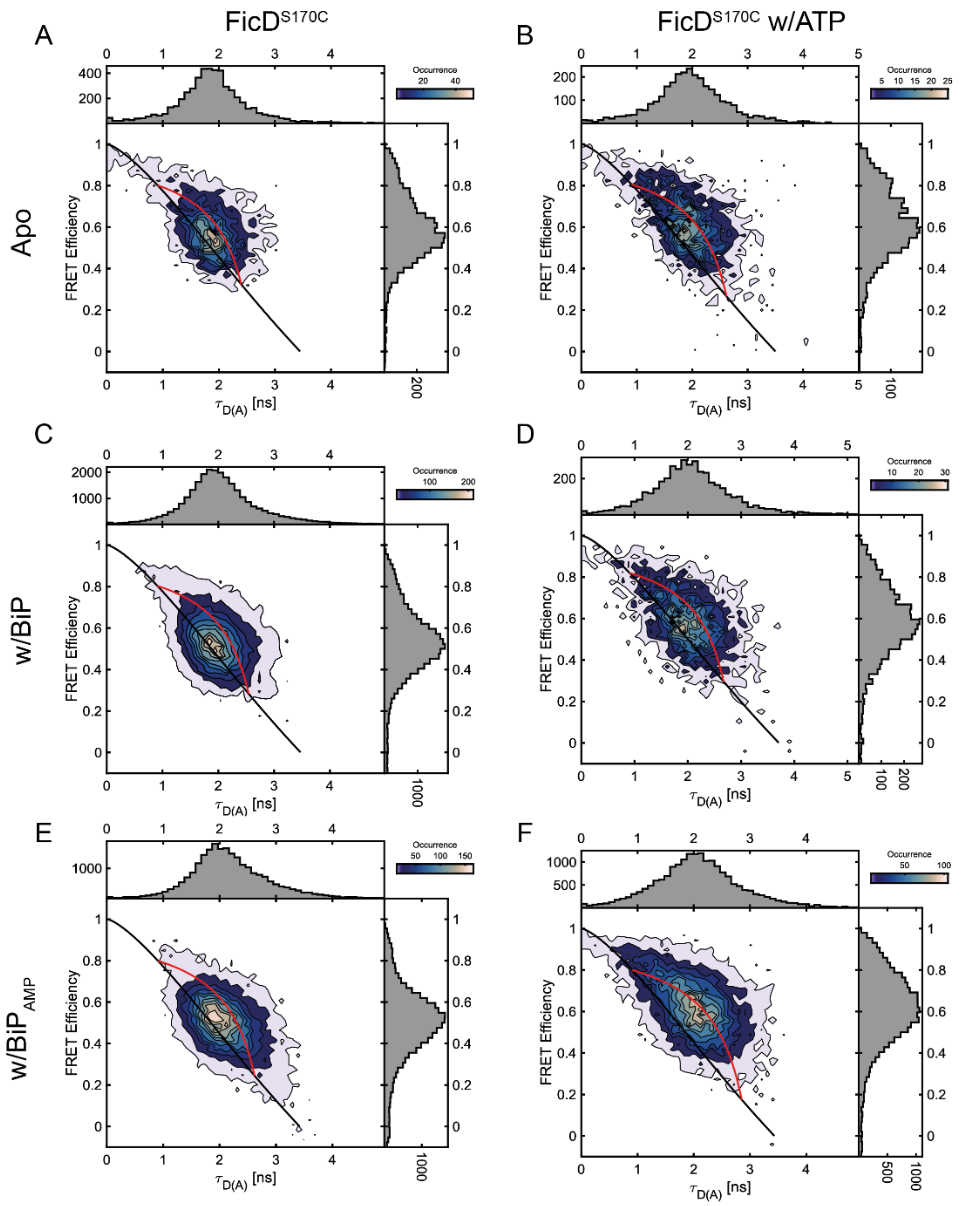

**Figure S8: 2D plots of FRET efficiency (E) vs. donor fluorescence lifetime in the presence of an acceptor ($\tau_{D(A)}$) for the monomeric FicD$^{S170C}$.** The 2D E-$\tau_{D(A)}$ plots are given for **A:** FicD$^{S170C}$:apo and FicD$^{S170C}$ in the presence of **B:** ATP, **C:** BiP, **D:** BiP and ATP, **E:** BiP$_{AMP}$ and **F:** BiP$_{AMP}$ and ATP. Populations of static molecules are described by the polynomial static FRET line (Eq. 2; black line). The molecules undergoing microsecond to millisecond conformational dynamics can be described theoretically using the dynamic FRET line (Eq. 3; red line). The start and end points were extracted from fluorescence lifetime analysis of all molecules using a biexponential model function (**Table S10**).

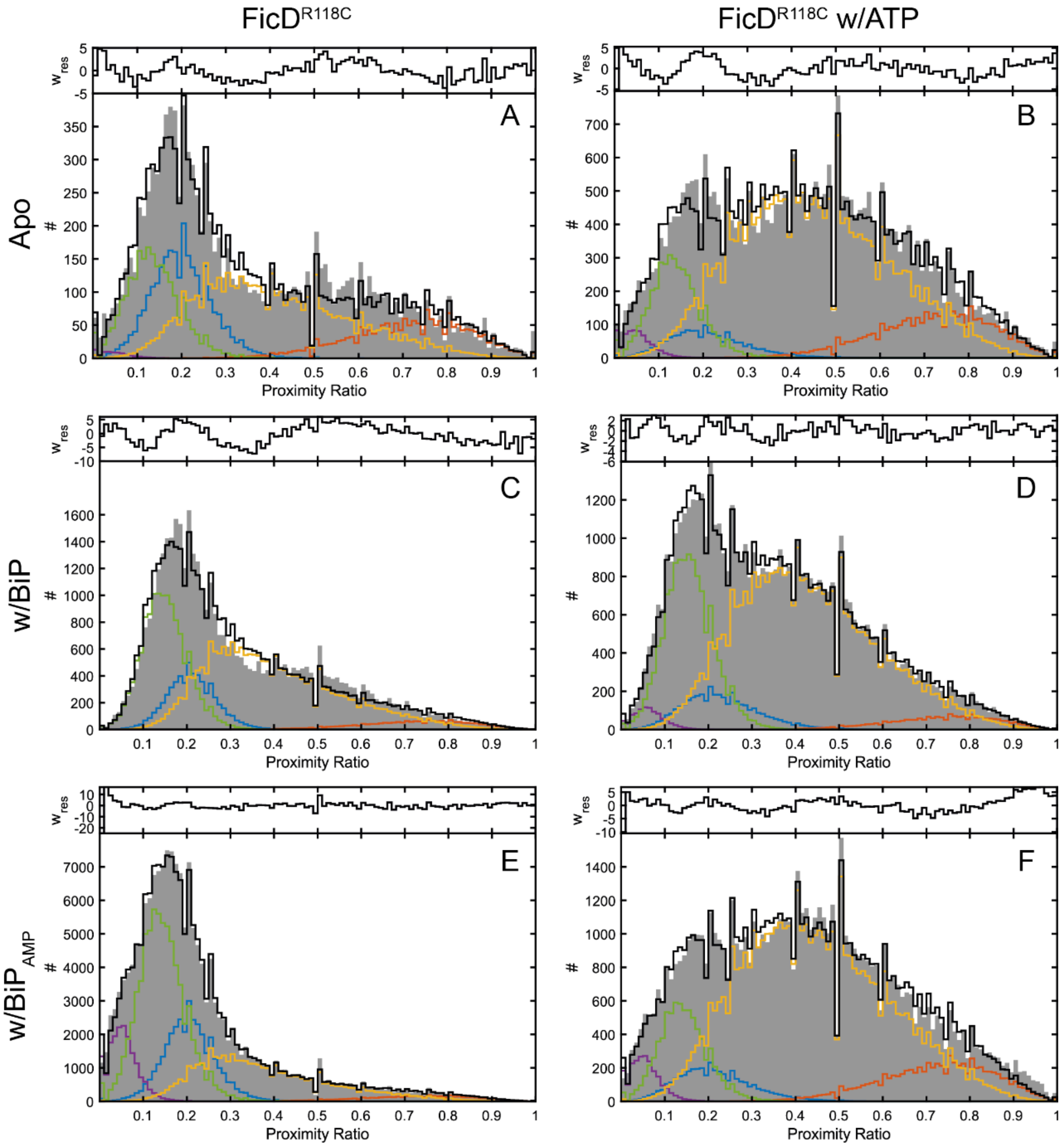

**Figure S9: Analysis of the conformational states and dynamics of the monomeric FicD$^{R118C}$ with photon distribution analysis (PDA).** For dynamic PDA, uncorrected FRET efficiency (proximity ratio) histograms with time binnings of 0.5 ms, 0.75 ms and 1.0 ms were used to calculate the distance distribution for different FRET populations and dynamics between them. The data for the time window size of 0.75 ms is shown for all the conditions. Proximity ratio histograms of the monomeric FicD$^{R118C}$ are given for **A:** apo, in the presence of **B:** ATP, **C:** BiP, **D:** BiP and ATP, **E:** $BiP_{AMP}$ and **F:** $BiP_{AMP}$ and ATP. The histograms for all measurement conditions were fitted using a 2-state dynamic PDA model. TPR-in and -out conformations are given in blue and orange and the dynamic interconversion between them are shown in yellow. An additional static low-FRET (TPR-in, green) and a static, low FRET state ($A_{R1}$, purple) for impurities were included in the PDA fits. Values for the fits are given in **Table S11**.

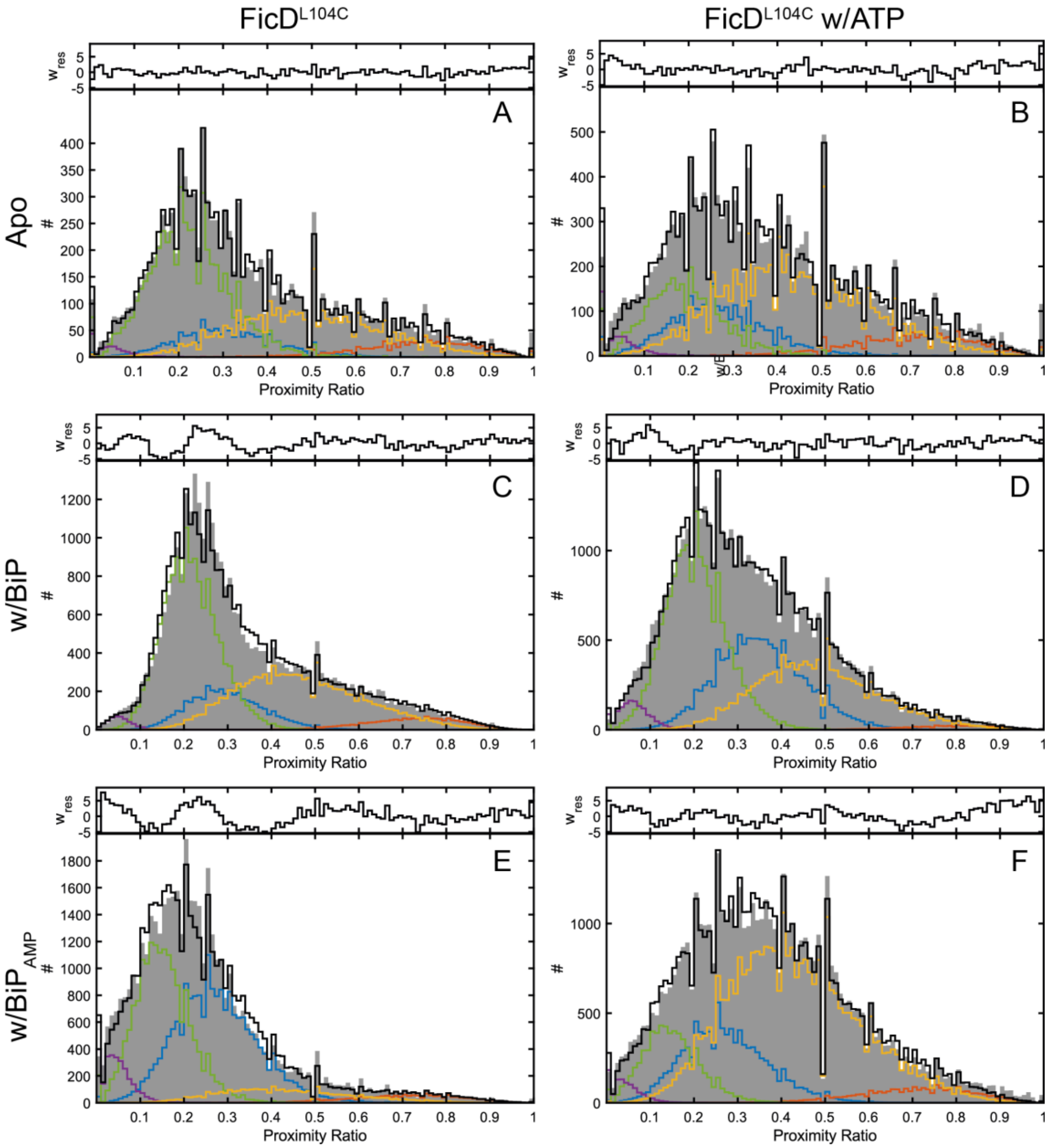

**Figure S10: Analysis of the conformational states and dynamics of the monomeric FicD$^{L104C}$ with photon distribution analysis (PDA).** For dynamic PDA, uncorrected FRET efficiency (proximity ratio) histograms with time binnings of 0.5 ms, 0.75 ms and 1.0 ms were used to calculate the distance distribution for different FRET populations and dynamics between them. The data for the time window size of 0.75 ms is shown for all the conditions. Proximity ratio histograms of the monomeric FicD$^{L104C}$ are given for **A:** apo, in the presence of **B:** ATP, **C:** BiP, **D:** BiP and ATP, **E:** BiP$_{AMP}$ and **F:** BiP$_{AMP}$ and ATP. The histograms for all conditions were fitted using a 2-state dynamic PDA model. TPR-in and -out conformations are given in blue and orange and the dynamic interconversion between them are shown in yellow. An additional static low-FRET (TPR-in, green) and a static, low FRET state ($A_{R1}$, purple) for impurities were included in the PDA fits. Values for the fits are given in **Table S12**.

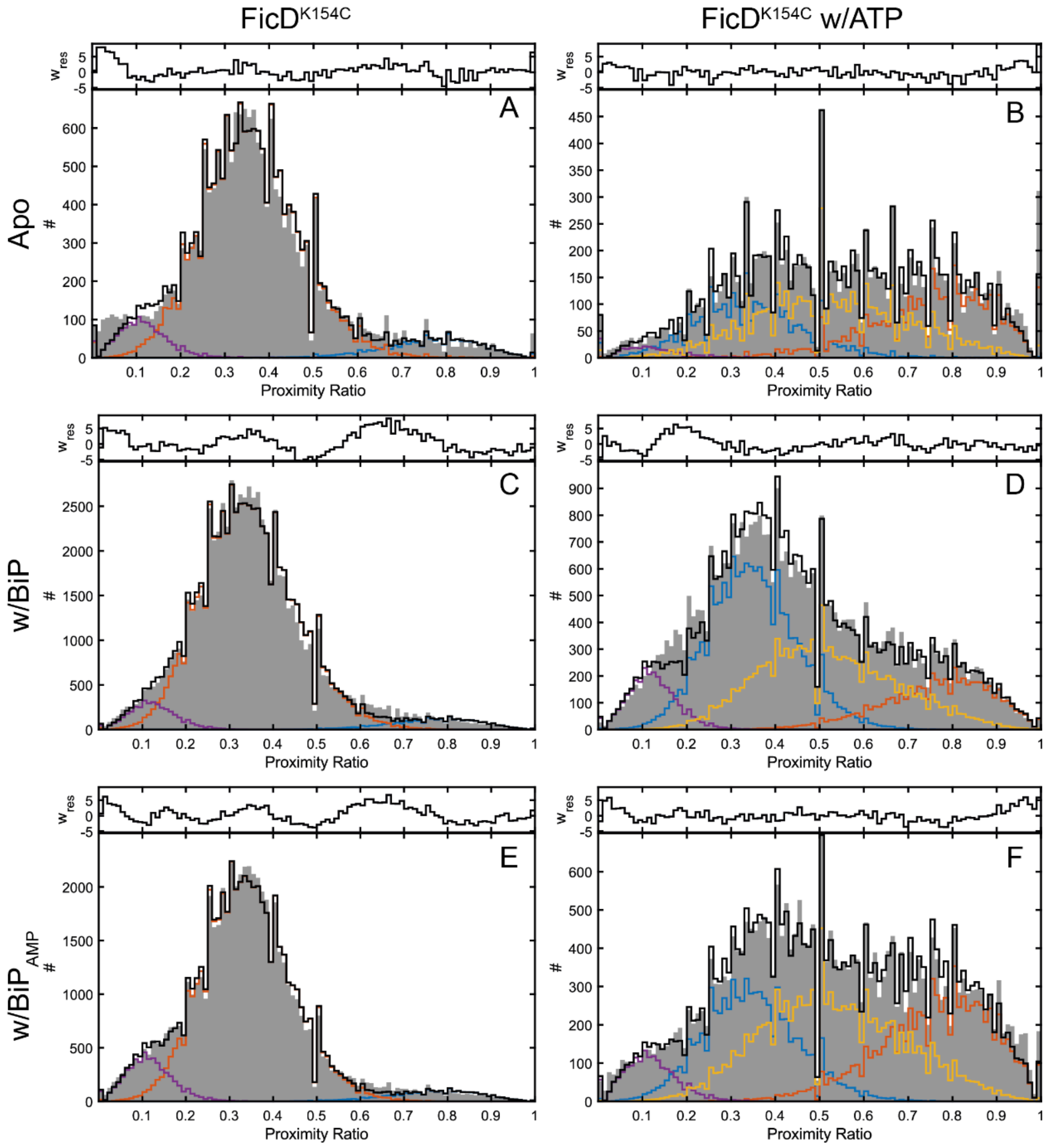

**Figure S11: Analysis of the conformational states and dynamics of the monomerc FicD$^{K154C}$ with photon distribution analysis (PDA).** For both static and dynamic PDA, uncorrected FRET efficiency (proximity ratio) histograms with time binnings of 0.5 ms, 0.75 ms and 1.0 ms were used to calculate the distance distribution for different FRET populations and dynamics between them. The data for the time window size of 0.75 ms is shown for all the conditions. Proximity ratio histograms of the monomeric FicD$^{K154C}$ **A:** apo and in the presence of **C:** BiP and **E:** BiP-AMP are fitted using a 3-state static PDA fit model. TPR-in and -out conformations are given in orange and blue. A second static low-FRET (TPR-in, purple) state is included in the PDA fits. Proximity ratio histograms of FicD$^{K154C}$ in the presence of **B:** ATP, **D:** BiP+ATP and F**:** BiP-AMP+ATP were fitted using the 2-state dynamic PDA model. TPR-in and -out conformations are given in orange and blue and the dynamic interconversion between them are shown in yellow. An additional static low-FRET (TPR-in, purple) state is included in the PDA fits. Values for the fits are given in **Table S13**.

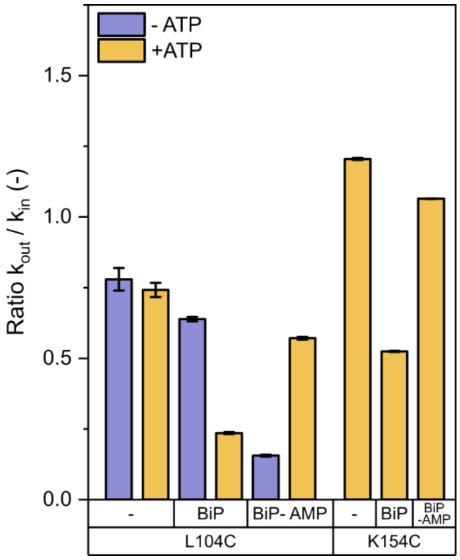

**Figure S12: Comparison of the $k_{out}/k_{in}$ ratios derived from the PDA analysis for FicD$^{L104C}$ and FicD$^{K154C}$.** The $k_{out}$ and $k_{in}$ values for FicD$^{L104C}$ and FicD$^{K154C}$ are given in **Tables S12** and **S13**, respectively.

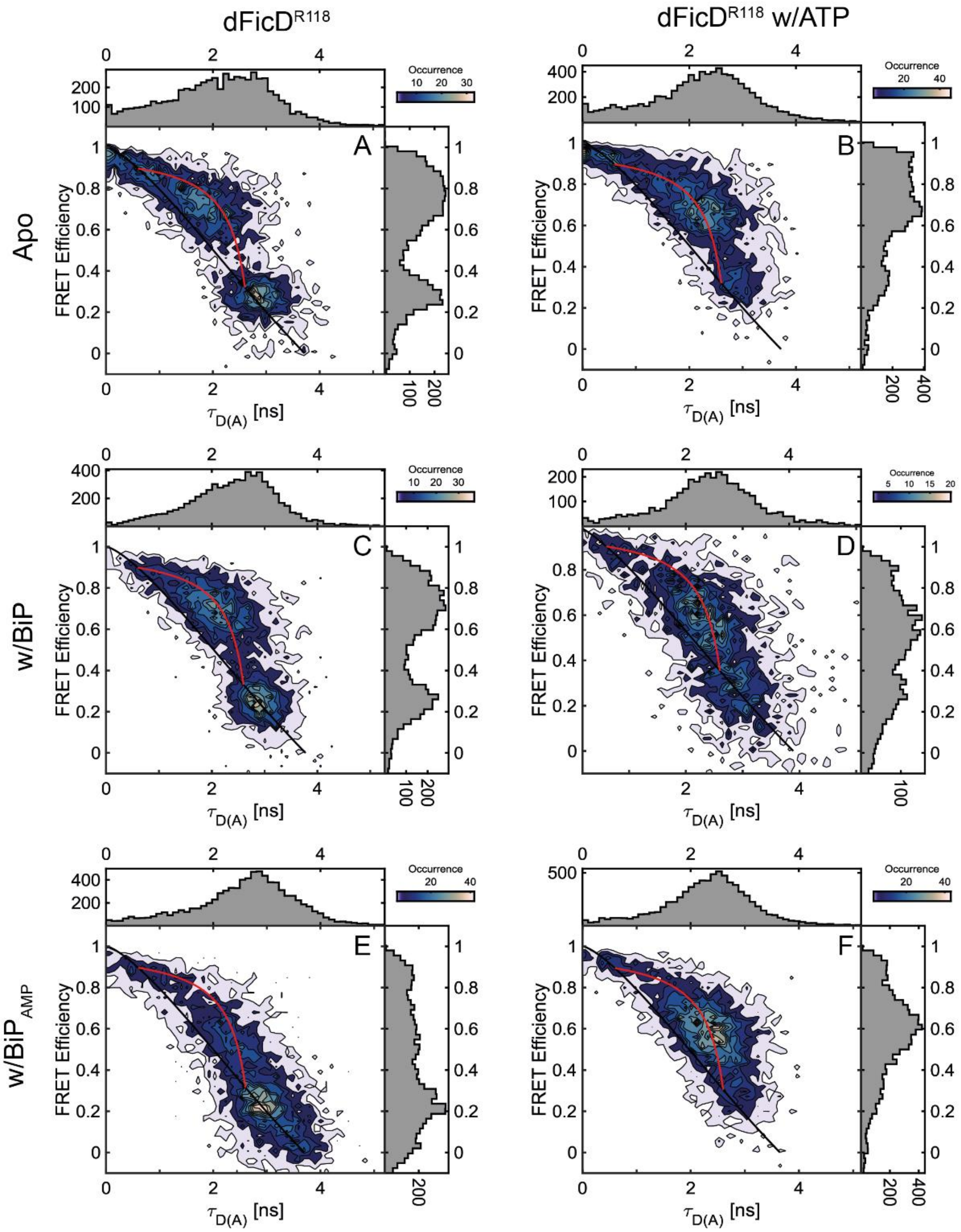

**Figure S13: 2D plots of FRET efficiency (E) vs. donor fluorescence lifetime in the presence of an acceptor ($\tau_{D(A)}$) for the dimeric FicD$^{R118C}$.** The 2D E-$\tau_{D(A)}$ plots are given for **A:** dFicD$^{R118C}$ apo, dFicD$^{R118C}$ in the presence of **B:** ATP, **C:** BiP, **D:** BiP and ATP, **E:** BiP$_{AMP}$ and **F:** BiP$_{AMP}$ and ATP. Populations of static molecules are described by the polynomial static FRET line (Eq. 2; black line). The molecules undergoing microsecond to millisecond conformational dynamics can be described theoretically using the dynamic FRET line (Eq. 3; red line). The start and end points were extracted from fluorescence lifetime analysis of all molecules using a biexponential model function (**Table S14**).

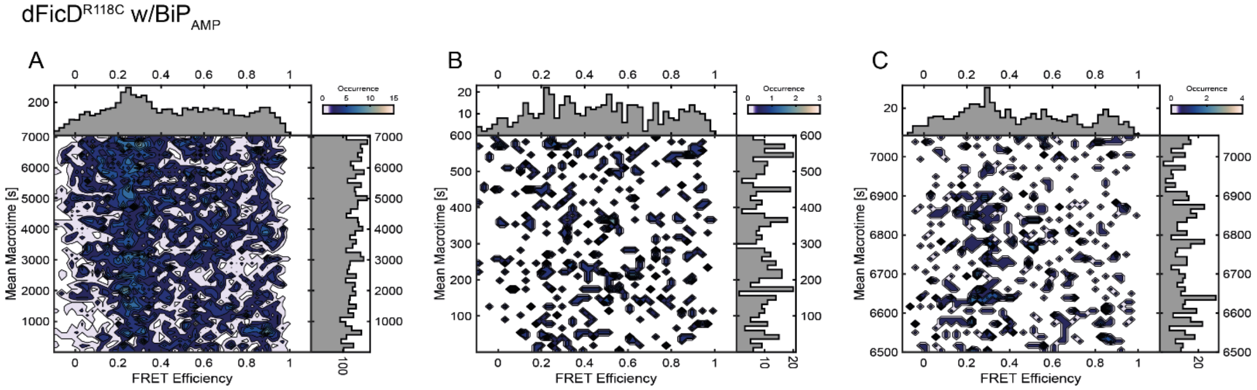

**Figure S14: 2D mean macrotime versus FRET efficiency plots of the dimeric FicD$^{R118C}$ measurement in the presence of BiP$_{AMP}$. A:** 2D mean macrotime verus FRET efficiency plots of the dimeric FicD$^{R118C}$ measurement in the presence of BiP$_{AMP}$ for the 2-hour time interval. **B:** 2D mean macrotime versus FRET efficiency plots of the dimeric FicD$^{R118C}$ measurement in the presence of BiP$_{AMP}$ for the first 10 minutes and **C:** the last 10 minutes of the same measurement.

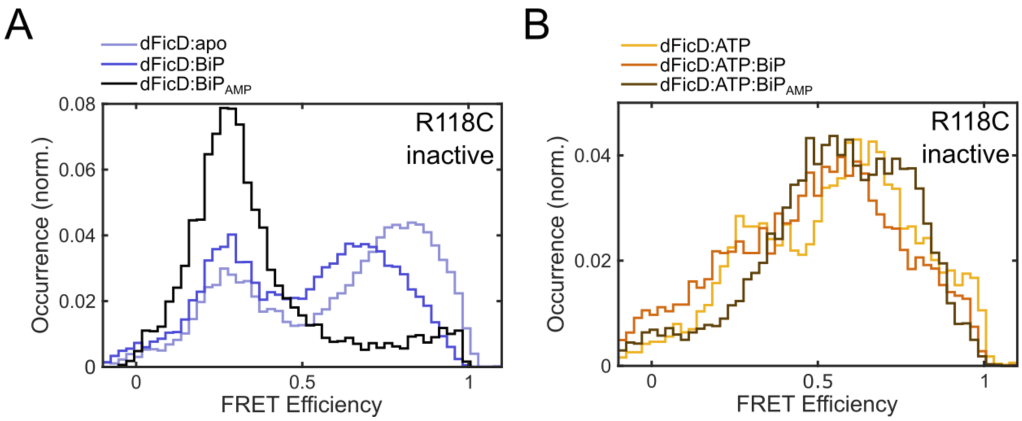

**Figure S15: FRET efficiency histograms of the dimeric, inactive FicD$^{R118C}$ construct. A:** SmFRET efficiency histograms of the dimeric, inactive FicD$^{R118C}$:apo (light blue) in comparison to in the presence of BiP (blue) and BiP$_{AMP}$ (black). **B:** SmFRET efficiency histograms of the dimeric inactive FicD$^{R118C}$:ATP (yellow) in comparison to in the presence of BiP (orange) and BiP$_{AMP}$ (brown).

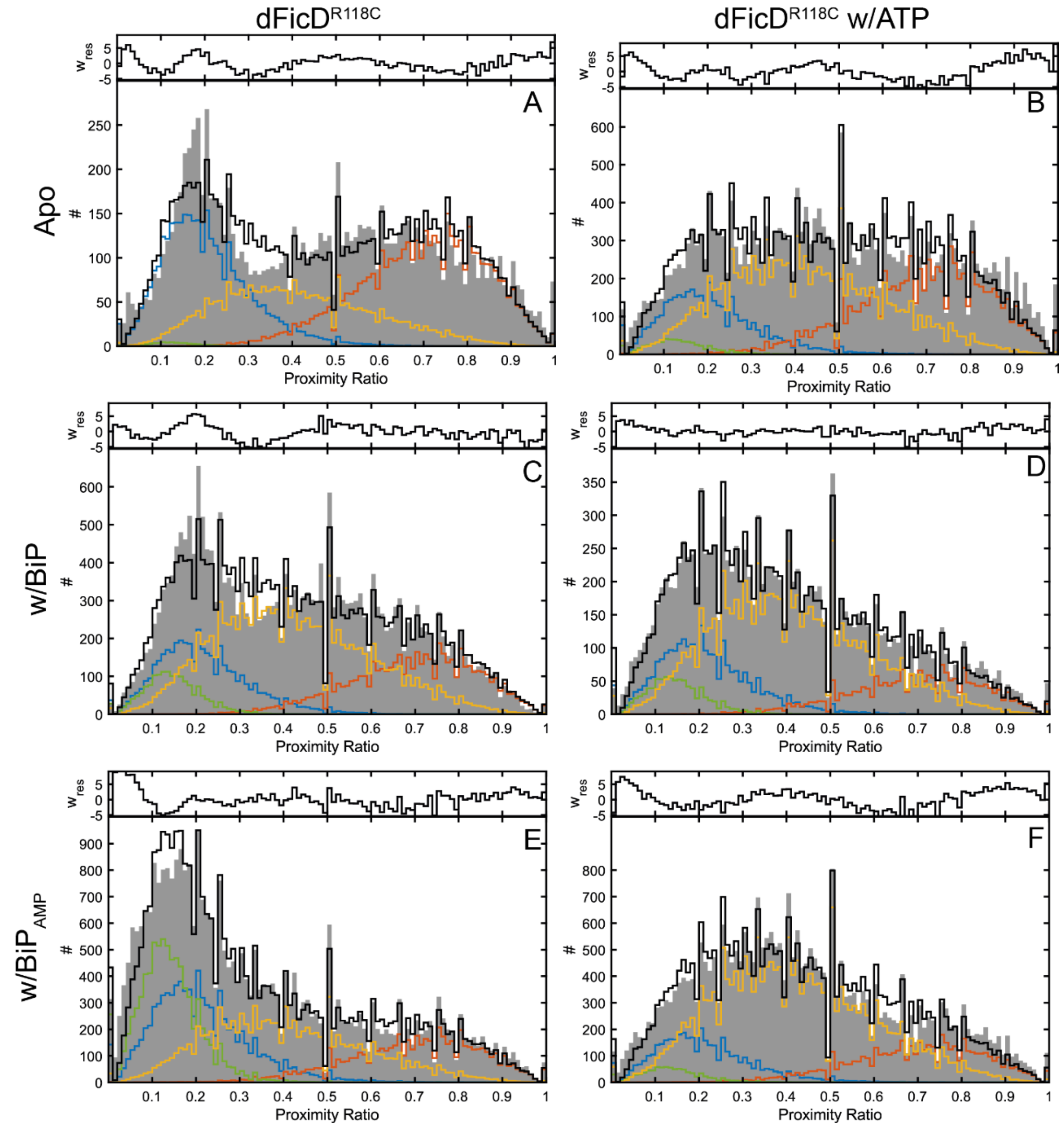

**Figure S16: Analysis of the conformational states and dynamics of dimeric FicD$^{R118C}$ monomer with photon distribution analysis (PDA).** For dynamic PDA, uncorrected FRET efficiency (proximity ratio) histograms with time binnings of 0.5 ms, 0.75 ms and 1.0 ms were used to calculate the distance distribution for different FRET populations and dynamics between them. The data for the time window size of 0.75 ms is shown for all the conditions. Proximity ratio histograms of the dimeric FicD$^{R118C}$ are given for **A:** apo, in the presence of **B:** ATP, **C:** BiP, **D:** BiP and ATP, **E:** BiP$_{AMP}$ and **F:** BiP$_{AMP}$ and ATP. The histograms for all conditions were fitted using a 2-state dynamic PDA model. TPR-in and -out conformations are given in blue and orange and the dynamic interconversion between them are shown in yellow. An additional static low-FRET (TPR-in, green) was included in the PDA fits. Values for the fits are given in **Table S16**.

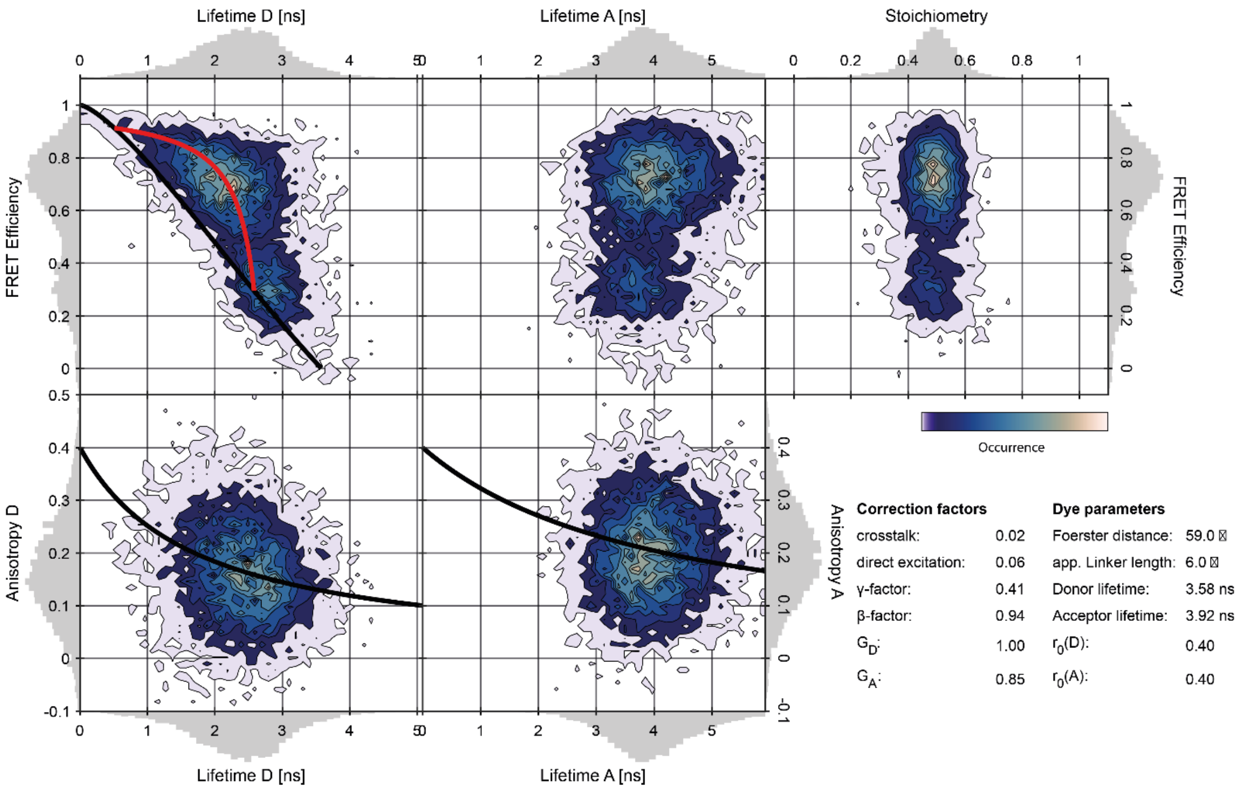

**Figure S17: A representative all-in-one plot for the two-color smFRET measurement of monomeric FicD$^{R118C}$ in the presence of ATP.** The upper left panels show the relationship between FRET efficiency and donor lifetime in the presence of acceptor (Lifetime D in ns) and acceptor lifetime (Lifetime A in ns). The upper right panel shows a stoichiometry (S) of ~0.5 (typical for double labeled molecules) and the FRET efficiency for the analyzed molecules. The lower panels depict the burst-wise anisotropy values for donor (D) and acceptor (A) fluorophores with their respective lifetimes. Black lines are fits to the Perrin equation. The applied correction factors for the FicD$^{R118C}$:ATP condition are shown in the lower right panel. The values used for all FicD FRET constructs under all measurement conditions are given in Table S14.

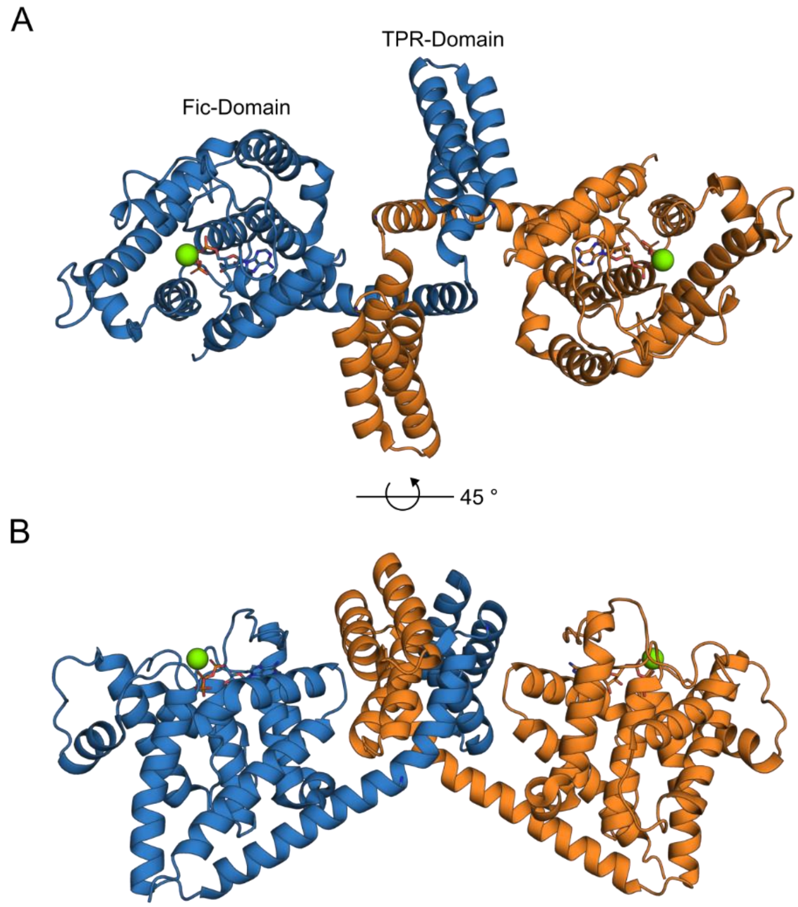

**Figure S18: Interaction of neighboring FicD molecules in the FicD:ATP crystal structure.** **A:** Top view and **B:** side view of the crystal packing in the crystal structure of FicD:ATP (PDB 6I7K). Each FicD molecule is intertwined with the TPR domain of a neighboring molecule.

**Table S1: Experimental FRET efficiency values and fractions for the monomeric FicD$^{R118C}$ construct.** $\langle E_i \rangle$ gives the mean FRET efficiency determined by a Gaussian fit to the FRET efficiency distribution histogram. The 95% confidence interval is given as the error and the $f_i$ is the fraction of the molecules at FRET efficiencies $\langle E_i \rangle$.

| FicD$^{R118C}$ | $\langle E_{static} \rangle$ | $F_{static}$ | $\langle E_{dynamic} \rangle$ | $f_{dynamic}$ |
|---|---|---|---|---|
| ***apo*** | 0.31±0.05 | 0.52 | 0.71±0.06 | 0.48 |
| ***ATP*** | 0.33±0.07 | 0.37 | 0.73±0.06 | 0.63 |
| ***BiP*** | 0.24±0.04 | 0.51 | 0.61±0.08 | 0.49 |
| ***BiP+ATP*** | 0.19±0.06 | 0.36 | 0.61±0.05 | 0.64 |
| ***BiP-AMP*** | 0.23±0.09 | 0.91 | 0.73±0.06 | 0.09 |
| ***BiP-AMP+ATP*** | 0.32±0.09 | 0.40 | 0.68±0.07 | 0.60 |

**Table S2: Distances between donor-acceptor fluorophores obtained from the accessible volume calculations for different crystal structures of FicD.** The labeling position on the Fic domain was kept constant for all the mutants (S288C). $R_{mp}$ is the mean position distance, $\langle R_{DA} \rangle$ is the average of $R_{DA}$ and $\sigma_{DA}$ is the relative error for the donor-acceptor (DA) separation calculations.

| PDB-ID | Position of dye 2 | $R_{mp}$ [Å] | $\langle R_{DA} \rangle$ [Å] | $\sigma_{DA}$ [Å] |
|---|---|---|---|---|
| ***6I7J*** | R118C | 73.7 | 73.9 | 7.8 |
| | L104C | 58.3 | 60.1 | 10.2 |
| | K154C | 59.8 | 61.5 | 7.8 |
| | S170C | 52.6 | 55.3 | 10.6 |
| | D138C | 51.1 | 54.3 | 9.8 |
| ***6I7K*** | R118C | 33.4 | 42.1 | 11.2 |
| | L104C | 56.6 | 58.6 | 12.4 |
| | K154C | 50.8 | 53.9 | 11.4 |
| | S170C | 61.8 | 61.1 | 10.5 |
| | D138C | 58.4 | 59.8 | 10.9 |
| ***6ZMD*** | R118C | 78.9 | 78.1 | 8.6 |
| | L104C | 70.2 | 69.7 | 9.3 |
| | K154C | 57.9 | 59.2 | 6.8 |
| | S170C | 61.2 | 61.8 | 9.5 |
| | D138C | 59.8 | 60.7 | 9.8 |

**Table S3: Fluorescence lifetime analysis for the monomeric FicD$^{R118C}$ construct.** For the dynamic molecules (for FRET efficiency values higher than 0.48) that appear right shifted from the static FRET lines given in the 2D E-$\tau_{D(A)}$ plots (black lines, **Figure 2** and **Figure 3**), $\tau_{out}$ and $\tau_{in}$ were determined by reconvolution fitting of the data with the instrument response function and fitting with two exponentials. The donor lifetimes $\tau_{out}$ and $\tau_{in}$ were used to determine the end points of the dynamic FRET lines (red ines, **Figure 2** and **Figure 3**) which represent the fluctuations between TPR-out and TPR-in states. For the molecules that appear static on the ms-timescales (for FRET efficiency values higher than 0.48) $\tau_{in,static}$ were determined by reconvolution fitting of the data with the instrument response function and fitting to a mono-exponential decay. The molecules that exhibit donor-lifetimes of $\tau_{in,static}$, lie on the static FRET line (black lines, **Figure 2** and **Figure 3**).The FRET efficiencies $E_{out}$, $E_{in}$ and $E_{in,static}$, and the distances $D_{out}$, $D_{in}$ and $D_{in,static}$ were calculated from the respective donor lifetimes with the lifetime of the donor-only population $\tau_{D(0)}$=3.7 ns and the Förster radius $R_0$ = 59 Å (Eq. 2*).*

| **FicD$^{R118C}$** | $\tau_{in}$ **(ns)** | **$E_{in}$** | **$D_{in}$ (Å)** | ***$f_{in}$*** | $\tau_{out}$ **(ns)** | **$E_{out}$** | **$D_{out}$ (Å)** | ***$f_{out}$*** | $\tau_{in,static}$ **(ns)** | **$E_{in,static}$** | **$D_{in,static}$ (Å)** |
|---|---|---|---|---|---|---|---|---|---|---|---|
| ***apo*** | 2.38 ±0.06 | 0.35 | 65.1 | 0.49 ±0.03 | 0.47 ±0.07 | 0.87 | 42.7 | 0.51 ±0.03 | 2.83 ±0.02 | 0.24 | 71.6 |
| ***ATP*** | 2.58 ±0.04 | 0.30 | 67.8 | 0.51 ±0.01 | 0.51 ±0.05 | 0.86 | 43.6 | 0.49 ±0.01 | 2.79 ±0.02 | 0.25 | 71.1 |
| ***BiP*** | 2.61 ±0.02 | 0.29 | 68.2 | 0.53 ±0.01 | 0.50* | 0.86 | 43.3 | 0.47 ±0.01 | 2.83 ±0.01 | 0.23 | 71.6 |
| ***BiP+ATP*** | 2.79 ±0.02 | 0.24 | 71.1 | 0.52 ±0.01 | 0.50* | 0.86 | 43.2 | 0.48 ±0.01 | 2.97 ±0.01 | 0.19 | 74.4 |
| ***BiP-AMP*** | 2.70 ±0.02 | 0.27 | 69.6 | 0.49 ±0.01 | 0.50* | 0.86 | 43.2 | 0.51 ±0.01 | 3.00 ±0.01 | 0.19 | 75.1 |
| ***BiP-AMP+ATP*** | 2.67 ±0.02 | 0.28 | 69.2 | 0.45± 0.01 | 0.50* | 0.86 | 43.2 | 0.55 ±0.01 | 2.93 ±0.01 | 0.21 | 73.6 |

***:** Values were fixed.

**Table S4: Diffusion coefficients (D in μm²/s) of the FicD$^{R118C}$ FRET construct obtained by the fluorescence correlation spectroscopy (FCS) analysis under various conditions.**

| **Monomeric FicD$^{R118C}$** | **D(μm²/s)** |
|---|---|
| ***apo*** | 61.5±0.7 |
| ***ATP*** | 58.7±0.4 |
| ***BiP*** | 50.8±0.3 |
| ***BiP+ATP*** | 41.0±0.5 |
| ***BiP-AMP*** | 36.7±0.2 |
| ***BiP-AMP+ATP*** | 47.0±0.4 |
| **Dimeric FicD$^{R118C}$** | **D(μm²/s)** |
| ***apo*** | 45.3±0.5 |
| ***ATP*** | 44.1±0.6 |
| ***BiP*** | 37.1±0.5 |
| ***BiP+ATP*** | 30.7±0.5 |
| ***BiP-AMP*** | 26.7±0.3 |
| ***BiP-AMP+ATP*** | 31.7±0.4 |

**Table S5: Fluorescence lifetime analysis for the monomeric FicD$^{L104C}$ construct.** For the dynamic molecules (for FRET efficiency values higher than 0.40) that appear right shifted from the static FRET lines given in the 2D E-$\tau_{D(A)}$ plots (black lines, **Figure S4**), $\tau_{out}$ and $\tau_{in}$ were determined by reconvolution fitting of the data with the instrument response function and fitting with two exponentials. The donor lifetimes $\tau_{out}$ and $\tau_{in}$ were used to determine the end points of the dynamic FRET lines (red ines, **Figure S4**) which represent the fluctuations between TPR-out and TPR-in states. For the molecules that appear static on the ms-timescales (for FRET efficiency values higher than 0.40) $\tau_{in,static}$ were determined by reconvolution fitting of the data with the instrument response function and fitting to a mono-exponential decay. The molecules that exhibit donor-lifetimes of $\tau_{in,static}$, lie on the static FRET line (black lines, **Figure S4**).The FRET efficiencies $E_{out}$, $E_{in}$ and $E_{in,static}$, and the distances $D_{out}$, $D_{in}$ and $D_{in,static}$ were calculated from the respective donor lifetimes with the lifetime of the donor-only population $\tau_{D(0)}$=3.7 ns and the Förster radius $R_0$ = 59 Å (Eq. 2).

| FicD$^{L104C}$ | $\tau_n$ (ns) | $E_{in}$ | $D_{in}$ (Å) | $f_{in}$ | $\tau_{out}$ (ns) | $E_{out}$ | $D_{out}$ (Å) | $f_{out}$ | $\tau_{in,static}$ (ns) | $E_{in,static}$ | $D_{in,static}$ (Å) |
|---|---|---|---|---|---|---|---|---|---|---|---|
| ***apo*** | 2.4* | 0.35 | 65.3 | 0.47 ±0.02 | 0.75 ±0.08 | 0.79 | 47.0 | 0.53 ±0.02 | 2.57 ±0.01 | 0.31 | 67.5 |
| ***ATP*** | 2.4* | 0.35 | 65.3 | 0.54 ±0.06 | 0.43 ±0.07 | 0.88 | 42.0 | 0.46 ±0.06 | 2.53 ±0.01 | 0.32 | 67.1 |
| ***BiP*** | 2.4* | 0.35 | 65.3 | 0.56 ±0.03 | 0.59 ±0.05 | 0.84 | 44.6 | 0.44 ±0.03 | 2.69 ±0.01 | 0.27 | 69.4 |
| ***BiP+ATP*** | 2.4* | 0.35 | 65.3 | 0.60 ±0.03 | 0.44 ±0.05 | 0.88 | 42.2 | 0.40 ±0.03 | 2.69 ±0.01 | 0.27 | 69.4 |
| ***BiP-AMP*** | 2.4* | 0.35 | 65.3 | 0.71 ±0.09 | 0.37 ±0.07 | 0.90 | 41.0 | 0.39 ±0.07 | 2.77 ±0.01 | 0.25 | 70.7 |
| ***BiP-AMP+ATP*** | 2.4* | 0.35 | 65.3 | 0.61 ±0.03 | 0.40 ±0.04 | 0.89 | 41.4 | 0.39 ±0.07 | 2.82 ±0.01 | 0.23 | 71.6 |

***:** Values were fixed.

**Table S6: Experimental FRET efficiency values and fractions for the FicD$^{L104C}$ construct.** $<E_i>$ gives the mean FRET efficiency determined by a Gaussian fit to the FRET efficiency distribution histogram. The 95% confidence interval is given as the error and the $f_i$ is the fraction of the molecules at FRET efficiencies $<E_i>$.

| FicD$^{L104C}$ | $<E_{static}>$ | $F_{static}$ | $<E_{dynamic}>$ | $f_{dynamic}$ |
|---|---|---|---|---|
| ***apo*** | 0.36±0.05 | 0.61 | 0.71±0.07 | 0.39 |
| ***ATP*** | 0.39±0.05 | 0.42 | 0.69±0.07 | 0.58 |
| ***BiP*** | 0.30±0.03 | 0.49 | 0.61±0.09 | 0.51 |
| ***BiP+ATP*** | 0.27±0.06 | 0.48 | 0.54±0.07 | 0.52 |
| ***BiP-AMP*** | 0.32±0.05 | 0.78 | 0.70±0.07 | 0.21 |
| ***BiP-AMP+ATP*** | 0.33±0.08 | 0.39 | 0.60±0.07 | 0.61 |

**Table S7: Fluorescence lifetime analysis for the monomeric FicD$^{K154C}$ construct.** For the measurements of the monomeric FicD$^{K154C}$ in the absence of ATP, the fluorescence lifetimes $\tau_{in}$ were determined by reconvolution fitting of the data with the instrument response function and fitting to a mono-exponential decay. The molecules that exhibit donor-lifetimes of $\tau_{in}$ on the absence of ATP lie on the static FRET line (black lines, **Figure S5**). For the measurements in the presence ATP, dynamic molecules that appear right shifted from the static FRET lines given in the 2D E-$\tau_{D(A)}$ plots (black lines, **Figure S5**), $\tau_{out}$, $\tau_{in}$ and $\tau_{in,2}$ were determined by reconvolution fitting of the data with the instrument response function and fitting with three exponentials. The donor lifetimes $\tau_{out}$ and $\tau_{in}$ were used to determine the end points of the dynamic FRET lines (red ines, **Figure S5**) which represent the fluctuations between TPR-out and TPR-in states. The FRET efficiencies $E_{out}$, $E_{in}$ and $E_{in,2}$, and the distances $D_{out}$, $D_{in}$ and $D_{in,2}$ were calculated from the respective donor lifetimes with the lifetime of the donor-only population $\tau_{D(0)}$=3.7 ns and the Förster radius $R_0$ = 59 Å (Eq. 2).

| FicD$^{K154C}$ | $\tau_{in}$ (ns) | $E_{in}$ | $D_{in}$ (Å) | $f_{in}$ | $\tau_{out}$ (ns) | $E_{out}$ | $D_{out}$ (Å) | $f_{out}$ | $\tau_{in,2}$ (ns) | $E_{in,2}$ | $D_{in,2}$ (Å) | $f_{in,2}$ |
|---|---|---|---|---|---|---|---|---|---|---|---|---|
| ***apo*** | 2.0* | 0.46 | 60.5 | - | - | - | - | - | - | - | - | - |
| ***ATP*** | 2.0* | 0.46 | 60.5 | 0.30 ±0.03 | 0.4* | 0.89 | 41.4 | 0.54 ±0.03 | 2.8* | 0.24 | 71.2 | 0.16 ±0.01 |
| ***BiP*** | 2.0* | 0.46 | 60.5 | - | - | - | - | - | - | - | - | - |
| ***BiP+ATP*** | 2.0* | 0.46 | 60.5 | 0.16 ±0.01 | 0.4* | 0.89 | 41.4 | 0.47 ±0.01 | 2.8* | 0.24 | 71.2 | 0.37 ±0.01 |
| ***BiP-AMP*** | 2.0* | 0.46 | 60.5 | - | - | - | - | - | - | - | - | - |
| ***BiP-AMP+ATP*** | 2.0* | 0.46 | 60.5 | 0.14 ±0.02 | 0.4* | 0.89 | 41.4 | 0.53 ±0.02 | 2.8* | 0.24 | 71.2 | 0.33 ±0.02 |

***:** Values were fixed.

**Table S8: Experimental FRET efficiency values and fractions for the FicD$^{K154C}$ construct.** $\langle E_i \rangle$ gives the mean FRET efficiency determined by a Gaussian fit to the FRET efficiency distribution histogram. The 95% confidence interval is given as the error and the $f_i$ is the fraction of the molecules at FRET efficiencies $\langle E_i \rangle$.

| FicD$^{K154C}$ | $\langle E_{static} \rangle$ | $F_{static}$ | $\langle E_{dynamic} \rangle$ | $f_{dynamic}$ |
|---|---|---|---|---|
| ***apo*** | 0.53±0.06 | 1 | - | - |
| ***ATP*** | 0.58±0.06 | 0.50 | 0.86±0.04 | 0.50 |
| ***BiP*** | 048±0.05 | 1 | - | - |
| ***BiP+ATP*** | 0.58±0.08 | 0.82 | 0.87±0.03 | 0.18 |
| ***BiP-AMP*** | 0.48±0.05 | 1 | - | - |
| ***BiP-AMP+ATP*** | 0.59±0.06 | 0.56 | 0.87±0.05 | 0.44 |

**Table S9: Fluorescence lifetime analysis for the monomeric FicD$^{D138C}$ construct.** For the dynamic molecules that appear right shifted from the static FRET lines given in the 2D E-$\tau_{D(A)}$ plots (black lines, **Figure S7**), $\tau_{out}$ and $\tau_{in}$ were determined by reconvolution fitting of the data with the instrument response function and fitting with two exponentials. The donor lifetimes $\tau_{out}$ and $\tau_{in}$ were used to determine the end points of the dynamic FRET lines (red ines, **Figure S7**) which represent the fluctuations between TPR-out and TPR-in states. The FRET efficiencies $E_{out}$ and $E_{in}$ , and the distances $D_{out}$, and $D_{in}$ were calculated from the respective donor lifetimes with the lifetime of the donor-only population $\tau_{D(0)}$=3.7 ns and the Förster radius $R_0$ = 59 Å (Eq. 2).

| **FicD$^{D138C}$** | **$\tau_{in}$ (ns)** | **$E_{in}$** | **$D_{in}$ (Å)** | ***$f_{in}$*** | **$\tau_{out}$ (ns)** | **$E_{out}$** | **$D_{out}$ (Å)** | ***$f_{out}$*** |
|---|---|---|---|---|---|---|---|---|
| ***apo*** | 3.14±0.06 | 0.15 | 78.6 | 0.23±0.01 | 0.91* | 0.75 | 48.9 | 0.77±0.01 |
| ***ATP*** | 2.81±0.08 | 0.24 | 71.3 | 0.33±0.02 | 0.91* | 0.75 | 48.9 | 0.67±0.02 |
| ***BiP*** | 2.89±0.04 | 0.22 | 72.9 | 0.29±0.07 | 0.91* | 0.75 | 48.9 | 0.71±0.07 |
| ***BiP+ATP*** | 3.16±0.02 | 0.15 | 79.1 | 0.53±0.07 | 0.91* | 0.75 | 48.9 | 0.47±0.07 |
| ***BiP-AMP*** | 2.99±0.03 | 0.19 | 74.9 | 0.34±0.07 | 0.91* | 0.75 | 48.9 | 0.66±0.07 |
| ***BiP-AMP+ATP*** | 3.03±0.05 | 0.18 | 75.8 | 0.51±0.02 | 0.91* | 0.75 | 48.9 | 0.49±0.02 |

***:** Values were fixed.

**Table S10: Fluorescence lifetime analysis for the monomeric FicD$^{S170C}$ construct.** For the dynamic molecules that appear right shifted from the static FRET lines given in the 2D E-$\tau_{D(A)}$ plots (black lines, **Figure S8**), $\tau_{out}$ and $\tau_{in}$ were determined by reconvolution fitting of the data with the instrument response function and fitting with two exponentials. The donor lifetimes $\tau_{out}$ and $\tau_{in}$ were used to determine the end points of the dynamic FRET lines (red ines, **Figure S8**) which represent the fluctuations between TPR-out and TPR-in states. The FRET efficiencies $E_{out}$ and $E_{in}$ , and the distances $D_{out}$, and $D_{in}$ were calculated from the respective donor lifetimes with the lifetime of the donor-only population $\tau_{D(0)}$=3.7 ns and the Förster radius $R_0$ = 59 Å (Eq. 2).

| **FicD$^{S170C}$** | **$\tau_{in}$ (ns)** | **$E_{in}$** | **$D_{in}$ (Å)** | ***$f_{in}$*** | **$\tau_{out}$ (ns)** | **$E_{out}$** | **$D_{out}$ (Å)** | ***$f_{out}$*** |
|---|---|---|---|---|---|---|---|---|
| ***apo*** | 2.40±0.05 | 0.35 | 65.3 | 0.50±0.02 | 0.91* | 0.76 | 48.9 | 0.50±0.02 |
| ***ATP*** | 2.62±0.06 | 0.29 | 68.3 | 0.42±0.02 | 0.91* | 0.76 | 48.9 | 0.58±0.02 |
| ***BiP*** | 2.54±0.02 | 0.31 | 67.2 | 0.54±0.01 | 0.91* | 0.76 | 48.9 | 0.46±0.01 |
| ***BiP+ATP*** | 2.67±0.05 | 0.28 | 69.1 | 0.49±0.02 | 0.91* | 0.76 | 48.9 | 0.51±0.02 |
| ***BiP-AMP*** | 2.62±0.02 | 0.29 | 68.3 | 0.58±0.01 | 0.91* | 0.76 | 48.9 | 0.42±0.01 |
| ***BiP-AMP+ATP*** | 2.85±0.03 | 0.23 | 72.1 | 0.49±0.01 | 0.91* | 0.76 | 48.9 | 0.54±0.01 |

***:** Values were fixed.

**Table S11: Results of the global dynamic PDA of the monomeric FicD$^{R118C}$ construct.** For dynamic PDA, uncorrected FRET efficiency (proximity ratio) histograms with time binnings of 0.5 ms, 0.75 ms and 1.0 ms were used to calculate the distance distribution for different FRET populations and the dynamics between them. Global fits for the experimental conditions listed below were carried out on the datasets with 0.75 ms bin time. Error bars for the kinetic rates $k_{in}$ and $k_{out}$ are given as 95% confidence intervals as determined from the curvature of the $\chi^2_{red}$-surface.

| FicD$^{R118C}$ | $R_{out}$ (Å) | $\sigma_{out}$ (Å) | $k_{in}$ ($ms^{-1}$) | $R_{in}$ (Å) | $\sigma_{in}$ (Å) | $k_{out}$ ($ms^{-1}$) | $A_{in, static}$ | $R_{in, static}$ (Å) | $\sigma_{in, static}$ (Å) | $A_{R1}$ | $R_{R1}$ (Å) | $\sigma_{R1}$ (Å) | $\chi^2_{red}$ |
|---|---|---|---|---|---|---|---|---|---|---|---|---|---|
| ***apo*** | 43 | 4.5 | 1.08 ±0.05 | 65 | 1 | 0.88 ±0.05 | 0.25 | 71 | 2.5 | 0.01 | 92 | 3.5 | 4.0 |
| ***ATP*** | 43 | 4.5 | 1.78 ±0.05 | 65 | 4.5 | 2.57 ±0.05 | 0.15 | 71 | 2.5 | 0.02 | 92 | 3.5 | 4.7 |
| ***BiP*** | 43 | 4.5 | 2.32 ±0.05 | 65 | 1 | 1.50 ±0.05 | 0.45 | 71 | 2.5 | 0 | 92 | 3.5 | 12.2 |
| ***BiP+ATP*** | 43 | 4.5 | 2.84 ±0.05 | 65 | 4.5 | 2.32 ±0.05 | 0.31 | 71 | 2.5 | 0.02 | 92 | 3.5 | 2.4 |
| ***BiP-AMP*** | 43 | 4.5 | 2.00 ±0.05 | 65 | 1 | 0.75 ±0.05 | 0.86 | 71 | 2.5 | 0.21 | 92 | 3.5 | 15.6 |
| ***BiP-AMP +ATP*** | 43 | 4.5 | 1.99 ±0.05 | 65 | 4.5 | 2.50 ±0.05 | 0.13 | 71 | 2.5 | 0.04 | 92 | 3.5 | 8.36 |

**Table S12: Results of the global dynamic PDA of the monomeric FicD$^{L104C}$ construct. .** For dynamic PDA, uncorrected FRET efficiency (proximity ratio) histograms with time binnings of 0.5 ms, 0.75 ms and 1.0 ms were used to calculate the distance distribution for different FRET populations and the dynamics between them. Global fits for the experimental conditions listed below were carried out on the datasets with 0.75 ms bin time. Error bars for the kinetic rates $k_{in}$ and $k_{out}$ are given as 95% confidence intervals as determined from the curvature of the $\chi^2_{red}$-surface.

| FicD$^{L104C}$ | $R_{out}$ (Å) | $\sigma_{out}$ (Å) | $k_{in}$ (ms$^{-1}$) | $R_{in}$ (Å) | $\sigma_{in}$ (Å) | $k_{out}$ (ms$^{-1}$) | $A_{in, static}$ | $R_{in, static}$ (Å) | $\sigma_{in, static}$ (Å) | $A_{R1}$ | $R_{R1}$ (Å) | $\sigma_{d1}$ (Å) | $\chi^2_{red}$ |
|---|---|---|---|---|---|---|---|---|---|---|---|---|---|
| ***apo*** | 43 | 4 | 1.43 ±0.11 | 60 | 3.5 | 1.12 ±0.14 | 0.93 | 65 | 3 | 0.03 | 95 | 3 | 1.4 |
| ***ATP*** | 43 | 4 | 1.65 ±0.09 | 60 | 3.5 | 1.22 ±0.11 | 0.27 | 65 | 3 | 0.04 | 95 | 3 | 3.7 |
| ***BiP*** | 43 | 4 | 1.81 ±0.07 | 60 | 3.5 | 1.15 ±0.06 | 0.87 | 65 | 3 | 0.03 | 95 | 3 | 5.0 |
| ***BiP+ATP*** | 43 | 4 | 2.88 ±0.12 | 60 | 3.5 | 0.68 ±0.04 | 0.66 | 69 | 3 | 0.05 | 95 | 3 | 3.9 |
| ***BiP-AMP*** | 43 | 4 | 0.89 ±0.06 | 60 | 3.5 | 0.14 ±0.01 | 0.70 | 69 | 3 | 0.11 | 95 | 3 | 12.2 |
| ***BiP-AMP+ATP*** | 43 | 4 | 2.47 ±0.08 | 60 | 3.5 | 1.41 ±0.06 | 0.15 | 69 | 3 | 0.02 | 95 | 3 | 6.5 |

**Table S13: Results of the global dynamic PDA of the monomeric FicD$^{K154C}$ construct.** For both static and dynamic PDA, uncorrected FRET efficiency (proximity ratio) histograms with time binnings of 0.5 ms, 0.75 ms and 1.0 ms were used to calculate the distance distribution for different FRET populations and the dynamics between them. A 3-state static PDA model was used to fit the proximity ratio histograms of FicD$^{K154C}$ in the absence of ATP. A 2-state dynamic PDA model was used to fit the proximity ratio histograms of FicD$^{K154C}$ in the presence of of ATP. Global fits for the experimental conditions listed below were carried out on the datasets with 0.75 ms bin time. Error bars for the kinetic rates $k_{in}$ and $k_{out}$ are given as 95% confidence intervals as determined from the curvature of the $\chi^2_{red}$-surface.

| **FicD$^{K154C}$ static** | **$R_{out}$ (Å)** | **$\sigma_{out}$ (Å)** | **$A_{out}$** | **$R_{in}$ (Å)** | **$\sigma_{in}$ (Å)** | **$A_{in}$** | **$R_{in,2}$ (Å)** | **$\sigma_{in,2}$ (Å)** | **$A_{in,2}$** | **$\chi^2_{red}$** |
|---|---|---|---|---|---|---|---|---|---|---|
| ***apo*** | 41 | 3.5 | 0.07 | 56 | 3.2 | 0.84 | 74 | 3.0 | 0.07 | 5.9 |
| ***BiP*** | 41 | 3.5 | 0.04 | 57 | 3.6 | 0.89 | 74 | 3.0 | 0.05 | 11.1 |
| ***BiP-AMP*** | 41 | 3.5 | 0.04 | 57 | 3.1 | 0.86 | 74 | 3.0 | 0.10 | 6.8 |
| **FicD$^{K154C}$ 2 state dynamic** | **$R_{out}$ (Å)** | **$\sigma_{out}$ (Å)** | **$k_{in}$ ($ms^{-1}$)** | **$R_{in}$ (Å)** | **$\sigma_{in}$ (Å)** | **$k_{out}$ ( $ms^{-1}$)** | **$R_{in, static}$ (Å)** | **$\sigma_{in, static}$ (Å)** | **$A_{in, static}$** | **$\chi^2_{red}$** |
| ***BiP+ATP*** | 41 | 4.5 | 0.66 ±0.05 | 57 | 3 | 0.80 ±0.06 | 74 | 3.0 | 0.03 | 3.5 |
| ***BiP+ATP*** | 41 | 4.5 | 0.92 ±0.04 | 57 | 3 | 0.48 ±0.02 | 74 | 3.0 | 0.09 | 4.8 |
| ***BiP-AMP+ATP*** | 41 | 4.5 | 0.69 ±0.03 | 57 | 3 | 0.73 ±0.04 | 74 | 3.0 | 0.06 | 4.0 |

**Table S14: Fluorescence lifetime analysis for the dimeric FicD$^{R118C}$ construct.** For the dynamic molecules (for FRET efficiency values higher than 0.48) that appear right shifted from the static FRET lines given in the 2D E-$\tau_{D(A)}$ plots (black lines, **Figure S13**), $\tau_{out}$ and $\tau_{in}$ were determined by reconvolution fitting of the data with the instrument response function and fitting with two exponentials. The donor lifetimes $\tau_{out}$ and $\tau_{in}$ were used to determine the end points of the dynamic FRET lines (red ines, **Figure S13**) which represent the fluctuations between TPR-out and TPR-in states. For the molecules that appear static on the ms-timescales (for FRET efficiency values higher than 0.48) $\tau_{in,static}$ were determined by reconvolution fitting of the data with the instrument response function and fitting to a mono-exponential decay. The molecules that exhibit donor-lifetimes of $\tau_{in,static}$, lie on the static FRET line (black lines, **Figure S13**).The FRET efficiencies $E_{out}$, $E_{in}$ and $E_{in,static}$, and the distances $D_{out}$, $D_{in}$ and $D_{in,static}$ were calculated from the respective donor lifetimes with the lifetime of the donor-only population $\tau_{D(0)}$=3.7 ns and the Förster radius $R_0$ = 59 Å (Eq. 2).

| **dFicD$^{R118C}$** | **$\tau_{in}$ (ns)** | **$E_{in}$** | **$D_{in}$ (Å)** | ***$f_{in}$*** | **$\tau_{out}$ (ns)** | **$E_{out}$** | **$D_{out}$ (Å)** | ***$f_{out}$*** | **$\tau_{in,static}$ (ns)** | **$E_{in,static}$** | **$D_{in,static}$ (Å)** |
|---|---|---|---|---|---|---|---|---|---|---|---|
| ***apo*** | 2.45 ±0.07 | 0.34 | 65.9 | 0.33 ±0.02 | 0.65 ±0.05 | 0.82 | 45.6 | 0.67 ±0.02 | 2.75 ±0.02 | 0.26 | 70.3 |
| ***ATP*** | 2.75 ±0.07 | 0.26 | 70.4 | 0.37 ±0.02 | 0.75 ±0.05 | 0.79 | 47.0 | 0.63 ±0.02 | 2.82 ±0.02 | 0.24 | 71.5 |
| ***BiP*** | 2.60 ±0.09 | 0.30 | 68.1 | 0.37 ±0.01 | 0.60 ±0.01 | 0.84 | 44.9 | 0.63 ±0.01 | 2.80 ±0.05 | 0.24 | 71.2 |
| ***BiP+ATP*** | 2.63± 0.08 | 0.29 | 68.4 | 0.41 ±0.03 | 0.70 ±0.08 | 0.81 | 46.3 | 0.59 ±0.03 | 2.802 ±0.05 | 0.24 | 71.2 |
| ***BiP-AMP*** | 2.66 ±0.08 | 0.28 | 68.9 | 0.41 ±0.02 | 0.71 ±0.06 | 0.80 | 46.4 | 0.59 ±0.02 | 2.93± 0.03 | 0.21 | 73.6 |
| ***BiP-AMP+ATP*** | 2.77 ±0.07 | 0.25 | 70.7 | 0.46 ±0.02 | 0.84 ±0.06 | 0.77 | 48.1 | 0.54 ±0.02 | 2.75± 0.02 | 0.26 | 70.3 |
| **Inactive dFicD$^{R118C}$** | **$\tau_{in}$ (ns)** | **$E_{in}$** | **$D_{in}$ (Å)** | ***$f_{in}$*** | **$\tau_{out}$ (ns)** | **$E_{out}$** | **$D_{out}$ (Å)** | ***$f_{out}$*** | **$\tau_{in,static}$ (ns)** | **$E_{in,static}$** | **$D_{in,static}$ (Å)** |
| ***BiP-AMP*** | 2.63 ±0.10 | 0.29 | 68.5 | 0.52 ±0.05 | 0.93 ±0.10 | 0.75 | 49.2 | 0.58 ±0.05 | 2.65 ±0.02 | 0.28 | 68.8 |
| ***BiP-AMP+ATP*** | - | - | - | - | - | - | - | - | 2.69 ±0.02 | 0.28 | 69.3 |

***:** Values were fixed.

**Table S15: Experimental FRET efficiency values and fractions for the dimeric FicD$^{R118C}$ construct.** $\langle E_i \rangle$ gives the mean FRET efficiency determined by a Gaussian fit to the FRET efficiency distribution histogram. The 95% confidence interval is given as the error and the $f_i$ is the fraction of the molecules at FRET efficiencies $\langle E_i \rangle$.

| **dFicD$^{R118C}$** | **$\langle E_{static} \rangle$** | **$F_{static}$** | **$\langle E_{dynamic} \rangle$** | **$f_{dynamic}$** | **$\langle E_{dynamic,2} \rangle$** | **$F_{dynamic,2}$** |
|---|---|---|---|---|---|---|
| ***apo*** | 0.28±0.04 | 0.33 | 0.75±0.07 | 0.67 | - | - |
| ***ATP*** | 0.33±0.06 | 0.24 | 0.70±0.06 | 0.54 | 0.91±0.03 | 0.21 |
| ***BiP*** | 0.25±0.05 | 0.35 | 0.72±0.07 | 0.65 | - | - |
| ***BiP+ATP*** | 0.21±0.05 | 0.27 | 0.52±0.08 | 0.60 | 0.86±0.03 | 0.13 |
| ***BiP-AMP*** | 0.26±0.08 | 0.59 | 0.66±0.06 | 0.23 | 0.89±0.03 | 0.16 |
| ***BiP-AMP+ATP*** | - | - | 0.56±0.10 | 1 | - | - |
| **Inactive dFicD$^{R118C}$** | **$\langle E_{static} \rangle$** | **$f_{static}$** | **$\langle E_{dynamic} \rangle$** | **$f_{dynamic}$** | **$\langle E_{dynamic,2} \rangle$** | **$f_{dynamic,2}$** |
| ***BiP-AMP*** | 0.25±0.05 | 0.37 | 0.61±0.08 | 0.63 | - | - |
| ***BiP-AMP+ATP*** | 0.28±0.06 | 1 | - | - | - | - |

**Table S16: Diffusion coefficients (D in μm$^2$/s) of the FicD$^{R118C}$ FRET constructs at different FRET states obtained by the fluorescence correlation spectroscopy (FCS) analysis under various conditions.** Results of the FCS analysis on the single-molecule bursts that correspond to the static TPR-in state (E<0.4) and the dynamic population fluctuating between the TPR-in and TPR-out conformation (E>0.4).

| **Monomeric FicD$^{R118C}$** | **D(μm$^2$/s) of the static TPR-in state (E<0.4)** | **D(μm$^2$/s) of the dynamic high-FRET population (E>0.4)** |
|---|---|---|
| **BiP** | 52.6±1.0 | 54.0±1.1 |
| **BiP+ATP** | 38.3±0.7 | 40.8±0.6 |
| **BiP$_{AMP}$** | 31.9±0.9 | 34.1±0.5 |
| **BiP$_{AMP}$ +ATP** | 44.6±1.2 | 35.8±1.8 |
| **Dimeric FicD$^{R118C}$** | **D(μm$^2$/s) of the static TPR-in state (E<0.4)** | **D(μm$^2$/s) of the dynamic high-FRET population (E>0.4)** |
| **BiP** | 43.0±1.1 | 39.2±1.3 |
| **BiP+ATP** | 31.9±0.5 | 31.2±1.5 |
| **BiP$_{AMP}$** | 25.1±0.4 | 34.0±1.6 |
| **BiP$_{AMP}$ +ATP** | 35.5±0.8 | 32.8±1.8 |

**Table S17: BiP$_{AMP}$ bound and unbound fractions of monomeric and dimeric FicD$^{R118C}$ revealed by the fluorescence correlation spectroscopy (FCS).** The diffusion coefficient values were obtained from previous FCS fits for the unbound (FicD:apo, **Table S4**) and bound states (FicD:BiP$_{AMP}$ for the static TPR-in state (E<0.4), **Table S16**). A 2-component fitting was performed to reveal the fraction of the bound and unbound species in the different FRET states.

| **Monomeric FicD$^{R118C}$** | **D(μm$^2$/s) unbound** | **$N_{unbound}$** | **D(μm$^2$/s) bound** | **$N_{bound}$** | **$f_{bound}$** |
|---|---|---|---|---|---|
| **BiP$_{AMP}$, static TPR-in state (E<0.4)** | 61 | 0 | 32 | 0.1 | 100% |
| **BiP$_{AMP}$, high-FRET population (E>0.4)** | 61 | 0.01 | 32 | 0.09 | 90% |
| **BiP$_{AMP}$ +ATP, static TPR-in state (E<0.4)** | 61 | 0.04 | 32 | 0.04 | 50% |
| **BiP$_{AMP}$ +ATP, high-FRET population (E>0.4)** | 61 | 0.01 | 32 | 0.06 | 83% |
| **Dimeric FicD$^{R118C}$** | **D(μm$^2$/s) unbound** | **$N_{unbound}$** | **D(μm$^2$/s) bound** | **$N_{bound}$** | **$f_{bound}$** |
| **BiP$_{AMP}$, static TPR-in state (E<0.4)** | 45 | 0 | 25 | 0.06 | 100% |
| **BiP$_{AMP}$, high-FRET population (E>0.4)** | 45 | 0.02 | 25 | 0.02 | 50% |
| **BiP$_{AMP}$ +ATP, static TPR-in state (E<0.4)** | 45 | 0.03 | 25 | 0.02 | 40% |
| **BiP$_{AMP}$ +ATP, high-FRET population (E>0.4)** | 45 | 0.02 | 25 | 0.02 | 50% |

**Table S18: Results of the global dynamic PDA of the dimeric FicD$^{R118C}$ construct.** For dynamic PDA, uncorrected FRET efficiency (proximity ratio) histograms with time binnings of 0.5 ms, 0.75 ms and 1.0 ms were used to calculate the distance distribution for different FRET populations and the dynamics between them. Global fits for the experimental conditions listed below were carried out on the datasets with 0.75 ms bin time. Error bars for the kinetic rates $k_{in}$ and $k_{out}$ are given as 95% confidence intervals as determined from the curvature of the $\chi^2_{red}$-surface.

| dFicD$^{R118C}$ | $R_{out}$ (Å) | $\sigma_{out}$ (Å) | $k_{in}$ ($ms^{-1}$) | $R_{in}$ (Å) | $\sigma_{in}$ (Å) | $k_{out}$ ( $ms^{-1}$) | $A_{in, static}$ | $R_{in, static}$ (Å) | $\sigma_{in, static}$ (Å) | $\chi^2_{red}$ |
|---|---|---|---|---|---|---|---|---|---|---|
| ***apo*** | 43 | 5 | 0.29 ±0.03 | 65 | 6 | 0.37 ±0.05 | 0.01 | 71 | 3.5 | 6.2 |
| ***ATP*** | 43 | 5 | 0.55 ±0.03 | 65 | 6 | 0.86 ±0.06 | 0.03 | 71 | 3.5 | 9.4 |
| ***BiP*** | 43 | 5 | 0.76 ±0.04 | 65 | 6 | 0.87 ±0.07 | 0.07 | 71 | 3.5 | 7.3 |
| ***BiP+ATP*** | 43 | 5 | 1.04 ±0.06 | 65 | 6 | 0.95 ±0.09 | 0.07 | 71 | 3.5 | 3.3 |
| ***BiP-AMP*** | 43 | 5 | 0.62 ±0.03 | 65 | 6 | 0.48 ±0.04 | 0.32 | 71 | 3.5 | 9.8 |
| ***BiP-AMP +ATP*** | 43 | 5 | 1.10 ±0.04 | 65 | 6 | 1.21 ±0.07 | 0.03 | 71 | 3.5 | 9.4 |

**Table S19: Correction factors used for the smFRET measurements of FicD FRET constructs.** In the table, γ is the detection correction factor, β is the normalization of direct donor and acceptor excitation fluxes, α is the spectral crosstalk of donor fluorescence into the acceptor channel and δ is the ratio of indirect and direct acceptor excitation.

| | **FicD$^{R118C}$** | | | | | | **FicD$^{L104C}$** | | | | | |
|---|---|---|---|---|---|---|---|---|---|---|---|---|
| ***ATP*** | | + | | + | | + | | + | | + | | + |
| ***BiP*** | | | + | + | | | | | + | + | | |
| ***BiP$_{AMP}$*** | | | | | + | + | | | | | + | + |
| ***γ*** | 0.41 | 0.41 | 0.51 | 0.44 | 0.41 | 0.43 | 0.46 | 0.37 | 0.54 | 0.56 | 0.41 | 0.47 |
| ***β*** | 0.94 | 0.94 | 0.91 | 0.94 | 1.01 | 1.00 | 1.00 | 1.09 | 0.87 | 0.94 | 1.07 | 0.98 |
| ***α*** | 0.02 | 0.02 | 0.03 | 0.03 | 0.03 | 0.03 | 0.02 | 0.02 | 0.02 | 0.03 | 0.02 | 0.02 |
| ***δ*** | 0.07 | 0.07 | 0.06 | 0.10 | 0.10 | 0.08 | 0.05 | 0.06 | 0.07 | 0.03 | 0.08 | 0.11 |
| | **FicD$^{K154C}$** | | | | | | **FicD$^{S170C}$** | | | | | |
| ***ATP*** | | + | | + | | + | | + | | + | | + |
| ***BiP*** | | | + | + | | | | | + | + | | |
| ***BiP$_{AMP}$*** | | | | | + | + | | | | | + | + |
| ***γ*** | 0.44 | 0.40 | 0.44 | 0.40 | 0.44 | 0.44 | 0.47 | 0.47 | 0.47 | 0.47 | 0.47 | 0.47 |
| ***β*** | 1.11 | 1.11 | 1.11 | 1.11 | 1.11 | 1.11 | 1.11 | 1.11 | 1.11 | 1.11 | 1.11 | 1.11 |
| ***α*** | 0.02 | 0.02 | 0.04 | 0.03 | 0.02 | 0.03 | 0.02 | 0.02 | 0.03 | 0.03 | 0.02 | 0.03 |
| ***δ*** | 0.04 | 0.07 | 0.09 | 1.00 | 0.09 | 0.08 | 0.05 | 0.07 | 0.09 | 0.09 | 0.08 | 0.09 |
| | **FicD$^{D138C}$** | | | | | | **dFicD$^{R118C}$** | | | | | |
| ***ATP*** | | + | | + | | + | | + | | + | | + |
| ***BiP*** | | | + | + | | | | | + | + | | |
| ***BiP$_{AMP}$*** | | | | | + | + | | | | | + | + |
| ***γ*** | 0.42 | 0.42 | 0.42 | 0.41 | 0.41 | 0.41 | 0.48 | 0.38 | 0.38 | 0.52 | 0.41 | 0.45 |
| ***β*** | 1.15 | 1.15 | 1.15 | 1.16 | 1.11 | 1.11 | 1.22 | 1.22 | 1.22 | 0.94 | 1.07 | 0.96 |
| ***α*** | 0.02 | 0.02 | 0.04 | 0.04 | 0.03 | 0.02 | 0.02 | 0.02 | 0.04 | 0.03 | 0.02 | 0.02 |
| ***δ*** | 0.09 | 0.06 | 0.15 | 0.12 | 0.09 | 0.13 | 0.04 | 0.05 | 0.12 | 0.08 | 0.08 | 0.09 |
